# RawMod: Modification-Agnostic Detection of DNA Base Modifications from Nanopore Signal Pileups

**DOI:** 10.64898/2026.11.10.765445

**Authors:** Bhargav Srinivasan, Sabeena Nazar, Vir Gandhi, Ernest Zhang, Tsvetan Bachvaroff, Mikhail Kolmogorov, Allen Place, Can Firtina

**Author notes:** Corresponding authors {, }.

## Abstract

Identifying epigenetically modified bases (e.g., methylation) is crucial for many important applications such as discovering mechanisms of cancer development, cell differentiation, and bacterial defense. Nanopore sequencing generates electrical signals that can differentiate canonical bases from modified ones. To detect base modifications from signals, machine learning (ML) methods and statistical approaches are commonly used. These methods can either detect a specific modification type or require an additional unmodified copy of the same DNA as a control group. However, no released model detects modification types beyond those it was trained for, which makes it difficult to detect rare modification types without an unmodified control. Our goal is to accurately identify the presence of modified sites regardless of their modification type, which we define as modification-agnostic detection. To this end, we propose RawMod, a transformer-based modification-agnostic and control-free method for detecting epigenetically modified genomic positions. The key idea of RawMod is to compare the raw signals of many reads at each site with the unmodified reference sequence, using a small model that learns many modification types as a single modified class. Based on our extensive evaluations on the detection of various modification types (5mC, 5hmC, 4mC, 6mA, and 5hmU) using various genomes, RawMod achieves the highest mean AUROC across modifications compared to control-based methods and control-free modification models. A single RawMod model with 0.23M parameters detects five modification types with a mean AUROC of 0.964, against 0.799 for the best released model. When a modification type is withheld from training, RawMod still detects it (AUROC 0.91–0.98), provided another modification of the same base was seen in training.

We use RawMod to map 5hmU without a control across a 7.7 Mb contig of the Amphidinium carterae genome, consistent with two published trends. Further, we show that the modification-aware embeddings from RawMod can be used to differentiate read origin (e.g., cell types, tumor/normal origin, and native/PCR-amplified DNA). We discuss how RawMod can enable further work on label-free classification of cell types using these modification-aware embeddings.

**Availability and Implementation:** RawMod is available at https://github.com/STORM_group/RawMod and at https://huggingface.co/STORMResearch/RawMod. We also provide the scripts to fully reproduce our results on our GitHub page and release our datasets on SquiDBase.

## 1. Introduction

Nanopore sequencing technology [1–23] reads single DNA molecules that can be millions of bases long [24, 25]. Each molecule is sequenced by moving through a tiny pore, called a *nanopore*, disrupting the ionic current flow inside the nanopore as it moves. These disruptions are measured as electrical signals (i.e., *raw nanopore signals*) at a certain sampling frequency (e.g., 5000 signals per second). Raw nanopore signals reflect the bases of the nucleic acid molecule and their chemical alter-ations, called *base modifications* as shown in Figure 1. Machine learning models can decode these signals into DNA sequences, or *reads*, a process known as *basecalling* [26–43], and identify base modifications such as 5-methylcytosine (5mC) [44–61].

**Figure 1.**
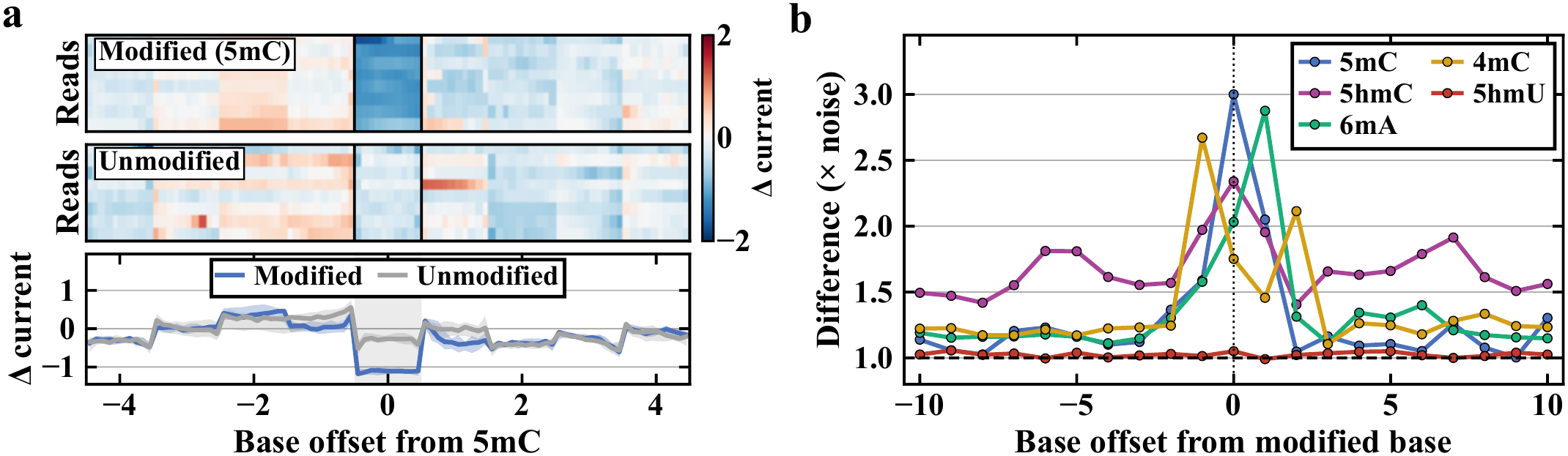
Modifications perturb raw signal depending on their type. (a) Pileups of one position in a 5mC-modified library and its unmodified control. (b) Library difference relative to the noise floor.

Base modifications let DNA carry information beyond its sequence [62]. Mammalian cells use 5mC and 5-hydroxymethylcytosine (5hmC) to regulate transcription and cell identity, while bacterial cells use N6-methyladenine (6mA) and N4-methylcytosine (4mC) for host defense, replication and virulence [63]. A few organisms utilize a *non-standard base*, a base outside the expected A, C, G and T, across much of their genome. For example, the bacteriophage SPO1 carries 5-hydroxymethyluracil (5hmU) in place of every thymine [64,65], and dinoflagellates such as *Amphidinium carterae* carry it in place of a large share of their thymines [66, 67]. In kineto-plastid parasites, base J, a glucosylated 5hmU, marks the ends of gene clusters, where it stops transcription [68–70]. Identifying these modifications and non-standard bases from raw nanopore signals enables further analysis of their downstream impacts. For example, cancer cells commonly show abnormal methylation across the genome [71, 72], which nanopore sequencing can map directly. Such analyses require methods that identify diverse modification types accurately and efficiently.

To identify these base modifications accurately, existing methods find modifications from raw signals mainly in two ways. First, *supervised callers* [31, 44, 45, 47–50, 54, 56, 58, 60, 61, 73], such as Dorado [31], Rockfish [48], DeepMod2 [58], and UniMeth [61] use machine learning (ML) models to classify certain modification types per read in specific sequence contexts. These callers are not *modification-agnostic*: each detects only the modification types it is trained to detect. Supporting a new modification type requires a carefully curated training dataset labeled for that modification and a newly trained model, which limits the identification of rare modification types such as 5hmU, for which no released caller exists. Building a modification-agnostic model is challenging because the model must separate signal perturbations caused by modifications from those caused by the sequence context, the organism, and the library preparation through multiple modification types. Since these factors often co-vary with modification presence labels in training data, a model can learn to falsely recognize these patterns in the signal. Second, *comparative callers* [46, 74–77], such as Tombo [74], nanodisco [75], and Uncalled4 [76], flag reference positions where a sample’s signal deviates from a baseline of unmodified signal, which makes them modification-agnostic by design. This baseline typically comes from sequencing a matched control in which amplification by Polymerase Chain Reaction (PCR) [46, 76] or whole-genome amplification (WGA) [75] has removed all modifications. Preparing and sequencing such a control for every sample increases cost and turnaround time, and is difficult for unculturable organisms or clinical tissue. Learning a model of unmodified signal instead of sequencing a control [77] removes this requirement, but is still limited by confounding variation between modifications and sample-specific factors. To our knowledge, no model can accurately identify modified DNA sites regardless of their modification type from the raw nanopore signals of a sample without an unmodified control.

Our goal is to accurately identify modified genomic positions, called *sites*, regardless of their modification type and without an unmodified control. To this end, we propose RawMod, a modification-agnostic model that can identify the presence of *any* DNA base modification in four key steps. First, to capture modification evidence, it builds a pileup of raw signals around each queried site. Second, to integrate information across reads, a convolutional encoder and transformer aggregate the signals into a site-level embedding. Third, to separate modified from unmodified sites in the embedding space, RawMod uses classification, supervised contrastive learning, and anomaly detection objectives. Fourth, to learn the modification signal before sequence shortcuts, a curriculum trains first on pairwise labels, whose modified and unmodified sites share the same sequence, and then on the full dataset. Thus, the earliest gradients come only from the modification contrast.

Across bacterial, phage, plant, mouse, human, and synthetic data covering five modification types, our *single* RawMod model detects the presence of 5mC, 5hmC, 6mA, 4mC and 5hmU. RawMod provides the highest mean area under the receiver operating characteristic curve (AUROC) of 0.964 across the five types compared with 0.799 for the best released model, Dorado 4mC+5mC. It does so with only 0.23M parameters, 5–435× fewer than other callers (1.24M–100.3M). When each modification type is withheld from training, our model still detects three of the five types (AUROC 0.91–0.98). Transfer occurs within a base, as all three are cytosine modifications. 6mA and 5hmU, the only types on their base in our data, do not transfer (0.59 and 0.50). Retraining with about 3,000 labeled 5hmU sites raises 5hmU to an AUROC of 0.82 (Section 3.4). To support methods for such rare modifications, we sequence and release raw nanopore signal of SPO1 and *A. carterae*, which carry 5hmU. We make the following key contributions:

### Key Contributions

- RawMod, a single model that detects the presence of base modifications of any type with mean AUROC 0.964 over the five modification types, higher than any other tested method. The five types include 5hmU, which no released caller can identify.
- We sequence and release raw nanopore signal of SPO1 and *Amphidinium carterae*, including the first public nanopore dataset of 5hmU DNA with an unmodified control (native and PCR SPO1).
- A control-free 5hmU map of an *Amphidinium carterae* contig that is consistent with two published trends and provides a testable hypothesis for 5hmU placement.
- Applications of modification-agnostic embeddings that separate reads with similar sequence in tumor/normal, cell, and native/PCR typing experiments.
- Open source code, model weights, and scripts to fully reproduce our results, use our tools easily, and train models on new modification types and datasets at https://github.com/STORMgroup/RawMod.

## 2. Methods

The goal of RawMod is to identify if a base modification is present at a reference position on one strand of a genome, called a *site*, regardless of the modification type, which we call *modification-agnostic* detection. To achieve this, for a site *i* with base *b*_*i*_, RawMod predicts the likelihood *p*_*i*_ ∈ [0, 1] that the site is modified, where every modification contributes to the *single* positive class *y*_*i*_=1 and an unmodified site has *y*_*i*_=0.

Figure 2 shows three key components of RawMod, which takes the raw signals of basecalled reads aligned to a reference genome and generates the per-site modification likelihood *p*_*i*_ that indicates whether a modification of any type is present at the queried site. First, to capture the aggregate evidence of modifications across reads, RawMod builds a per-site pileup in which the *rows* hold the reference sequence and the raw signals of many reads around the site (Section 2.1). From this pileup, RawMod builds several features, called channels, such as raw signal amplitudes, bases, and how long each base stays in the nanopore (i.e., dwell time). Second, to compute the modification likelihood for each site from these channels, a convolutional encoder converts each row of the pileup into an embedding. A transformer with cross-read attention aggregates these embeddings into a site embedding that reflects how the signals *collectively* differ from the reference, from which a linear head computes the site’s likelihood (Section 2.2). Third, to enable detecting modifications regardless of their type, RawMod trains the model with classification, supervised contrastive, and anomaly-detection objectives, so that any modification can stand out from unmodified sites (Section 2.3). To learn the modification signal before sequence shortcuts, training first uses pairwise sites, whose modified and unmodified members share the same sequence, and then all data.

**Figure 2.**
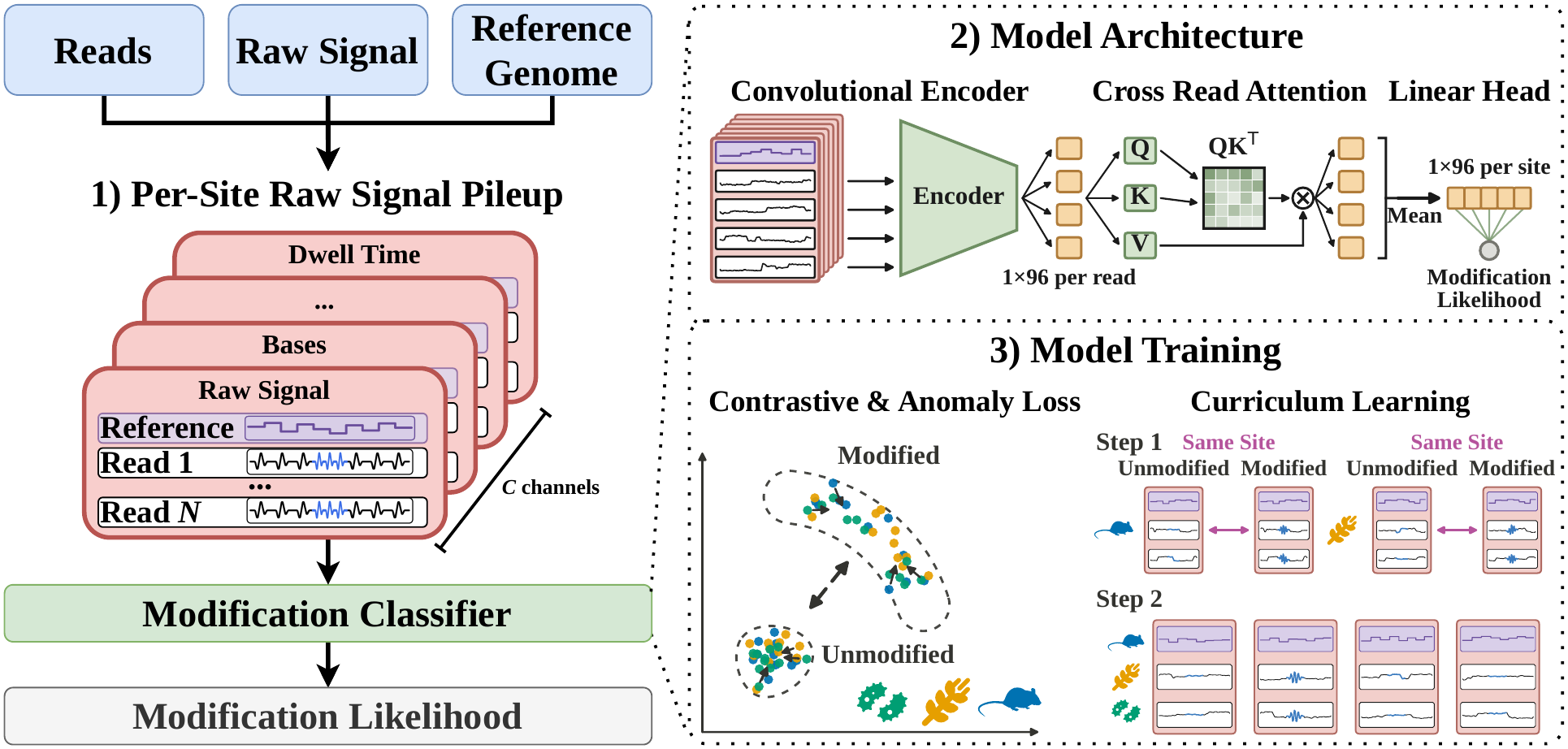
Overview of RawMod. (1) Reads, their raw signals, and the reference sequence form a raw signal pileup of up to *N* **reads and** *C* channels. (2) A convolutional encoder embeds each row of the pileup. A cross-read transformer combines the row embeddings into a site embedding, and a linear head gives the modification likelihood. (3) The model is trained with classification, contrastive and anomaly-detection objectives, first on pairwise sites (step 1) and then on all data (step 2).

### 2.1. Per-Site Raw Signal Pileup

RawMod predicts the modification likelihood of each site from a *raw signal pileup* of many signals aligned to a reference genome. The goal of the pileup is to separate the signal changes caused by a modification, which repeats across the reads of a site, from the noise of single reads, which differ between reads and datasets. RawMod builds the pileup in five steps.

First, RawMod takes the raw signals of the reads, their base-calls aligned to the reference genome, and the reference sequence as input to place the signal of every read on the reference. A modification changes the signal only slightly compared to the noise in a single read. This change differs across modification types and can appear at neighboring bases as well (Fig. 1b). RawMod normalizes the signal of each read so that reads from different runs share one scale. Second, to compare each read with an unmodified baseline, RawMod converts the reference sequence to its *expected* nanopore signal with a *k-mer level table* [76, 78], which provides the expected signal of each k-mer inside the nanopore. Third, for each reference base that a read covers, RawMod identifies the segment of the read’s signal that corresponds to that base. These segments are critical because DNA molecules do not move through the nanopore at a constant speed, so each base produces a different number of signal samples. The basecaller’s move table gives a first segmentation, and Remora [73] refines its boundaries against the same k-mer level table. RawMod requires this refinement for accurate detection (Supplementary Section S10.5). Fourth, since each base has a different number of signals, RawMod re-samples the signal of each base to the same number of samples, *L*=10, so that the signals of different reads align in the pileup.

Fifth, RawMod represents these signals as a pileup within a window of *W* =21 bases around the site. The first row holds the expected signal of the reference, and each subsequent row holds the signal of one read. A pileup holds up to *N* =15 reads of one strand. We keep the strands apart because a modification on one strand does not change the signal of the other. A site with more than *N* reads is split into random disjoint pileups whose scores are averaged, so that every read is used once, and the empty rows of a site with fewer reads are masked. The result is the pileup **X**_*i*_ ∈ ℝ^*C*×*H*×*W* ⋅*L*^ with *H*=*N* +1 rows.

#### Feature Extraction

Like the pileup images of DeepVariant and DeepSomatic [79, 80], RawMod constructs *C*=11 features (i.e., channels) for each row of the pileup:

- *Signal (2 channels):* the normalized current (*L* samples per base) and the log dwell time of each base.
- *Base (4 channels):* the one-hot reference base, which tells the model which bases surround the site.
- *Alignment (3 channels):* the strand, mapping quality, and whether the read covers the position.
- *Deviation (2 channels):* the difference between the read’s signal and the expected signal, at each position and at the site alone. These channels show the contrast between the read and an unmodified counterpart.

The reference row holds only the expected signal of the reference sequence, with the other channels set to zero. Supplementary Section S8.7 gives the remaining details of the pileup.

### 2.2. Model Architecture

RawMod turns the pileup of a site into its modification-agnostic likelihood in three steps (Fig. 2).

First, a convolutional encoder embeds each row of the pileup on its own. Each row, the reference row included, is a sequence of *W* ⋅*L* signal values with *C* channels, and the encoder, built from a three-layer convolutional stem, one Inception-style multi-branch block, and one reduction block [81, 82], converts it into one embedding vector **e**_*r*_ ∈ ℝ^96^ per row (Supplementary Section S8.7). We use convolutions because the signature of a modification is a local pattern spread over a few neighbouring bases, whose position inside the window differs between modification types (Fig. 1b), which a convolution can detect at any position of the window. Since the encoder runs on one row at a time, the resulting embedding also serves as a per-read modification-aware embedding which can be used for further classification tasks, as explored in Section 3.6.

Second, a transformer with cross-read attention compares the row embeddings to each other. We use attention across reads because the reads of a site differ in quality and strength of modification signal, which attention can consider better than a mean pool. A learned row embedding is added to each row so that the reference row is identifiable, and a two-layer transformer encoder lets each read attend to the reference and to the other reads, while a mask excludes the empty rows.

Third, a masked mean over the rows, followed by layer normalization, produces the site embedding **z**_*i*_, and a linear head maps it to the logit *f*_*θ*_(**X**_*i*_). Its sigmoid, *p*_*i*_ = *σ*(*f*_*θ*_(**X**_*i*_)), is the modification likelihood of the site, which we call its *site score*. The masked mean works for any number of reads. Since training also masks reads at random, one model learns to score sites of any coverage, down to a single read. The complete model has only 230,697 parameters because the pileup already places the signal of every read on the reference, so the model only has to compare aligned signals.

### 2.3. Model Training

The goal of training is a model that detects any modification type, including types that are absent from training. A classifier trained only to separate the labeled types can instead learn shortcuts to the label features that are specific to their sequence motif or the library they come from. We therefore train with three objectives intended to shape where sites land in the embedding space (Section 2.3.1), and in two stages that control which contrasts the model learns from first (Section 2.3.2).

#### 2.3.1. Training Objectives

The main objective, ℒ_BCE_, is a binary cross-entropy on the logit, which teaches the model to separate modified from unmodified sites. We add two auxiliary objectives to shape the space of the site embeddings. They make the embeddings of modified sites group together across datasets and those of unmodified sites group into a reference set, so that a modification type absent from training can still stand out by its distance from this reference set. For a mini-batch ℬ with labels *y*_*i*_, RawMod minimizes

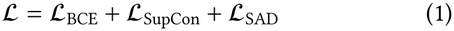

ℒ_SupCon_ is the supervised contrastive loss [83] on the *ℓ*_2_-normalized projection **q**_*i*_ of the site embedding with temperature *τ*, which pulls together sites that share a label across modification types and organisms so that the embedding groups sites by modification presence rather than by dataset.

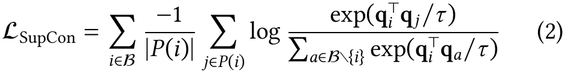

where *P* (*i*) is the set of other batch members with label *y*_*i*_

Deep Semi-Supervised Anomaly Detection (SAD) [84] pushes modified site embeddings away from the reference center, creating a compact reference set from which a modification of any type can stand apart. First, a linear map *h* projects each site embedding into a low-dimensional space. Unmodified sites are pulled toward a fixed center **c**, the mean of *h* over unmodified training pileups at the start of each training stage. Modified sites are treated as anomalies and pushed until they are a margin *m* away from the center.

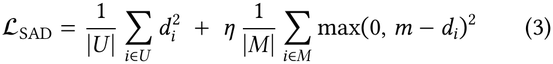

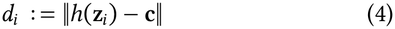

*U* and *M* are the unmodified and modified sites of the batch, and *η* weights the two terms. The hinge stops rewarding a modified site once it is *m* away from **c**, so that clearly modified sites cannot dominate the gradient. Since only the distance from **c** matters, modified sites do not group by type.

#### 2.3.2. Curriculum Learning

The goal of the curriculum is for the model to learn how a modification disturbs the signal before it can use sequence and organism-specific context as a shortcut. In most datasets, modified and unmodified sites also differ in sequence context or library, so a model trained on all data at once can separate the labels by these shortcuts instead. Thus, RawMod is trained in two stages. The first stage trains only on *pairwise sites*, sites sequenced in both a modified and an unmodified library (Supplementary Table S11). At pairwise sites, both labels come from the same position on the same genome, so the sequence context cannot predict the label. The pairwise labels come from synthetic constructs [85], PCR amplicons, or WGA libraries [86]. The second stage trains all data, which adds modifications and organisms that have no pairwise control. Supplementary Table S3 tests the curriculum and objectives, which change accuracy no more than the training seed does, unlike the deviation channels.

#### 2.3.3. Optimization and Data Preparation. RawMod is trained with AdamW, whose settings and schedule are listed in Supplementary Section S8.7. Data augmentation masks a random 0–30% of read rows for generalization to low coverage and adds Gaussian noise to the current channel

## 3. Results

### 3.1. Evaluation Methodology

We evaluate RawMod as a modification-agnostic caller in three parts. First, Section 3.2 compares RawMod with released, fine-tuned, and comparative callers inside genomic regions with-held from training across modification types. Second, Section 3.3 summarizes how RawMod generalizes to withheld modification types, to low coverage, and to limited labels of a new type. Third, Sections 3.5 and 3.6 apply RawMod to create a control-free 5hmU modification map and to perform label-free read-origin typing with RawMod embeddings.

### Implementation

RawMod is implemented in Python with PyTorch and released with its trained models and the scripts that regenerate every table and figure of this paper at https://github.com/STORMgroup/RawMod. Models train on one NVIDIA RTX A5000 GPU, and cost experiments use it with 8 cores of an AMD EPYC 7352 CPU. We 1) basecall reads with Dorado (R10.4.1, 5 kHz, `sup` model), and align with minimap2 [87], and 2) refine the signal-to-base boundaries with Remora [73] against the R10.4.1 k-mer level table released by Uncalled4 [76]. Supplementary Tables S14 and S13 list every tool version, model, and set of parameters.

### Training

We train on the datasets of Supplementary Table S11. Every pileup is labeled modified or unmodified at the site level. Training follows the two stages of Section 2.3.2, with hyperparameters shown in Supplementary Section S8.7. To keep the test data unseen, we remove a contiguous region of about 15% of every genome from training, together with every read that overlaps it, and we evaluate only inside these regions. Supplementary Section S9.7 lists the regions used. For the generalization experiments, we additionally withhold one modification type from training and report the mean over three random seeds. To test generalization across datasets, we instead withhold one organism or library. We use the released model trained on all data to map 5hmU in the dinoflagellate genome and to separate reads by sample without labels. On SPO1, whose reads it saw, we use the region-holdout model.

### Datasets

Supplementary Table S11 shows the details of 15 benchmark datasets that cover five modification types (i.e., 5mC, 5hmC, 4mC, 6mA and 5hmU) across bacterial, phage, plant, mouse, human and synthetic DNA. Supplementary Section S9.6 gives the source of the modification labels of every dataset. We sequence and release the novel native and PCR SPO1 libraries. Supplementary Section S8.3 describes their preparation, sequencing, and preprocessing.

### Compared tools

We compare RawMod with state-of-the-art tools in three ways. First, we identify the modified sites by using the modification-specific models of the tools Dorado [31], UniMeth [61], DeepMod2 [58], and Rockfish [48]. Second, we use Uncalled4 [76] and NovaMod [77] with their matched *unmodified control* of the same genome. Third, we fine-tune UniMeth on RawMod’s own training data, 1) for each of the three modification types without a released UniMeth model (UniMeth-FT) and 2) on all modification types together (UniMeth-Agn), to test RawMod’s advantage when a state-of-the-art model is trained on the *same* data. Most of these tools identify modified sites on each read, whereas RawMod scores each site from a pileup of reads. We therefore score every tool on the same 10 reads per site, which also removes coverage as a variable. A tool’s site score is the mean of its per-read probabilities over these reads, with a read it does not call counted as unmodified. RawMod scores the pileup of the same 10 reads. All sites lie inside the held-out regions, which neither RawMod nor the fine-tuned models were trained on. Supplementary Section S9 gives the full procedure.

### Metrics

We measure accuracy with the area under the receiver operating characteristic curve (AUROC) at the site level. For each tool, we report the AUROC of each dataset, its mean and its lowest value over the five modification types, and a pooled AUROC that applies one threshold to all datasets, with each type weighted equally. Because 8 of the 15 datasets are 5mC, we weight each modification type equally rather than each dataset, so that no single type dominates these summaries. We also report the fraction of modified sites that a tool detects at a false positive rate of 1% for a threshold analysis. To test whether a tool reads the modification rather than the library, we compute the AUROC between a native library and its un-modified control at positions that carry no known modification. A tool that ignores the library gives an AUROC of 0.5. We report the library invariance 1 − 2 |AUROC − 0.5|, which is 1 for such a tool and 0 for a tool that separates the libraries perfectly. For read origin, we report the Adjusted Rand Index between the label-free partition of the reads and their true origin, and the oriented AUROC max(AUC, 1−AUC) of the partition score, both per window. We orient the AUROC because a label-free partition does not contain a group label. We measure computational cost as wall time, CPU time, peak memory and GPU busy time on the same 13,110 *H. pylori* reads for every tool. Supplementary Section S10.4 gives the full setup.

#### 3.2. Comparison with Modification-Specific Callers

Table 1 shows the site-level accuracy of each model on the same held-out genomic sites. Figure 3 combines the models of each tool into one score and compares the tools across all datasets and at the positions without a known modification, to test library invariance. We make three key observations.

**Table 1.**
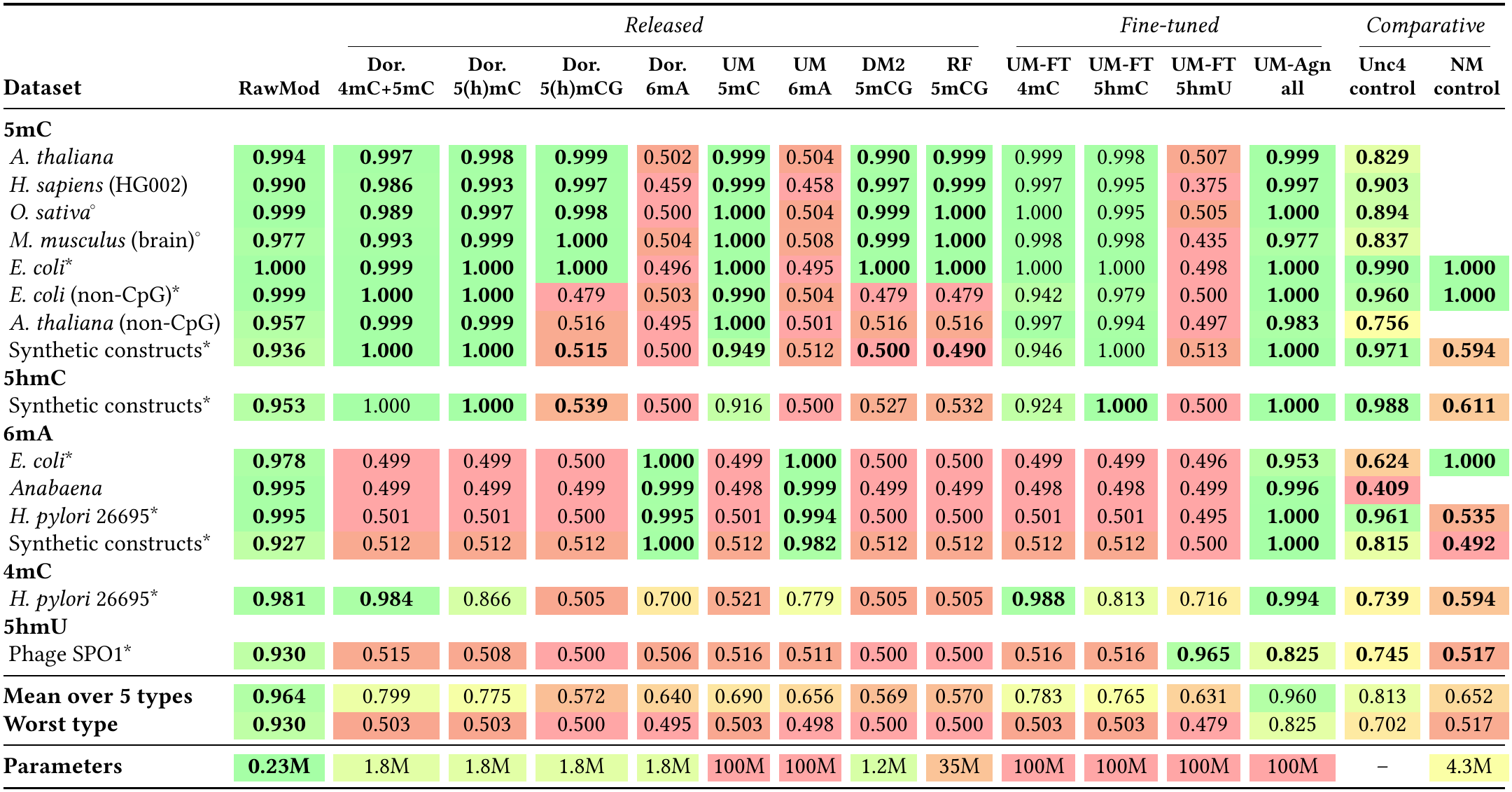
Site-level AUROC of each modification model inside held out genomic regions, from 10 reads per site. Each column shows one model. Dor.: Dorado, UM: UniMeth, DM2: DeepMod2, RF: Rockfish, UM-FT: UniMeth fine-tuned on one modification type, UM-Agn: UniMeth fine-tuned on all types, Unc4: Uncalled4, NM: NovaMod. A blank cell has no matched unmodified library, on which we train NovaMod. Mean and worst are over the five modification types and use all 15 datasets. ^°^Absent from RawMod’s training. ^∗^**Scored against a matched unmodified library (Supplementary Section S9.6). Bold: trained for the row’s modification. Further information in Supplementary Section S9.9**.

| **Dataset** | **RawMod** | *Released* |  |  |  |  |  |  |  | *Fine-tuned* |  |  |  | *Comparative* |  |
|---|---|---|---|---|---|---|---|---|---|---|---|---|---|---|---|
|  |  | **Dor.<br>4mC+5mC** | **Dor.<br>5(h)mC** | **Dor.<br>5(h)mCG** | **Dor.<br>6mA** | **UM<br>5mC** | **UM<br>6mA** | **DM2<br>5mCG** | **RF<br>5mCG** | **UM-FT<br>4mC** | **UM-FT<br>5hmC** | **UM-FT<br>5hmU** | **UM-Agn<br>all** | **Unc4<br>control** | **NM<br>control** |
| **5mC** |  |  |  |  |  |  |  |  |  |  |  |  |  |  |  |
| *A. thaliana* | **0.994** | **0.997** | **0.998** | **0.999** | 0.502 | **0.999** | 0.504 | **0.990** | **0.999** | 0.999 | 0.998 | 0.507 | **0.999** | **0.829** |  |
| *H. sapiens* (HG002) | **0.990** | **0.986** | **0.993** | **0.997** | 0.459 | **0.999** | 0.458 | **0.997** | **0.999** | 0.997 | 0.995 | 0.375 | **0.997** | **0.903** |  |
| *O. sativa*<sup>°</sup> | **0.999** | **0.989** | **0.997** | **0.998** | 0.500 | **1.000** | 0.504 | **0.999** | **1.000** | 1.000 | 0.995 | 0.505 | **1.000** | **0.894** |  |
| *M. musculus* (brain)<sup>°</sup> | **0.977** | **0.993** | **0.999** | **1.000** | 0.504 | **1.000** | 0.508 | **0.999** | **1.000** | 0.998 | 0.998 | 0.435 | **0.977** | **0.837** |  |
| *E. coli*<sup>\*</sup> | **1.000** | **0.999** | **1.000** | **1.000** | 0.496 | **1.000** | 0.495 | **1.000** | **1.000** | 1.000 | 1.000 | 0.498 | **1.000** | **0.990** | **1.000** |
| *E. coli* (non-CpG)<sup>\*</sup> | **0.999** | **1.000** | **1.000** | 0.479 | 0.503 | **0.990** | 0.504 | 0.479 | 0.479 | 0.942 | 0.979 | 0.500 | **1.000** | **0.960** | **1.000** |
| *A. thaliana* (non-CpG) | **0.957** | **0.999** | **0.999** | 0.516 | 0.495 | **1.000** | 0.501 | 0.516 | 0.516 | 0.997 | 0.994 | 0.497 | **0.983** | **0.756** |  |
| Synthetic constructs<sup>*</sup> | **0.936** | **1.000** | **1.000** | **0.515** | 0.500 | **0.949** | 0.512 | **0.500** | **0.490** | 0.946 | 1.000 | 0.513 | **1.000** | **0.971** | **0.594** |
| **5hmC** |  |  |  |  |  |  |  |  |  |  |  |  |  |  |  |
| Synthetic constructs<sup>*</sup> | **0.953** | 1.000 | **1.000** | **0.539** | 0.500 | 0.916 | 0.500 | 0.527 | 0.532 | 0.924 | **1.000** | 0.500 | **1.000** | **0.988** | **0.611** |
| **6mA** |  |  |  |  |  |  |  |  |  |  |  |  |  |  |  |
| *E. coli*<sup>\*</sup> | **0.978** | 0.499 | 0.499 | 0.500 | **1.000** | 0.499 | **1.000** | 0.500 | 0.500 | 0.499 | 0.499 | 0.496 | **0.953** | **0.624** | **1.000** |
| *Anabaena* | **0.995** | 0.499 | 0.499 | 0.499 | **0.999** | 0.498 | **0.999** | 0.499 | 0.499 | 0.498 | 0.498 | 0.499 | **0.996** | **0.409** |  |
| *H. pylori* 26695<sup>\*</sup> | **0.995** | 0.501 | 0.501 | 0.500 | **0.995** | 0.501 | **0.994** | 0.500 | 0.500 | 0.501 | 0.501 | 0.495 | **1.000** | **0.961** | **0.535** |
| Synthetic constructs<sup>*</sup> | **0.927** | 0.512 | 0.512 | 0.512 | **1.000** | 0.512 | **0.982** | 0.512 | 0.512 | 0.512 | 0.512 | 0.500 | **1.000** | **0.815** | **0.492** |
| **4mC** |  |  |  |  |  |  |  |  |  |  |  |  |  |  |  |
| *H. pylori* 26695<sup>\*</sup> | **0.981** | **0.984** | 0.866 | 0.505 | 0.700 | 0.521 | 0.779 | 0.505 | 0.505 | **0.988** | 0.813 | 0.716 | **0.994** | **0.739** | **0.594** |
| **5hmU** |  |  |  |  |  |  |  |  |  |  |  |  |  |  |  |
| Phage SPO1<sup>*</sup> | **0.930** | 0.515 | 0.508 | 0.500 | 0.506 | 0.516 | 0.511 | 0.500 | 0.500 | 0.516 | 0.516 | **0.965** | **0.825** | **0.745** | **0.517** |
| **Mean over 5 types** | **0.964** | 0.799 | 0.775 | 0.572 | 0.640 | 0.690 | 0.656 | 0.569 | 0.570 | 0.783 | 0.765 | 0.631 | 0.960 | 0.813 | 0.652 |
| **Worst type** | **0.930** | 0.503 | 0.503 | 0.500 | 0.495 | 0.503 | 0.498 | 0.500 | 0.500 | 0.503 | 0.503 | 0.479 | 0.825 | 0.702 | 0.517 |
| **Parameters** | **0.23M** | 1.8M | 1.8M | 1.8M | 1.8M | 100M | 100M | 1.2M | 35M | 100M | 100M | 100M | 100M | – | 4.3M |

**Figure 3.**
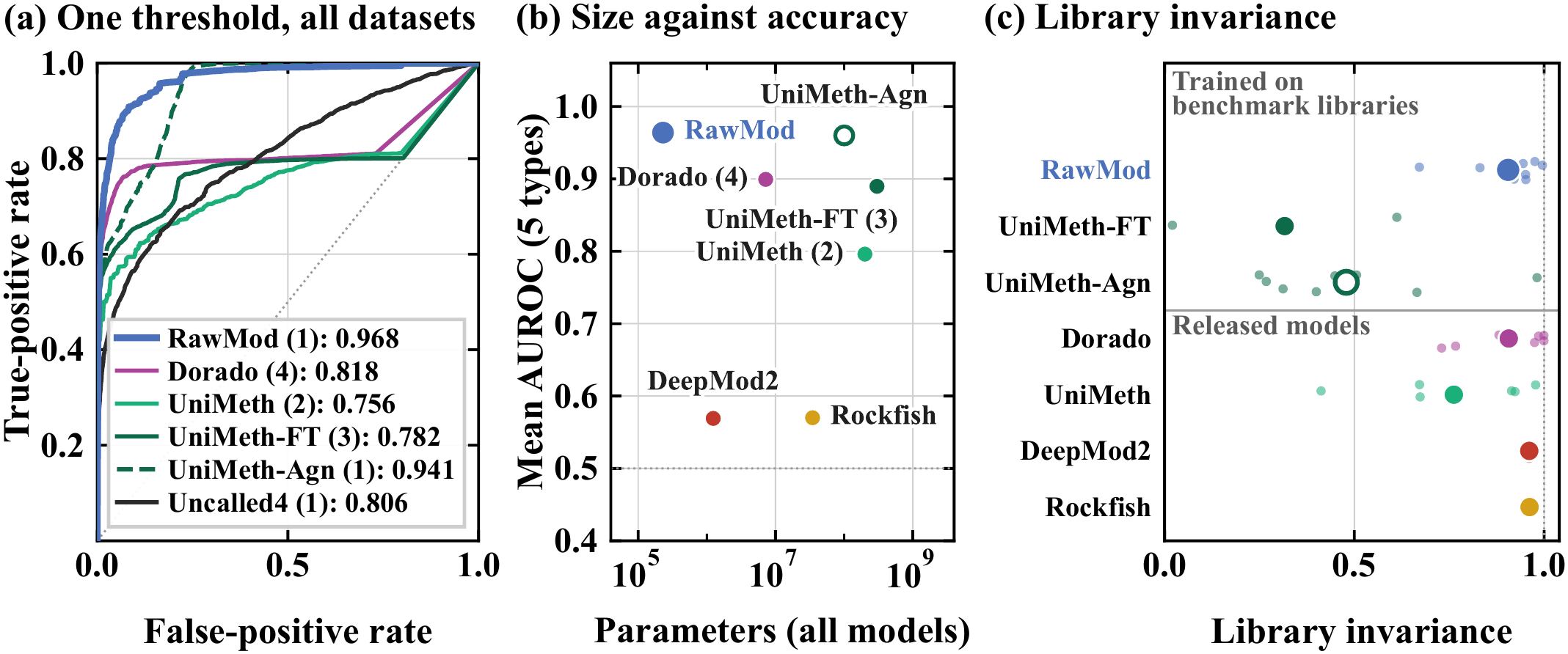
One RawMod model against every tool inside held-out genomic regions. (a) Pooled ROC curves across all datasets. (b) Parameters of all models for a tool versus its mean AUROC over them. (c) Library invariance between a native library and its unmodified control at positions without a known modification. Small points show each dataset. Models above the line were trained on the benchmark libraries. Tool abbreviations as in Table 1. Notes in Supplementary Section S9.9.

First, a single RawMod model detects *all* five modification types, whereas *no* released model generalizes beyond the base it was trained for. RawMod scores a 0.964 mean AUROC over the five modification types, against 0.799 for the next highest released model (Dorado 4mC+5mC), and 0.899 for all four Dorado models combined, which lack 5hmU. RawMod’s weakest modification type is 5hmU at 0.930, whereas the weakest type of every released model is *at most* 0.503.

Second, RawMod’s scores are comparable across modification types, which lets one threshold serve all datasets. With each modification type weighted equally, its pooled AUROC is 0.968 (95% CI 0.958–0.975), against 0.818 (0.816–0.820) for all Dorado models pooled and 0.756 (0.751–0.761) for UniMeth’s two released models. At one threshold set to 1% false positives on the unmodified sites, RawMod detects 60% of the modified sites averaged over the five modification types, against 57% for Dorado’s four models. Dorado detects only 16% of the 4mC sites and none of the 5hmU sites, for which it has no model, against 66% and 58% for RawMod. Supplementary Table S23 gives this comparison across every dataset. One RawMod threshold also separates 6mA and 5mC on the same *E. coli* genome (AUROC 0.987). Each released model detects only its own modification (0.49–0.69), and all four Dorado models together reach 0.999. Supplementary Table S24 gives the score of every model. Third, RawMod uses 0.23M parameters, 31× and 435× fewer than Dorado’s four models and each UniMeth model, yet is at least as accurate (Fig. 3b).

RawMod has two limitations in this comparison. First, Supplementary Table S20 shows that the mean AUROC over the five types drops when the organism or library of a dataset is withheld from training, mainly for 6mA, 4mC and 5hmU. For 4mC and 5hmU, the withheld library is the only source of the modification, so withholding it removes the type from training, which Section 3.3 examines. Second, Dorado and UniMeth remain more accurate than RawMod for their own modification types and sequence contexts (mean AUROC above 0.99), as Supplementary Section S9.9 shows for each specialist.

To test whether RawMod’s results come from its design or from its training data, we compare it with UniMeth fine-tuned on the same data, UniMeth-FT and UniMeth-Agn. We make two key observations. First, UniMeth-Agn, the only other control-free modification-agnostic model tested, is slightly worse than RawMod, despite 435× more parameters. It reaches a mean AUROC of 0.960 over the five modification types against RawMod’s 0.964, a difference within the bootstrap interval (−0.009 to +0.013), and a type-balanced pooled AUROC of 0.941 (0.937–0.945) against RawMod’s 0.968 (0.958–0.975). Its weakest type, 5hmU, reaches 0.825 against RawMod’s weakest, 0.930 (difference +0.069 to +0.117). UniMeth-FT, fine-tuned on 5hmU alone, reaches 0.965 there. Second, UniMeth-Agn’s accuracy comes partly from library recognition rather than modification identification. At unmodified positions of modified datasets, UniMeth-Agn and UniMeth-FT tell a native library from its unmodified control (mean AUROC 0.76 and 0.84, Fig. 3c), whereas RawMod (0.48) and the released models (0.45–0.50) do not. UniMeth-Agn and UniMeth-FT therefore also learn dataset-specific features, not only the presence of a modification. Scored against each library’s own unmodified background, where the library gives no cue, the mean AUROC of UniMeth-Agn falls to 0.917, while RawMod stays at 0.948. Supplementary Section S9.3 tests this for every dataset.

The speed of a modification-agnostic caller matters because it must score every position of a genome rather than only known motifs. We make three key observations. First, RawMod is faster than tools covering several modifications. On 13,110 *H. pylori* reads, RawMod takes 5.4 min (8.6 min with basecalling), against 9.9 min for Dorado’s four models and 37.9 min for UniMeth’s two. Every additional modification adds another genome-wide pass for the other tools, but not for RawMod. Second, RawMod uses about a tenth of Dorado’s GPU time (0.9 against 9.7 min), because it builds its pileups on the CPU. For the same reason, it uses 17.4 GB of memory with eight parallel processes, against Dorado’s 4.5 GB. Third, RawMod keeps its accuracy on datasets with the faster `hac` and `fast` basecalls, which take 12–36% of the `sup` basecalling time, but requires the Remora refinement of the signal boundaries. Without it, the mean AUROC falls from 0.974 to 0.725 with `sup` basecalls and to near chance with the faster ones. Supplementary Table S28 gives the results.

### Key Result 1: One RawMod model replaces a set of modification-specific callers

One RawMod model detects the presence of all five modification types, including 5hmU, which no released caller detects. Under one joint threshold, it is more accurate than any released caller, even when combining all of its models. With 31× fewer parameters than Dorado’s four models, RawMod runs faster and uses less GPU time.

#### 3.3 Modification and Dataset Generalization

A modification-agnostic caller should also detect modification types absent from its training. Fig. 4 shows how RawMod generalizes to a modification type withheld from training, to a new modification type with few labeled sites, and to fewer reads per site. Section 3.4 and Supplementary Section S1 give the details. This test withholds the modified sites of one type, so it measures transfer across modification types rather than across organisms. We make three key observations. First, transfer to an unseen modification occurs within a base. Withholding one modification type from training leaves three of the five types detected, with a mean zero-shot AUROC of 0.792 over three seeds across all five types. 5hmC (0.983), 5mC (0.978) and 4mC (0.906) remain detected when another cytosine modification stays in training. In contrast, 6mA (0.594) and 5hmU (0.497), the only modifications of their base in our data, do not transfer. The released model is trained on modifications of A, C and T, so a new modification of these bases has a relative in training. Modified guanines are rare in DNA. The best known, the 7-deazaguanines of some phages and bacteria [88], have no relative in training, so we do not expect them to transfer. Second, a modification without a relative in training can be added with about 3,000 labels. Retraining with them raises the 5hmU AUROC to 0.82. The need for so many labels is consistent with the small change that 5hmU causes in the signal at the modified base (Fig. 1b). Third, RawMod is robust to low coverage. The `mixed` model reaches an AUROC of 0.898 from a single read and 0.987 from 15 reads, as Fig. 4b shows.

**Figure 4.**
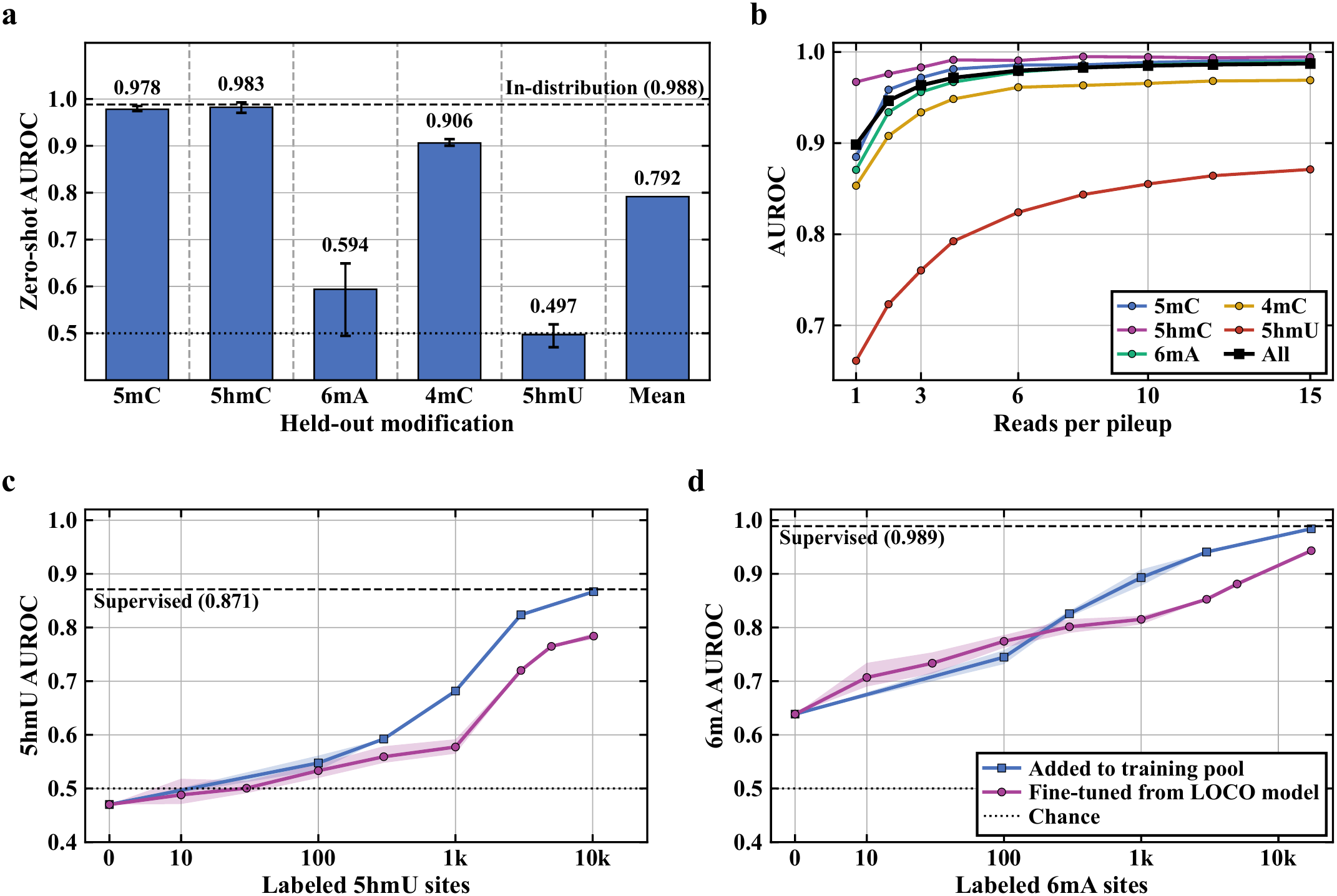
Generalization of RawMod. (a) Zero-shot AUROC of each held-out modification over 3 seeds, `mixed` **at dashed line. (b)** `mixed` **AUROC against reads per pileup. (c) 5hmU AUROC against labeled 5hmU sites, from a support half of 10**,**173 positive positions. (d) 6mA AUROC against labeled 6mA sites, from a support half of 17**,**366 positive positions. In (c, d), bands show the range over draws and the dashed lines show the** `mixed` **baseline. Supplementary Section S2.1 gives the protocol**.

#### 3.4 Few-Shot Detection of 5hmU and 6mA

Since 5hmU does not transfer zero-shot, we measure how many labeled 5hmU sites RawMod requires to learn to detect the modification. We draw the labeled sites from the 5hmU positions outside the held-out regions, a support half of 10,173 positive positions chosen by coordinate hash, and we evaluate every model inside the held-out regions on the 5hmU test set of Supplementary Table S1. We then provide *K* labeled 5hmU positions from the support half, each paired with *K* control positions, in two ways. First, we fine-tune the 5hmU `LOCO` model on the labeled positions together with replayed training pileups of the other modifications. Second, we add the labeled positions to the training pool and retrain the `LOCO` model from scratch. In both, *K* ranges from 10 to all 10,173 positions. Supplementary Section S2.1 provides the details and Figure 4c shows the results. We make four key observations.

First, RawMod needs thousands of labels to detect 5hmU. With fine-tuning, the AUROC stays near chance with 10 and 30 sites (0.488 and 0.501) and rises only to 0.533 with 100 and 0.577 with 1,000 sites; it reaches 0.720 with 3,000 sites and 0.784 with all 10,173. We attribute this to the small change that 5hmU causes in the signal at the modified base (Fig. 1b) and to the absence of any other thymine modification in training.

Second, retraining is better than fine-tuning once about 1,000 labeled sites are available. With 100 and 300 sites the two approaches agree within variance (0.548 against 0.533, and 0.592 against 0.559). With 1,000 sites, retraining reaches 0.682 against 0.577 with fine-tuning, with 3,000 sites 0.824 against 0.720, and with all 10,173 sites 0.867 against 0.784. With all sites, retraining matches the `mixed` model on the same positions (0.871), even though that model saw twice as many 5hmU positions in training, as Supplementary Section S2.1 describes.

Third, 6mA follows the same pattern as 5hmU, from a higher starting point (Fig. 4d). We repeat both experiments for the 6mA `LOCO` model, with a support half of 17,366 positive positions. Fine-tuning raises 6mA from 0.639 to 0.707 with 10 sites, 0.815 with 1,000 and 0.943 with all 17,366. Retraining is better from about 300 sites on (0.826 against 0.801), and reaches 0.893 with 1,000 sites, 0.941 with 3,000 and 0.984 with all, which is close to the `mixed` model on the same positions (0.989).

Fourth, the other three modifications need few or no labels (Supplementary Fig. S1). Inside the held-out regions, 5mC and 5hmC start at 0.974 and 0.971 and gain at most 0.024 with 1,000 sites; 4mC improves from 0.914 to 0.975. Only the two modifications without a relative in training, 5hmU and 6mA, need thousands of labeled sites.

We conclude that a modification without a chemical relative in training can be added to RawMod with ∼3,000 labeled sites (AUROC 0.82 when retrained with them, 0.72 when fine-tuned), and that a few hundred sites raise the AUROC to at most ∼0.6 with either approach for 5hmU. For 6mA, 1,000 labeled sites reach 0.89 when retrained and 0.82 when fine-tuned. With all labels, retraining comes within 0.005 of the supervised model.

### Key Result 2: A new modification requires thousands of labels instead of a new model

5hmU, which has no relative in training, reaches an AUROC of 0.82 when RawMod is retrained with ∼3,000 labeled sites. Fine-tuning reaches 0.72. A few hundred sites raise the AUROC to at most ∼0.6. 6mA reaches 0.89 with 1,000 labeled sites and 0.98 with all.

### 3.5. Control-Free 5hmU Map of a Dinoflagellate Contig

The goal of this experiment is to map a modification that no released caller detects, in a genome without an unmodified control. In *Amphidinium carterae*, 45% of thymines are 5hmU by mass spectrometry [67]. 5hmU has been proposed to play a role in the evolution of the dinoflagellate nucleus [89]. Previous studies use 5hmU immunoprecipitation (5hmU-DIP) and chemical labeling to find 5hmU in transposable elements (TEs) rather than exons, with more 5hmU on younger TE copies [67]. In *Breviolum minutum*, chemical mapping finds 5hmU enriched at the boundaries of the chromatin domains that form where gene arrays converge [90, 91]. Neither assay resolves single molecules or bases, no released caller has a 5hmU model, and an unmodified control is difficult to produce for a 1.25 Gb repeat-rich genome. We therefore test whether a control-free map reproduces these findings and whether its per-base resolution reveals patterns that previous assays miss.

We build the map in three steps. First, we take contig_3762 (GenBank JBNFNT010000839.1, 7.73 Mb), the largest contig of a recent R10.4.1 nanopore assembly of *Amphidinium carterae* strain NCMA 1314 [92], which has 237 genes that we annotated manually. Supplementary Section S8.4 describes the contig and its annotation. Second, we run RepeatMasker [93] to identify TEs along the contig. Third, we score all 2,170,929 forward-strand thymines with at least 10 reads (98.7% of the contig) with the model trained on all data (Section 2.3), which never saw this genome. We read the thymine scores as 5hmU, the only known thymine modification in *A. carterae*.

We make three key observations: the first two are consistent with published trends, and the third is new.

First, 5hmU is depleted in coding sequence. Intergenic thymines score 1.06× higher than thymines in coding sequences (0.713 versus 0.670, *p* = 0.003 against a circular-shift null), and 1.12–1.17× after the scores are residualized on the 5-mer and local GC content, which differ between coding and intergenic DNA. Supplementary Table S7 gives this correction.

Second, younger TEs carry more 5hmU. Thymines in repeat copies within 5% of their family consensus from RepeatMasker score 0.773 on average, against 0.680 for thymines outside repeats, as Supplementary Fig. S5c shows.

Third, the map shows one pattern that the published assays did not report. The two divergent strand-switch regions of the contig, 84 and 92 kb long, where adjacent gene arrays point away from each other, score 0.928 and 0.851, the highest and third-highest of the contig’s 102 intergenic regions, as Supplementary Fig. S5d shows. The two convergent regions, where gene arrays point to each other, score in the 89th and 49th percentiles. The contig holds only two junctions of each type, so we report this pattern as a preliminary hypothesis in need of more biologically validated data. Supplementary Table S6 gives every contrast we test against a circular-shift null. We hypothesize that 5hmU is concentrated in young, TE-rich DNA at divergent strand-switch regions in *Amphidinium carterae* and depleted from coding sequence.

Three properties of the data limit these claims: a draft assembly with manually annotated genes, a median read error of 5.4%, and published profiles from a different strain, which we therefore compare by feature class rather than by position. A full assembly and annotation would address them.

### Key Result 3: A control-free 5hmU map at per-base resolution

RawMod maps 5hmU on an *Amphidinium carterae* con-tig without a control library. The map is consistent with two published trends and leads to a testable hypothesis on 5hmU placement at divergent strand-switch regions.

### 3.6. Unsupervised Read-Origin Typing

The goal of read-origin typing is to separate the reads of one sample by their source, using only their modifications. Reads from one bulk sequencing sample can differ in their modifications while sharing an identical sequence. Tumors are the clearest case. Somatic mutations are sparse [94, 95], so most reads carry no somatic variant, but many carry an altered methylome [96]. On chr20 of COLO829 [97, 98], a human melanoma cell line, DeepSomatic [80] calls 1,110 somatic SNVs (17.2 per Mb), so a 10 kb window contains a somatic SNV with a probability of only 16%. cvlr [99] and NanoMethCluster [100] cluster nanopore reads on their methylation calls without labels, and MethylBERT [101] classifies them with labels. All three start from the calls of a released model for the modification they use. We therefore ask whether RawMod’s per-read embeddings can serve as a *fingerprint* to separate reads by origin, without methylation calls or sample labels. This matters most for 5hmU in SPO1, for which no released model exists.

We pool the reads of the COLO829 melanoma line [97, 98] and its matched lymphoblastoid line COLO829BL in 54 windows of 10 kb on chr20 with varying *CpG site* density, so that the windows span a range of differential methylation. A CpG site is differential for evaluation purposes when its bulk methylation differs by at least 50 percentage points between the two lines. In these windows, about one CpG in six is differential (1,564 of 8,852). RawMod partitions the reads of a window in two steps. First, at every CpG, we split the reads into two groups by the sign of the first principal component of their per-read embeddings. Second, a consensus over the CpGs of the window gives one partition of its reads, because the split at a single CpG is noisy. Supplementary Section S6 gives the consensus rule. Labels are used only to score the partition (Section 3.1). Supplementary Section S6 also gives the controls.

Figure 5 shows the agreement between the label-free partitions and read origin. The key result is that separation depends on how many CpGs differ in methylation between the origins (Spearman *ρ* =0.54, *p* =3×10^−5^ between the number of differentially methylated CpGs and ARI). The median ARI is 0.62 in the 33 windows with at least 20 differential CpGs, 0.03 in windows with 5–19, and 0.00 with fewer than 5.

**Figure 5.**
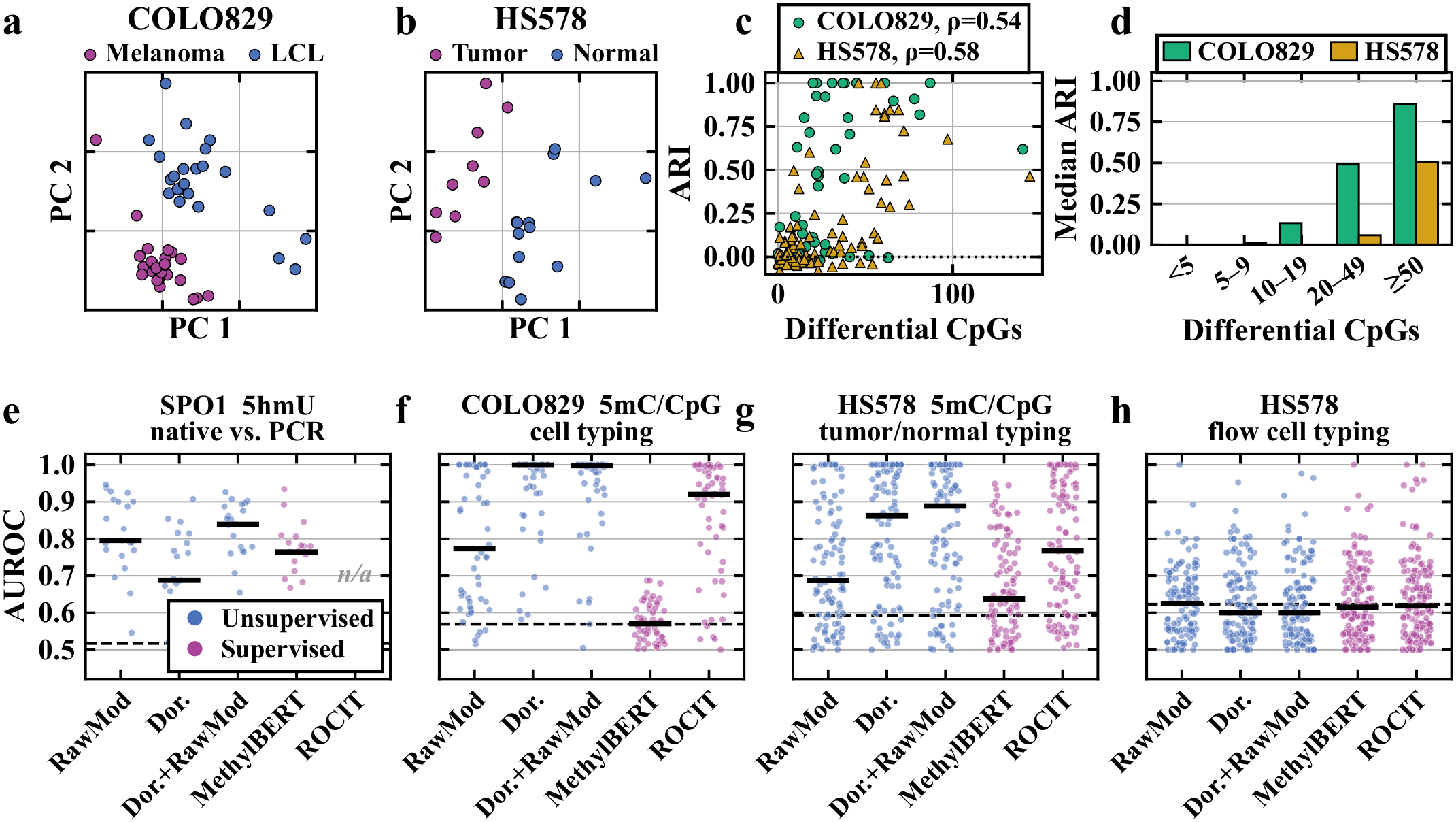
Label-free read-origin typing. (a, b) First two principal components of RawMod embeddings of the reads in one window of COLO829 and HS578. (c, d) ARI of read origin clustering against the number of differentially methylated CpGs per window. (e–h) Per-window AUROC of every method in SPO1 (native against PCR), COLO829 (cell typing), HS578 (tumor presence), and a control (flow cell typing). Dashed: null AUROC. Dor.: Dorado. Notes: Supplementary Section S6.

### 3.6.1. Tumor presence separation

COLO829 pairs a melanoma line with a lymphoblastoid line, so its two groups differ in cell type and tumor state. We therefore repeat on HS578 [80, 102], where HS578T is a breast carcinoma line and HS578Bst is a normal breast line from the same patient. Supplementary Section S6 shows the comparison of the cell lines. In 100 windows of 10 kb on chr1, about one CpG in four is differential (2,500 of 9,795). The same correlation with the number of differentially methylated CpG sites holds (Fig. 5d, Spearman *ρ* =0.58, *p* =2×10^−10^), with a median ARI of 0.50 in the 20 windows carrying at least 50 differential CpGs and −0.01 in the 20 windows carrying fewer than 5

### 3.6.2. Comparison across methods

To test what a modification-agnostic embedding costs against purpose-built methods, we compare RawMod on identical reads and sites with Dorado’s CpG calls under the same consensus rule and with supervised read classifiers MethylBERT and ROCIT. Table 2 and Fig. 5e–h show the per-window AUROC.

**Table 2.**
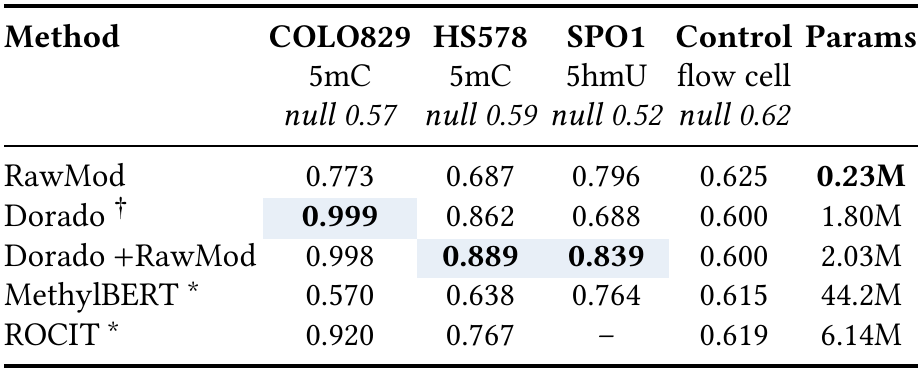
Per-window median oriented AUROC for separating the read origin of two samples. ^†^**CpG model on the human lines, all-context cytosine model on SPO1. Notes: Supplementary Section S6**. ^∗^**Supervised: trained with read labels. Params: parameters of the model each method runs; for Dorado, one modification model. On SPO1, RawMod is scored at T sites**.

| **Method** | **COLO829** | **HS578** | **SPO1** | **Control** | **Params** |
|---|---|---|---|---|---|
|  | 5mC | 5mC | 5hmU | flow cell |  |
|  | *null 0.57* | *null 0.59* | *null 0.52* | *null 0.62* |  |
| RawMod | 0.773 | 0.687 | 0.796 | 0.625 | **0.23M** |
| Dorado<sup>†</sup> | **0.999** | 0.862 | 0.688 | 0.600 | 1.80M |
| Dorado +RawMod | 0.998 | **0.889** | **0.839** | 0.600 | 2.03M |
| MethylBERT<sup>*</sup> | 0.570 | 0.638 | 0.764 | 0.615 | 44.2M |
| ROCIT<sup>*</sup> | 0.920 | 0.767 | – | 0.619 | 6.14M |

We make three key observations. First, Dorado calls partition better than RawMod embeddings on both cell lines, with median oriented AUROC 0.999 and 0.862, against RawMod’s 0.773 and 0.687, likely because its model is specially trained for 5mC at CpG sites. ROCIT, a supervised classifier pretrained on another tumor/normal pair, is also ahead of RawMod on both lines (0.920 and 0.767). Second, the separation is not a batch effect. We split the reads of each HS578 line by flow cell (Fig. 5h). Within one line, the two flow cells carry no biological difference, so any separation would be a batch effect. On these reads, every method stays at the chance level of oriented AU-ROC. Third, MethylBERT sits near the null on both cell lines, likely because it reads short (500 bp) fragments, which contain little evidence of differential methylation.

### 3.6.3. Native against PCR separation

Both cell lines carry human CpG methylation, for which a purpose-built Dorado caller exists. We therefore add a test on SPO1, a *Bacillus* phage whose genome carries 5hmU in place of every thymine, a modification for which no released model exists. We separate native reads from PCR copies of the same genome, which share their sequence but carry no 5hmU. Each of 21 windows of 1.2 kb compares one PCR library with native reads. We score each method at the base its model reads. RawMod scores 200 random thymine sites per window. Dorado has no thymine model, so we score 200 random cytosines with its closest model, the cytosine model 5mC_5hmC@v3. RawMod uses the region-holdout model of Section 3.2, which never saw one region of the genome. Supplementary Section S6 describes how we chose the windows. Figure 5e shows the per-window AUROC. We make three key observations.

First, RawMod at thymine sites reaches a median AUROC of 0.80, against 0.69 for Dorado’s cytosine model. RawMod is ahead of Dorado in 17 of the 21 windows (paired median difference +0.10, Wilcoxon signed-rank *p* = 0.002), including all three windows inside the held-out region. Second, the supervised classifiers do not perform better than RawMod. MethylBERT, fine-tuned on these reads with Dorado’s calls at CpG cytosines, reaches 0.76 on the same windows, likely because it reads Dorado’s cytosine calls and never sees the thymines that carry 5hmU. ROCIT cannot run on SPO1, because it requires somatic variants and a cell-type atlas, which SPO1 lacks. Third, RawMod embeddings transfer across datasets and experimental setups, and they can improve a specialist caller. The same RawMod method and consensus rule run on human cell lines and on a phage, without a specific model for the modification. Adding RawMod embeddings to Dorado’s calls raises Dorado from 0.69 to 0.84 on SPO1, ahead in 19 of the 21 windows (*p* = 5×10^−4^), where Dorado lacks a model for the modification. On HS578, the combination also raises the median AUROC of Dorado from 0.862 to 0.889, although this gain is not significant across windows. On COLO829, where Dorado is already near its ceiling, the two are level (0.998 against 0.999). RawMod needs only reads, raw signal and a reference, whereas the other methods need read labels, somatic variants or a model for the modification. Supplementary Section S6.5 gives further controls.

#### Key Result 4: RawMod embeddings can separate reads by origin without labels

RawMod per-read embeddings are a label-free, modification-aware *fingerprint* of read origin.

## 4. Conclusion

We introduce RawMod, a modification-agnostic method that detects DNA base modifications from raw nanopore signal without an unmodified control. One RawMod model with 0.23 million parameters detects five modification types in held-out genomic regions. It achieves a mean AUROC of 0.964, against 0.799 for the best released model. It also detects a withheld modification type when another modification of the same base remains in training. We use RawMod to map 5hmU on an *Amphidinium carterae* contig without a control. The map is consistent with two published trends and leads to a testable hypothesis that 5hmU is concentrated at divergent strand-switch regions. To promote further method development of 5hmU modification callers, we sequence and release raw nanopore signal of SPO1 and *Amphidinium carterae*. RawMod read embeddings also separate reads by origin without labels where the samples differ in modification, and they improve Dorado on SPO1, where it lacks a model for the modification.

We therefore see RawMod not only as a modification caller but also as a screen for non-standard bases. 5hmU in SPO1 and in dinoflagellates and base J in kinetoplastids replace a canonical base across a genome or large blocks of it. These organisms are rarely sequenced with an unmodified control, and no released caller has a model for their bases. Such bases even corrupt the basecall that every caller starts from. RawMod flags these bases without a model for them, as it does for 5hmU in SPO1 and *Amphidinium carterae*. Naming the base still requires an orthogonal assay, such as the mass spectrometry and chemical labeling of earlier 5hmU studies [67, 90].

RawMod also has three limitations, which Supplementary Section S7.3 discusses. Its scores are not calibrated modification fractions, it cannot name modifications, and its read-origin analysis is not yet a full method. Future work can apply RawMod to surveys of organisms with unknown epigenomes and non-standard bases, such as base J [68] in kinetoplastids, and use RawMod read embeddings for classification.

## Supporting information

Supplemental Manuscript

## Acknowledgments

We thank the STORM Research Group members for their feedback. Sequencing work was supported by grants from USDA EGP Improvement Grant, “Acquisition of Illumina NextSeq 2000 Sequencing System to Create BALANS (BioAnalytical Laboratory NextSeq) Core Supporting Salmon Aquaculture and Environmental DNA Research” 2025-2028. STORM Re-search Group acknowledges the gifts from Advanced Micro Devices (AMD) and AMD University Program’s AI & HPC Cluster.

