## Supplemental Manuscript for "RawMod: Modification-Agnostic Detection of DNA Base Modifications from Nanopore Signal Pileups"

### Supplementary Material for
### RawMod: Modification-Agnostic Detection of DNA Base Modifications from Nanopore Signal Pileups

This supplement continues the main text. Sections S1 and S2 report how RawMod generalizes to modification types withheld from training, to low coverage, and to a new modification with few labeled sites. Section S3 examines what the site embeddings encode, including modifications that no other caller can score, and Section S4 which parts of the design matter. Sections S5 and S6 hold the full analyses behind the two applications, and Section S7 discusses the design of RawMod and its limitations. The remaining sections are for reproduction. Section S8 describes the datasets, the two libraries we sequenced for this work, and the model. Section S9 describes how every tool was run and scored, and Section S10 what each one costs.

#### S1. Modification and coverage generalization

A modification-agnostic caller should detect modification types that are absent from its training data, and it should work at low read coverage. We test the first requirement by withholding one modification type from training and scoring it with a model that has not seen this modification. We test the second requirement by scoring the same sites with fewer reads per pileup, which simulates the low-coverage regions of a genome. For this, we train two configurations of RawMod. The `mixed` model is trained on all datasets, with one contiguous ∼15% region of every genome excluded for testing, as Section 3.1 describes. The leave-one-type-out model (LOCO) additionally withholds every modified site of one type and every benchmark genome that carries it. The oligonucleotides, *H. pylori* and SPO1 stay in training through their other modifications and unmodified controls, and the model is tested on the pairwise sites of that modification inside the held-out regions. Figure 4 and Table S1 show the results. We make three key observations.

First, the `mixed` model detects the modification types it was trained on. It achieves a mean AUROC of 0.988 over three training seeds (0.987–0.989) and an AUPRC of 0.962 at a positive fraction of 0.23 on the held-out regions, which hold 12,927 modified and 43,325 unmodified positions. The modification-specific callers remain more accurate on their own types, as Section 3.2 shows.

Second, withholding a modification type from training still allows RawMod to detect three of the five types. The mean zero-shot AUROC over the five LOCO folds is 0.792 over three seeds (Tables S1 and S2). 5hmC transfers best (0.983), followed by 5mC (0.978) and 4mC (0.906), whereas 6mA (0.594) and 5hmU (0.497) stay close to chance. Transfer therefore depends on whether a modification of the same base remains in training: 5mC, 5hmC and 4mC all modify cytosine, whereas 6mA and 5hmU are the only modification types on their base in our data. The `mixed` model, which is trained on all five, carries at least one modification on each of A, C and T. Guanine modifications are rare in DNA; the best known, the 7-deazaguanines, occur in some phages and bacteria [1, 2], and we do not expect them to transfer from the other bases.

Two further properties of 5hmU likely add to its poor transfer, beyond the lack of a thymine relative in training. It perturbs the signal at the modified base very little (Fig. 1b), and it replaces every thymine in SPO1, whereas every other type in our data is a sparse mark among unmodified copies of its base. The signal itself is sufficient to detect it: Section 3.4 shows that after retraining with all 10,173 labeled 5hmU sites, the model reaches an AUROC of 0.867. The 5hmU fold is therefore limited by zero-shot transfer and not by the signal. Table S3 shows how the ablations of RawMod affect zero-shot transfer.

Third, RawMod is robust to low coverage as Fig. 4b shows. When we mask reads at inference, the `mixed` model reaches an AUROC of 0.898 from a single read, 0.963 from three reads and 0.980 from six, and 0.987 with all $N$=15 reads. 5hmC and 5mC exceed 0.95 from two reads onward, whereas 5hmU improves steadily with depth, from 0.661 with one read to 0.824 with six and 0.871 with all 15.

#### S2. Zero-shot and few-shot evaluation

This section describes the configuration shared by the models of Table S1, and then the few-shot protocol that asks how many labeled sites a new modification type needs. We train the `mixed` model and the five LOCO models with one configuration and three random seeds, so that the folds differ only in the withheld modification type: (i) We train with the class-weighted binary cross-entropy, the supervised contrastive loss ($\lambda_{\text{con}}$=1, $\tau$=0.2, 128-dimensional projection) and a hinge Deep SAD loss ($\lambda_{\text{SAD}}$=1, 16 dimensions, margin 2), as defined in Section 2.3. (ii) The base-identity channels are visible to the model. (iii) In every LOCO model, we cap the training positives of the most frequent reference base at four times those of the second most frequent base, so that one base does not dominate the positives. (iv) In every LOCO model, we remove all training positives on thymine, so that no LOCO model is trained with 5hmU and incorrect basecalls cannot contaminate the positives. (v) We use two cross-read transformer layers and the curriculum of Section 2. (vi) We exclude positives that cannot be assigned a single modification, because their label is ambiguous. (vii) Each pileup holds the reads of one strand, and reverse-strand sites are built in read orientation.

6mA and 5hmU are the only modifications on their bases, so their folds need unmodified positions of the same base in the test set. For these two folds, we therefore move the control positions at the target base of the held-out organisms from training to

**Table S1: Zero-shot detection of held-out modifications.** $\pi$ **is the baseline AUPRC. Every model is trained without a contiguous ~15% region of every genome and scored inside those regions. AUROC and AUPRC are means over three training seeds, with the AUROC range.** $n_+$, $n_-$: **modified and unmodified positions inside the regions.**

| Held out | AUROC | Range (3 seeds) | AUPRC | $\pi$ | $n_+$ | $n_-$ |
|---|---|---|---|---|---|---|
| `mixed` | 0.988 | 0.987–0.989 | 0.962 | 0.23 | 12,927 | 43,325 |
| 5mC | 0.978 | 0.974–0.985 | 0.755 | 0.02 | 78 | 3,492 |
| 5hmC | 0.983 | 0.971–0.993 | 0.675 | 0.01 | 40 | 3,492 |
| 6mA | 0.594 | 0.494–0.649 | 0.287 | 0.21 | 6,274 | 24,071 |
| 4mC | 0.906 | 0.900–0.914 | 0.491 | 0.04 | 551 | 12,792 |
| 5hmU | 0.497 | 0.470–0.519 | 0.677 | 0.67 | 3,570 | 1,728 |
| *mean (LOCO)* | 0.792 | | 0.577 | | | |

testing, and we raise the cap on SPO1 control pileups to 200,000. Inside the held-out regions, the 5hmU test set contains 3,570 positive and 1,728 negative positions.

*Test sites of the zero-shot evaluation.* Table S2 lists the test sites of Table S1 by source. Every test set holds only sites inside the held-out regions, which leaves the 5mC and 5hmC folds with 78 and 40 positives, because only the oligonucleotides carry 5hmC and five of their 32 constructs are held out. The *E. coli*, *A. thaliana* and human sites inside the held-out regions are not part of Table S1. On these genomes, the model trained without 5mC detects 5mC with an AUROC of only 0.56–0.74, against 0.95 on the oligonucleotides, whereas the model trained with 5mC reaches 0.78–0.99 on the same sites. The high 5mC fold of Table S1 therefore rests mostly on the oligonucleotides. The two kinds of test site ask different questions. The 5mC fold removes every organism that carries 5mC, so a 5mC site on one of these genomes is new in both its modification and its organism, and the only genomic cytosine modification left in training is 4mC in *H. pylori* 26695, whose labels come from the motif annotation of one organism. We attribute the weak transfer on genomes to this lack of genomic training data. Transfer between modifications of the same base still holds where the training data cover the library type: the 5hmC oligonucleotide library stays in training, and the 5mC fold reaches 0.954 on the oligonucleotides. Training on more genomes that carry cytosine modifications should close this gap.

#### S2.1. Few-shot protocol

For each LOCO fold, we take labeled sites from positions of the withheld modification outside the held-out genomic regions. We split these positions into a support half and a query half by a hash of the genomic coordinate, keeping the modified and control pileups of a position together, so that the split depends neither on $K$ nor on the draw. Within the support half, we rank positions with a seeded hash, so that the $K$ positions of a draw are nested within those of a larger $K$. Every model, including $K = 0$, which is the LOCO model itself, is scored inside the held-out regions on the test set of Table S1 and not on the query half.

*Fine-tuning.* We initialize from the LOCO model and train for 8 epochs with a learning rate of $2\times10^{-4}$. The labeled pileups are repeated to make up ~25% of the training pileups (at most 50 repeats) and mixed with 30,000 replayed pileups from the fold's training set, so that the model keeps what it learned about the other modifications. We keep the final-epoch weights and select no checkpoint on labeled or query pileups, so that no label leaks into model selection; the validation set contains replayed pileups only. We use five draws for $K \leq 1{,}000$ and two above.

*Retraining.* We add the labeled pileups to the fold's training pool and train the LOCO model from scratch with the final configuration of Section S2. We use two draws for $K \leq 1{,}000$ and one above.

*Supervised reference.* The supervised reference of 0.871 in Fig. 4c and of 0.989 in Fig. 4d is the `mixed` model of Table S1, scored on the 5hmU or 6mA positions inside the held-out regions against the same negatives as the LOCO model. It was trained on every site of that modification outside the regions, about twice the support half of the few-shot runs.

### S3. Embedding analysis

This section evaluates the site embeddings of RawMod, because they explain the zero-shot results of Table S1. We embed the 56,252 positions inside the held-out regions, which form the test set of Table S1, with the `mixed` model of Table S1, and we use the 96-dimensional site embedding because it is the input of the classification head. We make three observations.

First, the embedding is organized by the label and not by the modification type. $k$-means with two clusters recovers modified against unmodified positions with an ARI of 0.75, whereas $k$-means with one cluster per modification recovers the modification of the positive positions with an ARI of only 0.26. A linear probe (logistic regression, 5-fold cross-validation) predicts the modification of the typed positive positions with 98.4% accuracy, but the reference base and the dataset alone predict it with 99.3%, because each modification sits on one base and nearly all positions of each modification come from one dataset (6mA and 4mC from *H. pylori*, 5hmU from SPO1). When we hold the base and the dataset fixed, the probe separates 4mC from 5mC on

**Table S2: Test sites of the region-holdout evaluation (Table S1) by source. Every site lies inside a held-out region. Pos., Neg.: positive and negative sites. AUROC: a source's positives against its own negatives, mean over three training seeds; "all" is the test set of Table S1. Parentheses: organisms that Table S1 does not score. For a held-out modification these organisms were removed from training entirely. The human row pools HG001 and HG002 and has only 71 negative sites from the training labels, whereas Table 1 scores HG002 at the benchmark's own sites from 10 reads per site, so the two AUROCs are not comparable.**

| Held out | Source | Pos. | Neg. | AUROC |
|---|---|---|---|---|
| `mixed` | ONT oligonucleotides | 120 | 2,702 | 0.990 |
| | SPO1 | 3,578 | 6,988 | 0.969 |
| | *H. pylori* 26695 | 9,229 | 33,635 | 0.991 |
| | *E. coli* | (11,681) | (59,025) | (0.993) |
| | *A. thaliana* | (390) | (18,945) | (0.987) |
| | HG001 + HG002 | (837) | (71) | (0.781) |
| | *all (Table S1)* | 12,927 | 43,325 | 0.988 |
| 5mC | ONT oligonucleotides | 40 | 720 | 0.954 |
| | SPO1 | 0 | 2,772 | – |
| | *H. pylori* 26695 | 38 | 0 | – |
| | *E. coli* | (10,209) | (36,563) | (0.559) |
| | *A. thaliana* | (390) | (18,945) | (0.740) |
| | HG001 + HG002 | (837) | (71) | (0.596) |
| | *all (Table S1)* | 78 | 3,492 | 0.978 |
| 5hmC | ONT oligonucleotides | 40 | 720 | 0.956 |
| | SPO1 | 0 | 2,772 | – |
| | *all (Table S1)* | 40 | 3,492 | 0.983 |
| 6mA | ONT oligonucleotides | 40 | 740 | 0.545 |
| | SPO1 | 8 | 2,488 | 0.351 |
| | *H. pylori* 26695 | 6,226 | 20,843 | 0.592 |
| | *E. coli* | (523) | (22,462) | (0.527) |
| | *all (Table S1)* | 6,274 | 24,071 | 0.594 |
| 4mC | *H. pylori* 26695 | 551 | 12,792 | 0.906 |
| | *all (Table S1)* | 551 | 12,792 | 0.906 |
| 5hmU | SPO1 | 3,570 | 1,728 | 0.497 |
| | *all (Table S1)* | 3,570 | 1,728 | 0.497 |

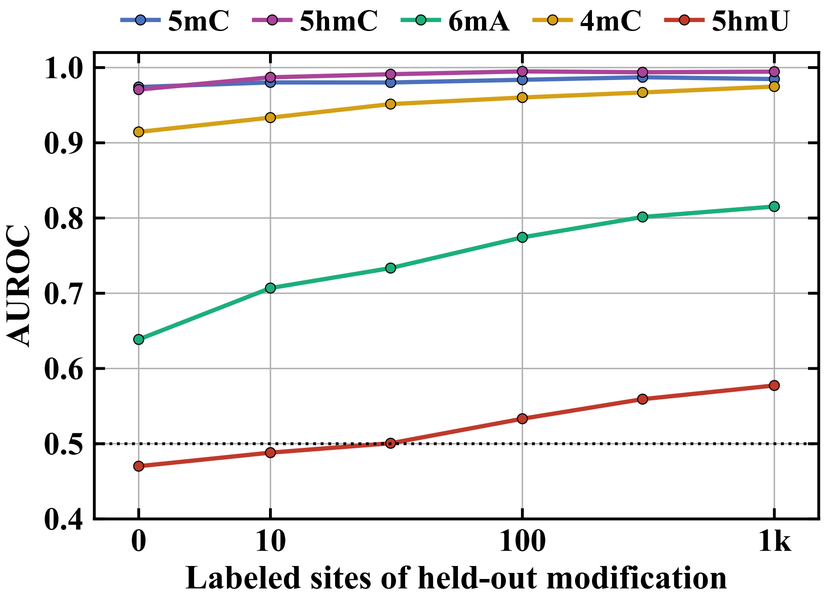


**Figure S1: Fine-tuning curves of all five LOCO models. AUROC against the number of labeled positions of the held-out modification (Section S2.1). 5mC blue, 5hmC purple, 6mA aqua, 4mC gold, 5hmU red; dotted: chance. Each curve is scored inside the held-out regions on the test set of Table S1.** $K = 0$ **is the LOCO model. Each point is the mean over two draws, and over five for 5hmU and 6mA up to 1,000 sites.**

*H. pylori* cytosines with a balanced accuracy of 0.50, and 5mC from 5hmC on the few oligonucleotide cytosines inside the regions with 0.70.

Second, the dataset is encoded even where there is no modification. Among unmodified positions, the probe predicts the dataset with a balanced accuracy of 0.87, against 0.48 from the reference base alone, although $k$-means does not recover the dataset (ARI 0.02). The zero-shot results of Table S1 do not require type-specific groups, because they score only modified against unmodified positions.

Third, the same structure holds on many more organisms (Fig. S2). We take up to 150 modified and 150 unmodified positions from each of the 15 benchmark datasets inside the held-out regions, 3,737 positions from nine organisms, each scored from the same 10-read pileups as in Table 1. Mouse brain and *O. sativa* were never in training at all.

The embedding forms one arc, ordered by the model score from unmodified to modified positions, and the positions of every organism follow the same arc. The unmethylated CpGs of human, mouse, rice and *A. thaliana* lie at the same end as the unmethylated adenines of *E. coli* and *Anabaena*, and their methylated CpGs lie at the other end beside the methylated adenines. Among the 15 nearest neighbors of a position in the 96-dimensional embedding, 91% share its label (50% by chance) and 44% its organism (14% by chance). The one exception is SPO1, whose positions have neighbors from the same organism (95%) more often than neighbors with the same label (80%) and form their own branch of the arc; they are still ranked well, with an AUROC of 0.94 in this sample. The lowest AUROC in the sample is that of the oligonucleotides (0.93).

In summary, the site embedding separates modified from unmodified positions along one axis that all organisms share, it carries the dataset of origin, and it does not arrange the modifications into type-specific groups.

#### S3.1. Modifications that other callers cannot score

A modification-specific caller reports only the modifications it has a model for, and every other modified site is invisible to it. Figure S3a marks, on the embedding of Fig. S2, the modified positions for which each released single-sample caller has a model. Dorado covers 92% of these positions, UniMeth 82%, MethyNano 55%, and DeepMod2 and Rockfish, which call 5mC at CpG only, 43%. RawMod scores all of them, and 5hmU is the only modification that no other caller can score. These shares refer to the sampled positions, not to a genome.

We repeat the comparison on a genome without ground truth, contig_3762 of *Amphidinium carterae* (Fig. S3b). Dorado calls the reads with its all-context 6mA, 5mC+5hmC and 4mC+5mC models, and we keep forward-strand sites with at least 10 reads. We count a site as modified when at least half of its reads are called modified, and as unmodified when at most 10% are. We sample 4,578 sites from these classes and from the T and G sites, for which Dorado has no model, and embed them with the RawMod model trained on all data. The embedding splits into a high-score and a low-score lobe. 1,007 of the 1,482 T sites, where no other caller has a model, score at least 0.5; this is the thymine signal of Section 3.5. The CpG 5mC calls of Dorado also score high (mean 0.85), whereas its 5hmC and non-CpG 5mC calls lie between the lobes (0.56 and 0.54).

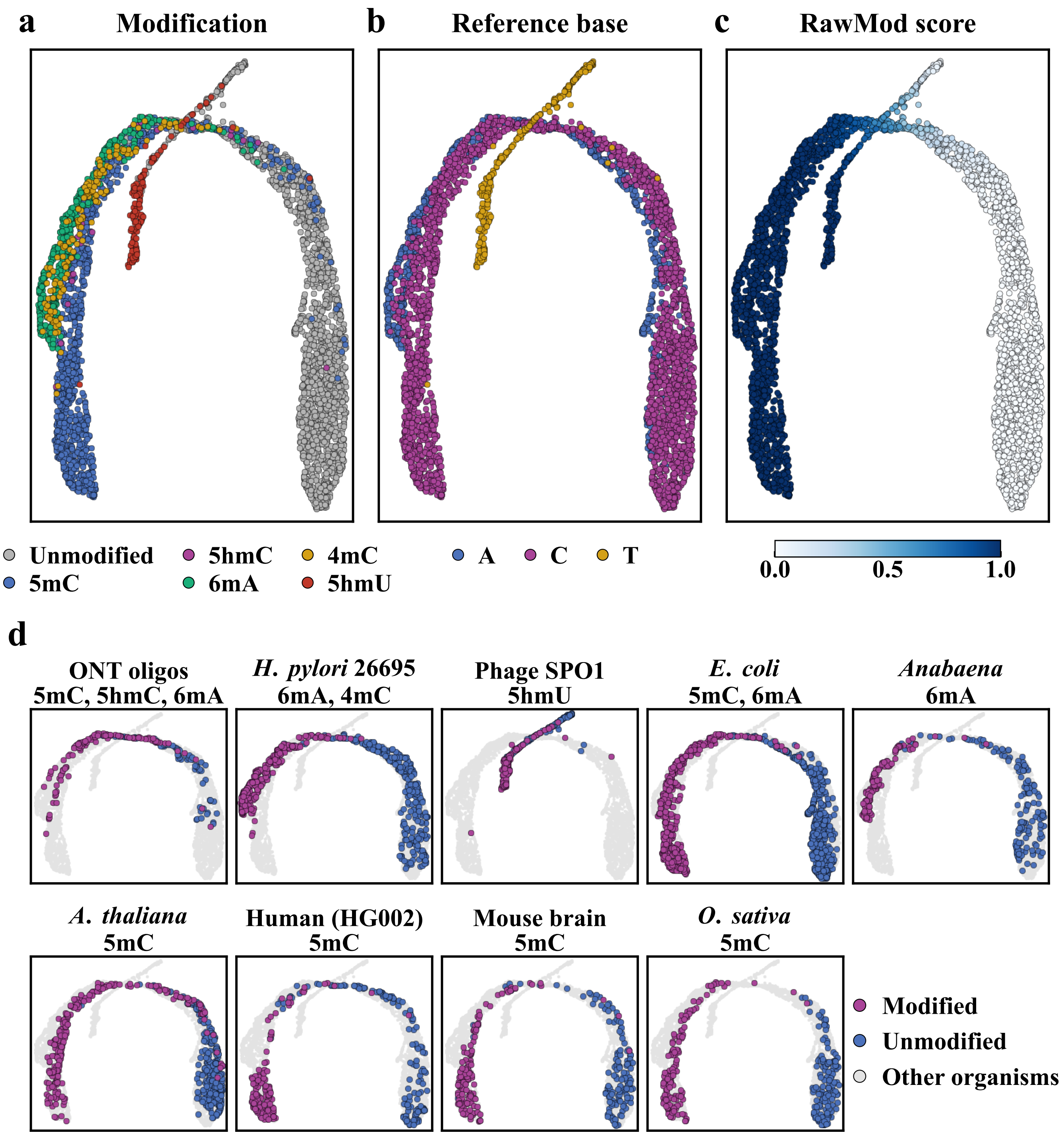


**Figure S2:** UMAP of the site embeddings of the final `mixed` model at 3,737 benchmark positions from nine organisms. The positions lie inside the held-out regions and are colored by (a) modification (unmodified grey, 5mC blue, 5hmC purple, 6mA aqua, 4mC gold, 5hmU red), (b) reference base (A blue, C purple, T gold), (c) RawMod score (light to dark blue) and (d) organism (modified purple, unmodified blue, all others grey). The embedding is the 96-dimensional site embedding. At most 150 modified and 150 unmodified positions are drawn per dataset, all at positions the model never trained on. Panel (d) shows the positions of each organism over all other positions, with the modifications it contributes.

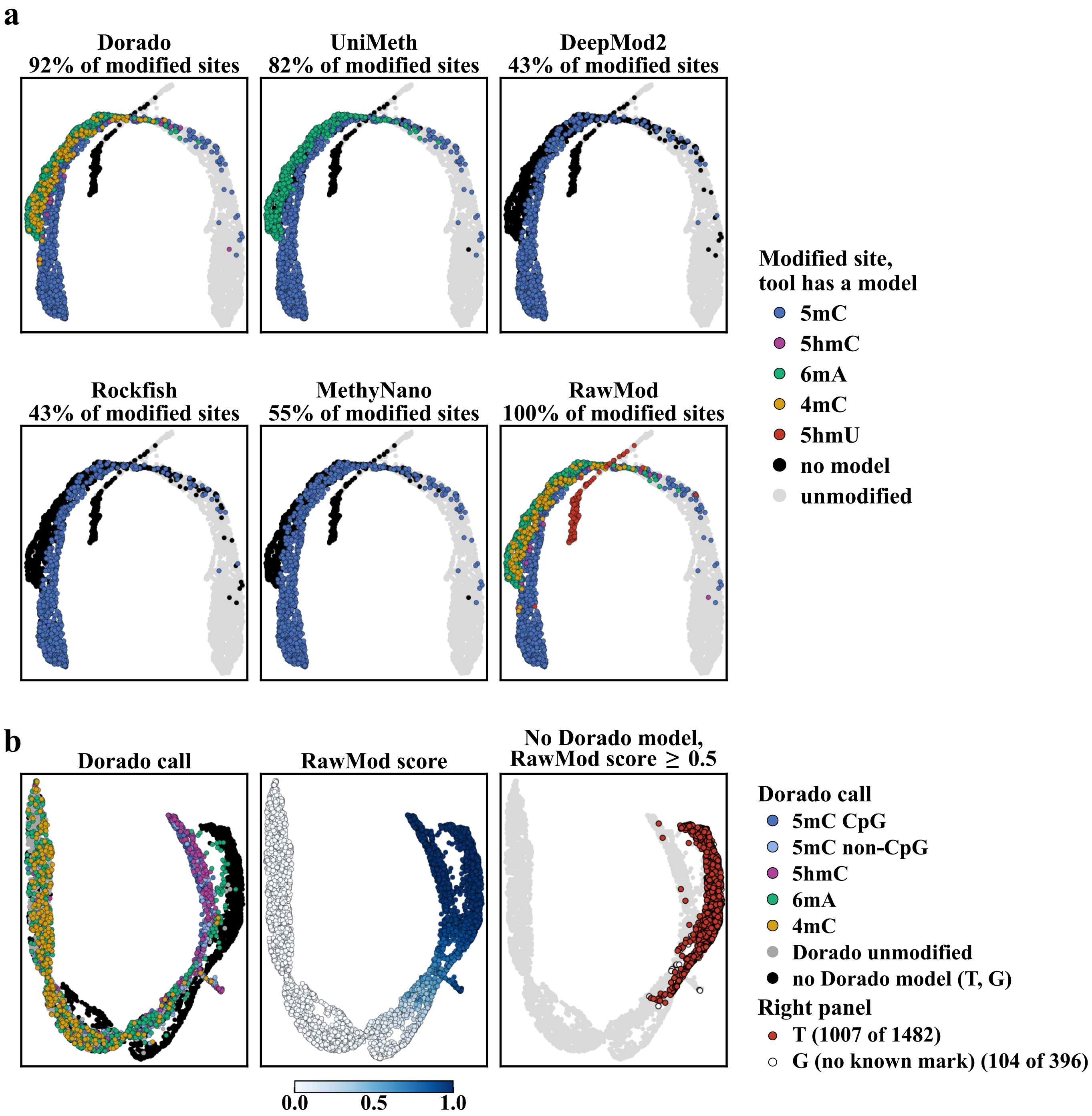


**Figure S3: Modified sites that each caller can score. (a) The embedding of Fig. S2, once per released single-sample caller. Modified positions are colored by modification where the caller has a model for it and black where it has none; unmodified positions are grey. Titles give the share of modified positions with a model. (b) 4,578 forward-strand sites of *Amphidinium carterae* contig_3762, embedded with the model trained on all data. Left: Dorado call; black: T and G sites, for which Dorado has no model. Middle: RawMod score. Right: sites without a Dorado model that score at least 0.5 (T red, G open).**

#### S4. Ablations

##### S4.1. Component ablations

Table S3 evaluates the contribution of RawMod's input features, architecture, and training choices by changing one component of the final configuration at a time. Across three training seeds, the final model achieves a mean zero-shot AUROC of 0.777–0.804, providing a reference for run-to-run variation. The largest effect comes from removing the site-deviation channel, which reduces the `mixed` AUROC from 0.987–0.989 to 0.913–0.916 across three seeds. Zero-shot accuracy also decreases for 5mC, 5hmC, and 4mC, although the effects vary across modification types. By contrast, 6mA improves when the channel is removed, and 5hmU shows no consistent decrease. The downward trend for both the mixed model and the majority of LOCOs show that explicitly representing the difference between observed and expected signals substantially improves modification detection, but does not benefit every modification type.

Other changes have smaller effects. Removing the cross-read transformer, learned row embedding, or base identity channel produces relatively small changes in overall accuracy, generally within the variation observed across training seeds. Similarly, removing the curriculum has little effect on mean zero-shot AUROC and does not improve library invariance: at the background positions of Table S15a, mean invariance is 0.92–0.94 without the curriculum and 0.84–0.93 with it. We keep it for its small gain on 4mC. These results suggest that RawMod is relatively robust to individual architectural and training choices, with the site-deviation channel providing the clearest contribution to detection accuracy.

We examine the contribution of the auxiliary training losses separately in Section S4.2.

**Table S3: Comparisons of RawMod architectures and training objectives. Zero-shot AUROC of each LOCO fold and of the `mixed` model, scored inside the held-out regions. Every row changes one component of the final configuration (seed 42). Seeds 43 and 44 give the run-to-run spread of select experiments.**

| Configuration | `mixed` | 5mC | 5hmC | 6mA | 4mC | 5hmU | Mean |
|---|---|---|---|---|---|---|---|
| RawMod (final, seed 42) | 0.987 | 0.974 | 0.971 | 0.639 | 0.914 | 0.470 | 0.794 |
| seed 43 | 0.989 | 0.985 | 0.984 | 0.494 | 0.904 | 0.519 | 0.777 |
| seed 44 | 0.989 | 0.976 | 0.993 | 0.649 | 0.900 | 0.502 | 0.804 |
| *Input features* | | | | | | | |
| − site-deviation channel | 0.913 | 0.970 | 0.939 | 0.692 | 0.807 | 0.483 | 0.778 |
| seed 43 | 0.916 | 0.962 | 0.919 | 0.656 | 0.790 | 0.510 | 0.768 |
| seed 44 | 0.914 | 0.958 | 0.896 | 0.662 | 0.816 | 0.502 | 0.767 |
| Base identity masked | 0.984 | 0.974 | 0.983 | 0.606 | 0.859 | 0.507 | 0.786 |
| *Architecture* | | | | | | | |
| No cross-read Transformer (mean-pooled read tokens) | 0.979 | 0.962 | 0.970 | 0.636 | 0.881 | 0.477 | 0.785 |
| 1 Transformer layer | 0.988 | 0.977 | 0.983 | 0.684 | 0.899 | 0.500 | 0.809 |
| 4 Transformer layers | 0.989 | 0.970 | 0.976 | 0.583 | 0.898 | 0.492 | 0.784 |
| − learned row embedding | 0.988 | 0.967 | 0.984 | 0.602 | 0.884 | 0.515 | 0.790 |
| *Training objective* | | | | | | | |
| − supervised contrastive loss | 0.989 | 0.981 | 0.977 | 0.676 | 0.921 | 0.504 | 0.812 |
| − Deep SAD loss | 0.989 | 0.976 | 0.977 | 0.599 | 0.904 | 0.469 | 0.785 |
| − both (classification loss only) | 0.989 | 0.978 | 0.988 | 0.682 | 0.922 | 0.556 | 0.825 |
| seed 43 | 0.988 | 0.972 | 0.976 | 0.519 | 0.901 | 0.500 | 0.774 |
| seed 44 | 0.989 | 0.977 | 0.963 | 0.573 | 0.889 | 0.535 | 0.787 |
| Deep SAD width 64 | 0.989 | 0.984 | 0.987 | 0.607 | 0.928 | 0.474 | 0.796 |
| *Training data* | | | | | | | |
| − curriculum (no matched-only stage) | 0.989 | 0.983 | 0.988 | 0.567 | 0.887 | 0.506 | 0.786 |
| seed 43 | 0.989 | 0.967 | 0.988 | 0.628 | 0.894 | 0.485 | 0.792 |
| seed 44 | 0.989 | 0.980 | 0.985 | 0.643 | 0.882 | 0.487 | 0.795 |

Every model here is trained and scored with the held-out regions of Section S9.7, so the zero-shot results and the benchmark of Table 1 share one split.

##### S4.2. Auxiliary losses

Table S3 shows that training with classification loss alone achieves comparable zero-shot detection accuracy to the final model across three seeds (mean AUROC 0.774–0.825 versus 0.777–0.804). We therefore examine whether the supervised contrastive and Deep SAD losses improve the learned embeddings beyond detection accuracy. Table S4 compares the final configuration with the classification-only model on three tasks: site-embedding separation, zero-shot detection, and label-free read-origin typing.

First, we evaluate how well the site embeddings separate modified and unmodified positions (Table S4a). For each dataset and reference base with both classes in the `mixed` test set, we cluster the 96-dimensional site embeddings into two groups using $k$-means. We report the ARI of these clusters against the true labels and the silhouette of the labeled classes. The final model separates the classes better in four of the five groups, with SPO1 as the exception (ARI 0.40 versus 0.45). Pooled over the groups, the final model improves ARI by 0.11 (bootstrap 95% CI 0.10–0.13).

Second, we evaluate zero-shot detection using the classifier logits of the LOCO models (Table S4b). In the reported seed, the classification-only model performs better on 5hmC, 6mA, and 5hmU. However, across three seeds, the two configurations achieve comparable mean zero-shot AUROC (0.777–0.804 versus 0.774–0.825, Table S3).

Third, we evaluate label-free read-origin typing using the read embeddings of both configurations on the windows of Section 3.6. On COLO829, the median ARI is 0.21 for the final model and 0.29 for classification only, with the final model ahead in 14 of 32 windows (Wilcoxon $p = 0.72$). On HS578, both achieve a median ARI of 0.02 ($p = 0.08$). The median oriented AUROC is 0.77 versus 0.84 on COLO829 ($p = 0.06$) and 0.69 versus 0.72 on HS578 ($p = 0.13$).

Overall, the auxiliary losses improve the separation of modified and unmodified sites in the embedding space but do not improve zero-shot detection or label-free read-origin typing. We retain them because they produce more clearly separated site embeddings without a demonstrated loss in detection accuracy.

**Table S4: Auxiliary losses against classification only. (a) Separation of modified and unmodified positions in the site embedding, without labels. (b) Zero-shot AUROC of the withheld modification.**

**(a) Site embedding of the `mixed` model, positions inside the withheld regions**

| | | | *k*-means ARI | | Silhouette | |
|---|---|---|---|---|---|---|
| **Dataset, base** | **Mod.** | **Unmod.** | Final | Class. only | Final | Class. only |
| *H. pylori* 26695, A | 6,226 | 20,843 | 0.747 | 0.571 | 0.672 | 0.573 |
| *H. pylori* 26695, C | 3,003 | 12,792 | 0.880 | 0.836 | 0.796 | 0.700 |
| Oligonucleotides, A | 40 | 740 | 0.336 | 0.100 | 0.579 | 0.447 |
| Oligonucleotides, C | 80 | 720 | 0.776 | 0.481 | 0.710 | 0.607 |
| SPO1, T | 3,570 | 1,728 | 0.402 | 0.448 | 0.353 | 0.354 |

**(b) Withheld modification (LOCO models), zero-shot AUROC of the classifier logit**

| **Objective** | **5mC** | **5hmC** | **4mC** | **6mA** | **5hmU** |
|---|---|---|---|---|---|
| Final | 0.974 | 0.971 | 0.914 | 0.639 | 0.470 |
| Classification only | 0.978 | **0.988** | 0.922 | **0.682** | **0.556** |

#### S4.3. No-learning baselines

Table S5 scores the test sets of Table S1 with fixed statistics of the pileup instead of the model: the mean over reads of the absolute and signed deviation of the current from the expected level at the site's base (the model's site-deviation channel), the mean absolute deviation over the 21-base window, and the fraction of reads on one side of the expected level. Pileups are averaged per site as for the model. No statistic exceeds 0.80 on any fold. Values below 0.5 are statistics that run the other way.

**Table S5: No-learning baselines on the region-holdout test sets. AUROC of fixed statistics of the signal against the model. $n_+$, $n_-$: positions.**

| **Model** | $n_+$ | $n_-$ | **RawMod** | **$\|\Delta\|$ at base** | **$\Delta$ at base** | **window $\|\Delta\|$** | **side fraction** |
|---|---|---|---|---|---|---|---|
| `mixed` (all modifications) | 12,927 | 43,325 | 0.987 | 0.668 | 0.454 | 0.568 | 0.601 |
| 5mC held out | 78 | 3,492 | 0.974 | 0.598 | 0.388 | 0.512 | 0.486 |
| 5hmC held out | 40 | 3,492 | 0.971 | 0.795 | 0.213 | 0.550 | 0.508 |
| 6mA held out | 6,274 | 24,071 | 0.639 | 0.551 | 0.650 | 0.513 | 0.592 |
| 4mC held out | 551 | 12,792 | 0.914 | 0.562 | 0.460 | 0.562 | 0.421 |
| 5hmU held out | 3,570 | 1,728 | 0.470 | 0.479 | 0.501 | 0.430 | 0.601 |

#### S4.4. Site score and modification frequency

RawMod is trained to distinguish modified from unmodified sites, not to estimate the fraction of molecules modified at a position. To assess whether its site scores represent calibrated probabilities, Fig. S4 compares the scores with the observed fraction of modified positions across 56,252 sites in the held-out regions.

The mean predicted score exceeds the observed fraction of modified positions in every score bin, with a difference of up to 0.41 in the intermediate bins. The expected calibration error is 0.058 and the Brier score is 0.050. This overestimation is consistent with the class-weighted training loss, which assigns greater weight to modified sites. Despite this offset, RawMod accurately ranks modified above unmodified sites (AUROC 0.987). Calibration error also varies by modification type, from 0.019 for 5mC to 0.164 for 5hmU.

Thus, RawMod's site scores are useful for ranking modification likelihood but should not be interpreted directly as modification frequencies. Estimating modification fractions would require additional calibration against quantitative modification measurements.

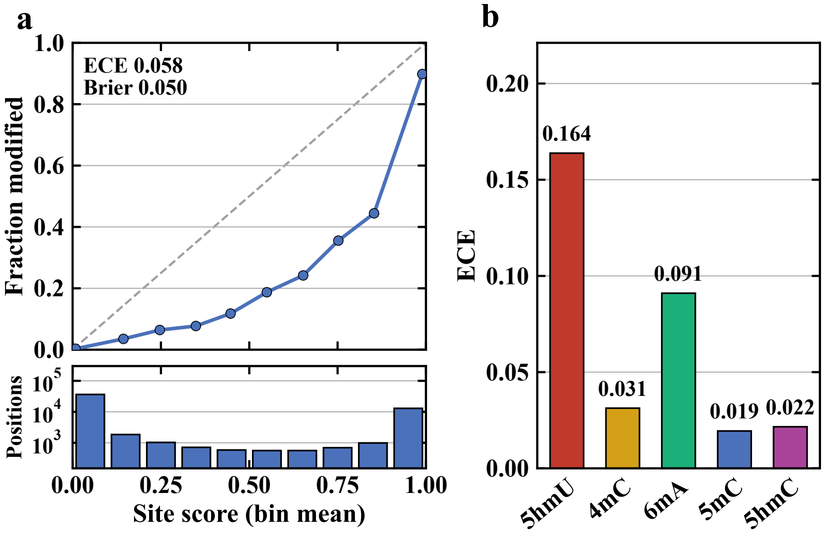


**Figure S4: The site score ranks sites and is not a modification frequency. Site score of the `mixed` model against the fraction of modified positions, on the held-out regions. (a) Fraction of modified positions against the mean site score in ten equal-width bins; dashed: score equal to the fraction; bottom: positions per bin. (b) Expected calibration error, the mean gap between score and fraction, per modification, each against the negatives of its organisms and base.**

#### S5. Additional *Amphidinium carterae* analyses

This section gives the full comparison table behind the *Amphidinium carterae* map of Section 3.5 and the checks that bound its claims. Because the map has no ground truth, we test three things: which comparisons hold under the model trained on all data, whether sequence composition explains them, and whether the published assays corroborate them.

Table S6 lists every comparison of Section 3.5 under the model trained on all data, together with the published finding that each one tests. We make three observations. First, two of the published findings hold on our map: the depletion of 5hmU in coding sequence, and the decline of the score with repeat divergence. The elevation of the divergent strand-switch regions is new, since the published enrichment lies where gene arrays converge. Second, two comparisons are not significant, the depletion at gene starts ($z = -0.6$) and the enrichment of all repeats over unique DNA ($z = -1.3$), so we count them neither for nor against the published findings. Third, exons score 1.12× higher than their ±1 kb flanks, which is the opposite of the published depletion. Exons are also much richer in GC than their flanks (0.53 against 0.39 within 50 bp). After the 5-mer and GC correction of Table S7, exons score 0.97× their flanks ($z = -4.3$), which is the published direction. Figure S6 shows the score across gene bodies, by repeat copy number, against the read error rate, and around the strand-switch regions.

**Table S6: Every comparison of Section 3.5 under the model trained on all data, with the published finding it tests. $z$: circular-shift null (score track rotated against the annotation by ≥200 kb, 1,000 rotations). Percentiles of strand-switch regions are among the 102 intergenic gaps within gene arrays.**

| Comparison | Published finding | RawMod |
|---|---|---|
| Intergenic vs. CDS | exons depleted [3] | 1.06× ($z$=3.0) |
| composition-adjusted | | 1.12–1.17× |
| Introns vs. CDS | – | 1.00× ($z = -0.1$) |
| Repeats (≤5% div.) vs. unique; score vs. divergence | younger TEs carry more 5hmU [3] | 0.773 vs. 0.680; $\rho = -0.10$ ($z = -8.6$) |
| All interspersed repeats vs. unique | TEs enriched [3] | $z = -1.3$ |
| Exons vs. ±1 kb flanks | depleted [3] | 1.122× ($z$=12.2) |
| Gene start ±1 kb vs. 1–5 kb | depleted (*B. minutum*) [4] | 0.995× ($z = -0.6$) |
| Divergent strand switches: mean (percentile) | – | 0.93 (100), 0.85 (99) |
| Convergent strand switches: mean (percentile) | boundaries (*B. minutum*) [4] | 0.77 (89), 0.63 (49) |
| 10 kb window score vs. GC | – | $\rho$=0.48 |

Sequence composition could also drive these comparisons. The 10 kb window scores follow local GC content ($\rho = 0.48$). Controlling for composition widens the intergenic/CDS contrast (Table S7). Short-read mappability within the contig does not

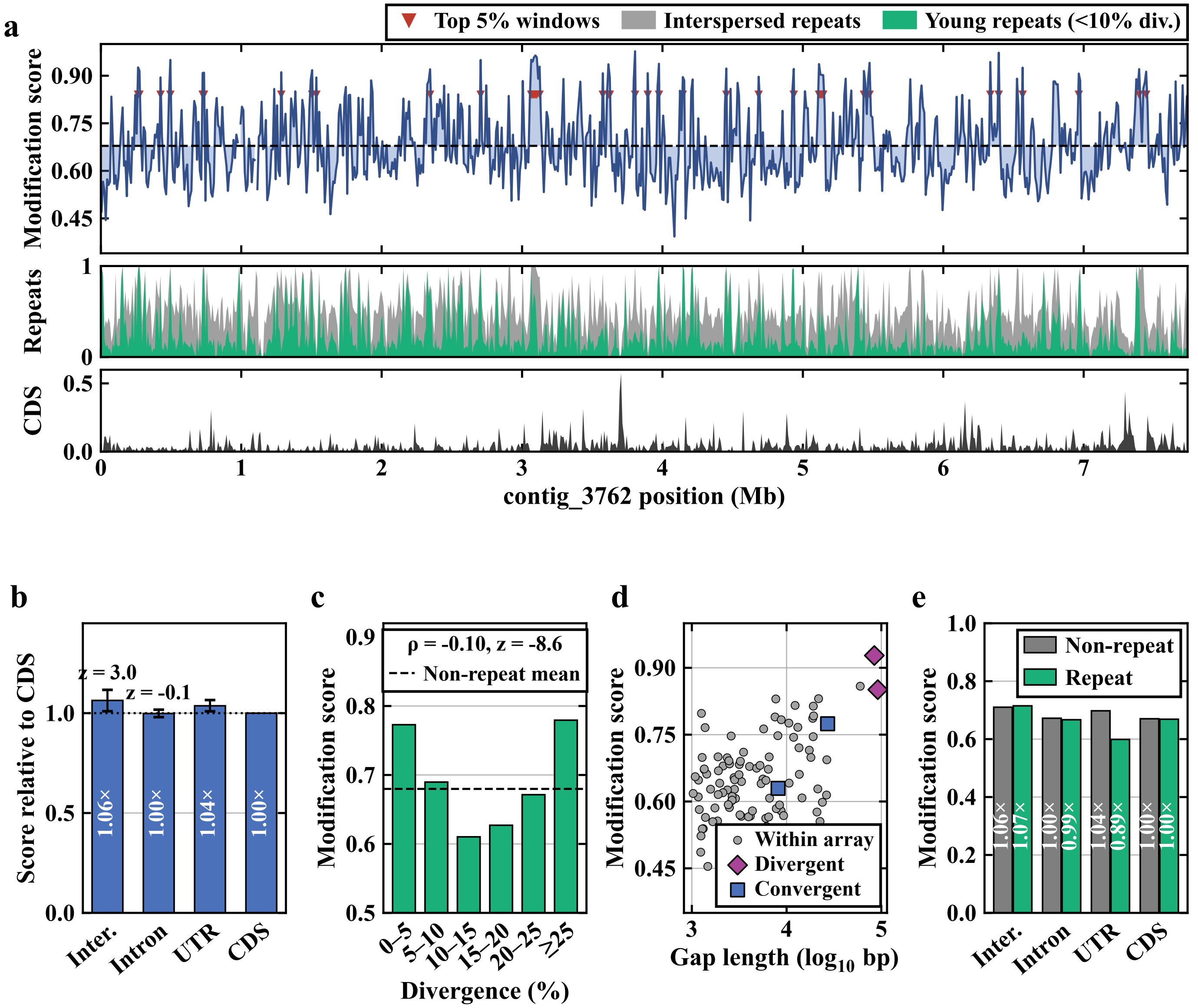


**Figure S5: Control-free 5hmU landscape of *Amphidinium carterae* contig_3762. (a) Mean thymine score in 10 kb windows, with interspersed-repeat (gray), young-repeat (aqua) and CDS (black) fractions. (b) Mean 5hmU score by annotation class relative to CDS. (c) Repeat thymines by divergence from their consensus. (d) Intergenic regions against length. (e) Repeat (aqua) and non-repeat (grey) thymines by class. Scores are from the model trained on all data at forward-strand thymines. Young repeats in (a) have <10% divergence from their family consensus; the young-repeat comparison of Section 3.5 uses copies within 5%. (b) Block-bootstrap 95% confidence intervals and circular-shift $z$. (c) Dashed line: mean of non-repeat thymines.**

**Table S7: Controlling for composition widens the intergenic/CDS contrast.**

| | intergenic | intron | UTR | CDS | IG/CDS | $R^2$ of context |
|---|---|---|---|---|---|---|
| observed | 0.713 | 0.669 | 0.695 | 0.670 | 1.06 | – |
| 5-mer residualised | 0.704 | 0.672 | 0.656 | 0.629 | 1.12 | 0.077 |
| local GC residualised | 0.707 | 0.673 | 0.641 | 0.607 | 1.16 | 0.022 |
| 5-mer × GC residualised | 0.703 | 0.674 | 0.632 | 0.600 | 1.17 | 0.095 |

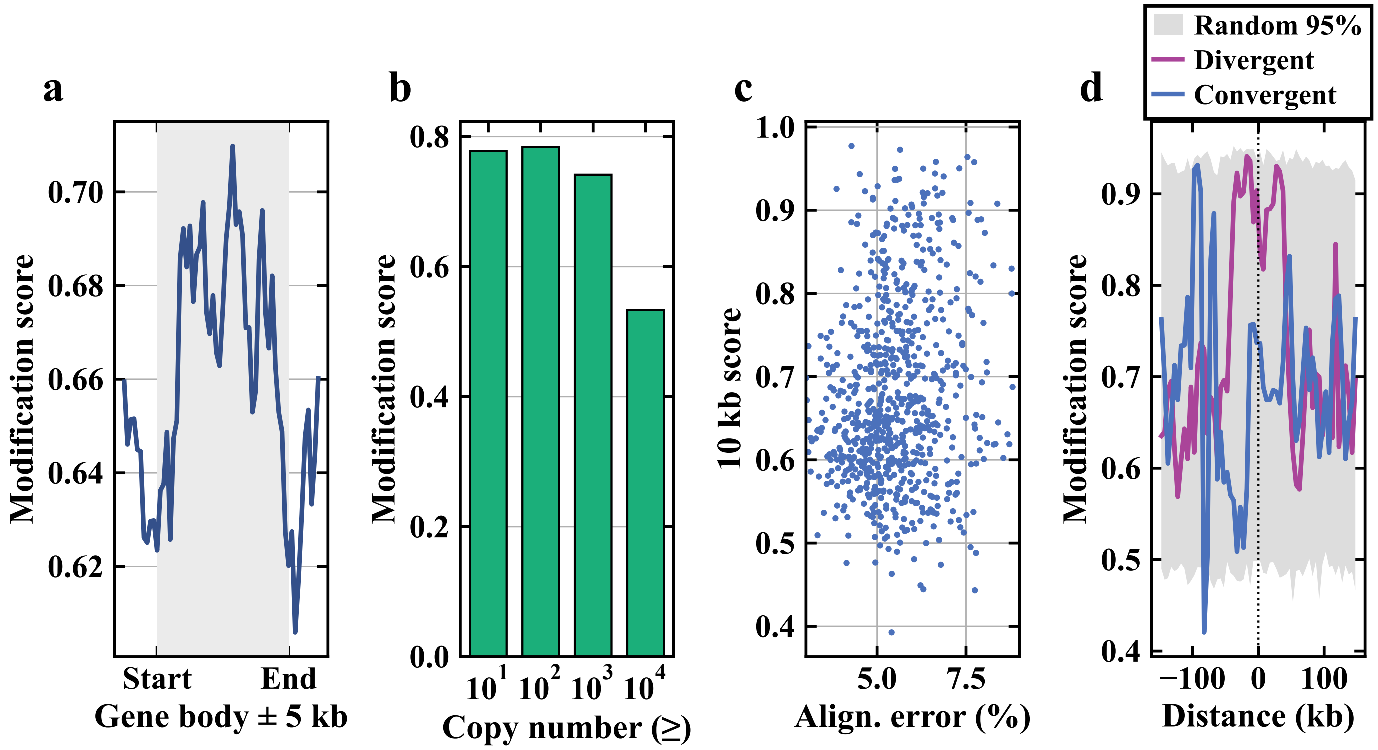


**Figure S6: Further views of the *Amphidinium carterae* 5hmU map. (a) Mean score across gene bodies and 5 kb flanks. (b) Mean score of repeat Ts by the genome-wide copy number of their RepeatModeler family. (c) 10 kb window scores against the mean alignment error rate of reads. (d) Mean score around the midpoints of divergent and convergent strand-switch regions. Divergent regions are purple and convergent regions blue; the grey band is the 95% envelope of random positions.**

explain the divergent regions. 87% and 98% of their 100-mers occur only once in contig_3762, against 94% for the whole contig. These numbers ignore copies elsewhere in the 1.25 Gb genome, which can only lower them.

We run two further checks on the map. First, we score the contig with a model of the same configuration that was trained without 15% of every genome. This model keeps the excess at the divergent junctions (100th and 98th percentiles) and the young-repeat effect. Second, we compare the map with the 5hmU-DIP and chemical pull-down profiles of the published study [3], lifted onto the contig. This comparison is weak for three reasons. The published strain differs from ours, only 69% of the contig aligns to it, and the two published assays agree poorly with each other on this contig ($\rho = 0.14$ at 1 kb). After we control for GC content, CpG density and TE density, the antibody profile is weakly anti-correlated with our scores at 1 kb (partial $\rho = -0.19$). It is also anti-correlated per TE ($\rho = -0.27$, without that control) and uncorrelated at 10 kb. The chemical profile is flat. The weak anti-correlation therefore cannot confirm the map, and the disagreement between the two published assays limits how far it can refute it.

In summary, the excess at the divergent junctions and the young-repeat effect hold under both models. Confirming the map still requires an orthogonal 5hmU assay on the sequenced strain.

#### S6. Tumor read separation

This section describes the data, the partitioning procedure and the control experiments behind the read-origin results of Section 3.6. We test whether the per-read embeddings of RawMod can split the reads of a window by sample without read labels or modification calls. We make four key observations. First, the label-free partitions follow the sample. When we orient them with methylation measured on independent flow cells, they are correct in 103 of 108 window–direction pairs, and every method stays at chance when it is run on the two flow cells of one line. Second, somatic variants cannot orient the partitions, because only 5 of the 54 windows contain one. Third, the CpG calls of Dorado separate the human cell lines at least as well as the embeddings of RawMod, whereas RawMod is ahead on SPO1 (0.796 against 0.688), where no released model detects 5hmU. Fourth, the supervised classifiers transfer when the two samples differ by a consistent tumor-versus-normal methylation direction, but not when they differ by cell type.

*Notes to Fig. 5.* COLO829 contrasts two cell types (melanoma and lymphoblastoid). HS578 contrasts a breast tumor line with a normal breast line. In Fig. 5e–h, blue methods use no read labels on these data. Of the supervised (purple) methods, MethylBERT is trained on held-out folds of these reads, and ROCIT on another sample (HG008) with no labels from ours. The

Dorado modification model differs between panels. The human panels use the CpG-only caller, and the SPO1 panel uses the all-context cytosine model. The expected oriented AUROC of random scores is drawn on every panel, as Section S6.5 defines it. The library-identity control for SPO1 is reported in the text rather than drawn.

#### S6.1. Data and windows

*COLO829.* We use R10.4.1 reads of COLO829 and COLO829BL from two flow cells per sample (tumor: PAU59949 and PAU61426; normal: PAU59807 and PAU61427). All windows are 10 kb long and start at a multiple of 10 kb. To score a partition, we need to know which CpGs differ between the samples. A CpG counts as differentially methylated when its bulk methylation (`modkit pileup`, CpG context, 5mC and 5hmC combined) differs by at least 50 percentage points between the two samples. On COLO829, the bulk values pool both flow cells of each sample, and a CpG needs at least 25 valid calls in both samples.

The 54 COLO829 windows on chr20 come in two sets. The first 24 are drawn at random among windows with at least 20 CpGs covered by 25 or more reads in both samples, without reference to methylation. Only two of them had fewer than five differential CpGs. We therefore added 30 more windows, so that the windows span a range of differential methylation: 20 from the 5% of chr20 windows with the most CpGs (at least 263 per 10 kb) and 10 at random. The tiers of <5, 5–9, 10–19, 20–49 and ≥50 differential CpGs then hold 5, 5, 11, 25 and 8 windows. A window holds a median of 40 reads in the first set and 48.5 in the second. The 54 windows hold 8,852 CpGs that pass these filters, and 1,564 of them (18%) are differential. A window holds between 0 and 140 differential CpGs, with a median of 22.5.

*HS578.* We repeat the analysis on HS578, which pairs the breast carcinoma line HS578T with the normal breast line HS578Bst [5, 6]. The two lines are reported to come from the same patient. We compare their germline genotypes. We called germline variants on chr20 with DeepVariant at a matched depth of about 10× per flow cell. First, HS578T and HS578Bst share 56–57% of their variants, and lines of HS578 and COLO829 share 54–60%. In contrast, two flow cells of one line share 87–90%, and COLO829 and COLO829BL, which come from one patient, share 87–94%. Second, at 15% of the sites where one line is confidently homozygous for the alternate allele, the other line has no read with that allele. This rate is 16–18% between lines of HS578 and COLO829 and 0% between COLO829 and COLO829BL (0 of 19,913 sites). We therefore do not call somatic variants on this pair, and we use HS578 only for methylation-based separation. The 100 HS578 windows of 10 kb on chr1 are a stratified random draw (seed 0) of 20 windows per tier of differential CpGs. We draw them among the windows with at least 10 reads in each sample and at least 40 CpGs with 8 or more valid calls in both samples. The windows hold 9,795 such CpGs, and 2,500 of them (26%) are differential. A window holds a median of 26 forward-strand reads. We use forward-strand reads only, because pooling both strands makes the consensus recover the strand rather than the sample.

*SPO1.* The windows come from the five PCR amplicon libraries sequenced with the native DNA. Each library copies a different stretch of the genome. We take each stretch where coverage reaches 1,000 reads, trim 400 bp of primer and coverage ramp from each end, and tile the rest into 21 windows of 1.2 kb. Each window compares 6,000 reads of one PCR library with 6,000 native reads. The native and amplicon arms also differ in library preparation and read length. Their read-length distributions differ enough that the consensus can partition on coverage alone. We therefore compute every SPO1 score on coverage-matched read subsets, five draws per window.

*Basecalling and modification models.* Every dataset is basecalled with Dorado 0.9.2 and the `dna_r10.4.1_e8.2_400bps_sup@v5.0.0` model. Modification calls on COLO829 and HS578 use `5mCG_5hmCG@v3`, which scores cytosine in CpG context and nothing else. This is a property of our invocation rather than a limit of Dorado, which also ships all-context and adenine models. No 5hmU model exists, so on SPO1 we use the all-context `5mC_5hmC@v3` model. On SPO1, we run it with `–modified-bases-threshold 0`, so that low-probability calls are kept rather than being indistinguishable from uncovered positions.

#### S6.2. Label-free partitioning

The partition must split the reads of a window by sample without any labels. We therefore split the reads at each site and then combine the splits over the window. For every forward-strand CpG of a window that is covered by at least 12 reads (10 on HS578), we standardize the 96-dimensional per-read embeddings. We split the reads by the sign of their first principal component. A window needs at least three such sites. We then combine the per-site splits into one partition per window. For every pair of reads, the co-association $C_{ij}$ is the mean of $s_i s_j$ over the sites that both reads cover, where $s = \pm 1$ is the side of a read at a site. We double-center $C$: we subtract its row and column means and add back the grand mean, so that the leading eigenvector is not the constant vector. The leading eigenvector is the read score, and its sign splits the reads into two groups. ARI is computed on this partition and AUROC on the continuous score.

Since the sign of the eigenvector is arbitrary, the ARI and the undirected AUROC cannot distinguish a partition by sample from a partition by flow cell. The partition also shows only that two populations differ, not which modification differs, as Section S3 shows. Panels (a, b) of Fig. 5 are a display only. There, each read's per-site embeddings are concatenated over the window (reads covering at least 60% of its sites, missing sites filled with the column mean) and projected on their first two principal components. Table S8 gives the ARI of each method and site class on the 54 COLO829 windows, by tier of differential CpGs.

**Table S8: Separation by site class with different methods. RawMod row: model trained on all data.**

| | | Differential CpGs | | | | |
|---|---|---|---|---|---|---|
| **Features** | **ARI** | <5 | 5–9 | 10–19 | 20–49 | ≥50 |
| *Windows* | 54 | 5 | 5 | 11 | 25 | 8 |
| RawMod, CpG | 0.207 | −0.002 | −0.003 | 0.133 | 0.491 | 0.858 |
| Dorado 5mC/5hmC, CpG | **0.900** | 0.002 | **0.357** | **0.805** | **0.929** | **0.958** |
| Dorado 5mC/5hmC, non-CpG C | 0.010 | 0.001 | −0.007 | −0.003 | 0.006 | 0.903 |
| Dorado 6mA, A | 0.024 | **0.012** | 0.005 | 0.019 | 0.024 | 0.799 |

#### S6.3. Orienting the partitions

The sign of a partition is arbitrary, so an independent signal must tell which side holds the tumor reads. Somatic variants are the natural choice, but they tag few reads. We call somatic variants with DeepSomatic 1.9.0 [5] (`–model_type=ONT`, paired tumor and normal) on chr20 of the published COLO829 and COLO829BL alignments (both flow cells per sample) against GRCh38. We keep the PASS single-nucleotide calls: 1,110 somatic SNVs (17.2 per Mb). A 10 kb window therefore contains a somatic SNV with a probability of only 16%. In our windows, 57 of 1,239 COLO829 reads (4.6%) carry a somatic alternate allele, and no COLO829BL read does. Only 5 of the 54 windows contain a somatic SNV, so the variants cannot orient most partitions. In one of these five, the side of the partition that holds most of the alternate-allele reads is clearly tumor-dominated (72% tumor reads). In the other four, the alternate-allele reads are split between the sides, or that side is at most 56% tumor.

We therefore orient the partitions by methylation measured on held-out flow cells. In each window, we take the CpGs whose methylation differs by at least 25 percentage points between the tumor and normal reads of one flow-cell pair (at least 10 valid calls, at least 5 such CpGs per window). We score each read of the other pair by the mean of $\mathrm{sign}(\Delta)\, P(\text{5mC})$ over the CpGs it covers, using Dorado's per-read calls. We orient the consensus score by its correlation with this read score and compute a directed AUROC. Run in both directions over all 54 windows, the orientation is correct in 103 of 108 window–direction pairs ($p = 4{\times}10^{-25}$, one-sided binomial). With the signs of $\Delta$ shuffled, it is correct in 133 of 270 (five shuffles, one direction). The label-free partitions therefore follow the sample, as measured on independent flow cells.

*Notes to Table 2.* RawMod and Dorado use no read labels; MethylBERT is trained on held-out folds of these reads and ROCIT on another sample (HG008). A dash marks a dataset where the method cannot be run (no 5hmU model, no read labels, or no somatic calls and cell-type atlas). Margins over the null and window counts: Table S9.

#### S6.4. Methods compared

We compare RawMod with Dorado, which also uses no read labels, and with two supervised classifiers, MethylBERT and ROCIT. Every comparison is paired, so that differences come from the methods and not from the reads.

*RawMod model.* On the human cell lines, every RawMod arm in Fig. 5 and Table 2 uses the model trained on all data. On SPO1, we use the region-holdout model instead, which is trained on these barcodes with every read that overlaps 52.6–72.5 kb withheld. Both models are trained with the classification, supervised contrastive and Deep SAD objectives of Section 2.3, with the base identity channels visible. We build the read features exactly as in training, so the raw signal of each read is normalized by its median absolute deviation and the expected k-mer levels are z-scored. The windows, reads, sites, strands and coverage thresholds are the same as in the other arms, so that every comparison is paired read for read.

*Sites scored by each method.* We choose sites from the reference and read coverage only, never from calls or labels. Each method is queried at the base that carries the mark, as its model reads it. On the cell lines, the mark is 5mC at CpG, so both methods use every covered CpG in the window. On SPO1, the mark is 5hmU. RawMod therefore uses up to 200 T sites per window, drawn at random with a fixed seed. Using every T instead gives the same result (barcode 02: 0.79 against 0.78; barcode 05: 0.78 against 0.78). Dorado has no T model, so it uses its all-context cytosine model at 200 cytosines per window, drawn the same way with the same seed. The cap of 200 gives both methods the same number of sites. The other site classes in Table S9 (CpG and adenine) are controls, not main arms. MethylBERT sees only CpG calls.

*Combining Dorado and RawMod.* We test whether the two feature blocks are complementary. On the cell lines, we join them at the sites both score, the CpG sites. The per-site split standardizes every column, so a block's weight is set by its column count. We therefore first tile Dorado's two probabilities to the 96 columns of the RawMod embedding. On SPO1, our site rule gives the two methods no shared site, so a feature-level combination is not defined there. Instead, each method splits the reads at its own sites, Dorado at 200 random cytosines per window and RawMod at the T sites (up to 200), and both enter one consensus.

*Supervised baselines.* Neither supervised classifier runs as published on these data. Both are given advantages, which we state here because the comparison favors them.

MethylBERT expects short reads lying wholly inside a differentially methylated region. Its read extraction therefore returns nothing for 26 kb reads against 10 kb windows. We fragment reads to 500 bp, supply the window list and the direction of each window's methylation difference, and fine-tune out of fold. Its region encoder is an embedding indexed by region identity, so a window absent from training has no learned representation. We therefore use read-level folds, which is its own protocol. On SPO1, the five amplicon-versus-native contrasts are pooled into one data set of 21 windows with disjoint read sets. The native arm takes the role of the tumor class, and the direction of each window is taken from the bulk CpG methylation difference between the arms. MethylBERT's read parser assumes CpG-only calls. It is therefore given only the calls at CpG positions of the all-context basecall, which is the information a CpG-only model would supply.

ROCIT derives read labels from somatic variants, copy-number segments, phased haplotags and subclone assignments, which label roughly 1.4% of reads. Neither cell line here supplies that stack. COLO829 would need one assembled, and on HS578 the two lines differ at many germline sites, so somatic calls on that pair are not reliable. We therefore run its preprocessing unmodified on the HG008 sample, where the full labeling stack is published. We then apply the resulting model to our reads with no labels from our data. This is the only configuration in which it can be run here. The package itself is unmodified. The published example data for HG008 covers only three chromosomes. After the required validation and test chromosomes are held out, a single chromosome remains for training, giving 4,039 labeled reads, of which 384 are tumor.

##### S6.5. Further read-separation analyses

This subsection first defines how every method is scored. It then reports the analyses behind Table 2, first on the cell lines and then on SPO1.

*Scoring.* We score every method by per-window AUROC rather than ARI. ARI scores a partition, so it charges a method for where it places a threshold. That is the right question for a label-free clusterer, but the wrong one for a classifier that emits a calibrated probability. A label-free partition also has an arbitrary sign. Each window is therefore oriented by the truth, AUROC $:= \max(\text{AUC}, 1 - \text{AUC})$, which is one bit per window supplied by an oracle. We apply the same rule to the supervised methods, which do not need it, because a shared orientation rule gives a shared null. Under the null, this statistic is not 0.5 but $0.5 + \sqrt{(n+1)/3n^2}\sqrt{2/\pi}$, which exceeds 0.5 when a window holds few reads. We compute it identically for every dataset from the number of reads per window: SPO1 0.52, COLO829 0.57, HS578 0.59 and flow-cell control 0.62 (Table 2). Table 2 and Fig. 5e–h report the result. Table S9 gives the scores by site class, including the comparison at identical sites, and the window counts.

*Supervised methods on the cell lines.* The supervised classifiers also tell the two cell-line pairs apart in a way that the AUROC column does not show. A classifier trained on true labels emits a tumor probability directly, so it should not need the orientation step. On HS578, ROCIT is oriented correctly in 85% of the windows ($p = 2{\times}10^{-13}$) and reaches a median AUROC of 0.77, which does not differ from RawMod ($p = 0.34$), even though it was trained on another platform and cancer type from 384 labeled tumor reads. On COLO829, its orientation is inverted in 65% of the windows, so its score there comes from the orientation rule rather than from the classifier. Supervision therefore transfers between samples that differ by a consistent tumor-versus-normal methylation direction, and not between two cell types. This supports reading COLO829 as a cell-type contrast. MethylBERT stays near the null on both pairs, because a 500 bp fragment holds few of the differential CpGs of a window.

*Combining the feature blocks.* We test 11 ways of combining the RawMod embeddings with the Dorado calls, by tiling, by weighting, and by letting each method split the reads at its own sites. On the cell lines, none of them is better than Dorado alone. The only combination that scores higher, RawMod down-weighted to 24 columns on HS578 (0.876, $p = 0.042$), does not survive a Bonferroni correction for the 11 tests. On SPO1, the combination (0.839) is ahead of Dorado alone (0.688) in 19 of the 21 windows ($p = 5{\times}10^{-4}$) and ties with RawMod alone (0.796, paired $p = 0.9$).

*Flow-cell control.* Both HS578 lines were sequenced on two flow cells each. Within one line, reads from the two flow cells carry no biological difference. We therefore run every method unchanged on the reads of one line labeled by flow cell, separately for the tumor and the normal line. We use the same windows and the same minimum of 10 reads. The label-free methods run the same consensus. The supervised methods keep their per-read tumor probabilities and are scored against flow cell. Windows hold a median of 15 reads, which puts the chance level at 0.62. Every method lies at chance: RawMod 0.625, Dorado 0.600, the combination 0.600, MethylBERT 0.615 and ROCIT 0.619 (Wilcoxon against chance, all $p \geq 0.55$). The separation on the cell lines therefore does not come from flow-cell differences.

*Library preparation on SPO1.* The two SPO1 arms differ in library preparation as well as in modification, and no preparation-matched control exists for the native arm, because every thymine of the native genome carries 5hmU. We therefore estimate how much of the separation the library alone can explain. At the benchmark sites, two native libraries separate at an AUROC of 0.49–0.53 and two PCR libraries at 0.56–0.57, as Section S9.1 describes. Library identity therefore accounts for little of the 0.80 separation between the native and PCR arms. For this reason, Fig. 5e–h draws the chance level for SPO1 as for the cell lines, and we report the library controls here rather than drawing them.

*Models that did not train on these reads.* The region-holdout model of the main text was trained on the SPO1 reads outside 52.6–72.5 kb. Table S10 compares it with the released model, which saw every window in training, and with two models that were

**Table S9: Read separation by every method and site class on identical reads. Oriented AUROC: per-window median of** max(AUC, 1 − AUC); **−null: margin over the expected value for random scores. Correct sign: fraction of windows in which a supervised method's tumor probability is already oriented. Bold: best per dataset among the main arms. Indented rows score the same method at other site classes. On SPO1, the main arms are RawMod at T sites and Dorado's cytosine model at cytosines, 200 per window drawn at random. On the human lines, methods are also compared at identical sites; on SPO1, the site class scored is given per row. The first row of each method is the arm shown in Fig. 5e–h and Table 2. The null of the oriented AUROC depends on the number of reads in the window. A method trained on true labels should not need its sign corrected; correct sign is undefined for the label-free methods, whose cluster identity is arbitrary by construction.**

| Method | Labels used | Oriented AUROC med. | Oriented AUROC −null | Correct sign |
|---|---|---|---|---|
| *COLO829 5mC/CpG, two cell types (54 windows, null 0.57)* | | | | |
| RawMod | none | 0.773 | +0.201 | – |
| Dorado 5mC/5hmC | none | **0.999** | +0.424 | – |
| Dorado + RawMod | none | 0.998 | +0.424 | – |
| MethylBERT | these reads | 0.570 | +0.003 | 52% |
| ROCIT | another sample | 0.920 | +0.352 | 35% |
| *HS578 5mC/CpG, tumor/normal breast (100 windows, null 0.59)* | | | | |
| RawMod | none | 0.687 | +0.094 | – |
| Dorado 5mC/5hmC | none | 0.862 | +0.271 | – |
| Dorado + RawMod | none | **0.889** | +0.290 | – |
| MethylBERT | these reads | 0.638 | +0.047 | 73% |
| ROCIT | another sample | 0.767 | +0.160 | 85% |
| *SPO1 5hmU, PCR vs native (21 windows, null 0.52)* | | | | |
| RawMod, T sites | none | 0.796 | +0.272 | – |
| T + CpG sites | none | 0.837 | +0.284 | – |
| CpG sites only | none | 0.800 | +0.278 | – |
| A sites only | none | 0.796 | +0.268 | – |
| Dorado, C sites | none | 0.688 | +0.168 | – |
| T sites only | none | 0.592 | +0.076 | – |
| CpG sites only | none | 0.631 | +0.115 | – |
| Dorado C + RawMod T | none | **0.839** | +0.305 | – |
| MethylBERT | these reads | 0.764 | +0.248 | 86% |
| ROCIT | *n/a* (needs somatic calls, atlas) | | | |
| *Control: one HS578 line, two flow cells (null 0.62)* | | | | |
| RawMod | none | 0.625 | +0.000 | – |
| Dorado 5mC/5hmC | none | 0.600 | −0.023 | – |
| Dorado + RawMod | none | 0.600 | −0.023 | – |
| MethylBERT | these reads | 0.615 | −0.008 | – |
| ROCIT | another sample | 0.619 | −0.004 | – |

trained without SPO1 or without 5hmU. We make two observations. First, the region-holdout model matches the released model (0.796 against 0.811, paired $p = 0.09$) and is ahead of Dorado in all three windows inside its held-out region. Second, the models trained without 5hmU still separate the two arms (0.727 and 0.735, against a null of 0.52), but they are not ahead of Dorado. Part of the separation therefore needs no 5hmU in training, which agrees with the basecall identity result below, whereas the lead over Dorado comes from 5hmU in training.

**Table S10: SPO1 native-versus-PCR read separation with models that did not train on these reads. Median oriented AUROC over the 21 windows of Section S6, RawMod at 200 random T sites per window, identical reads and protocol for every row. Ahead: windows in which the model is ahead of Dorado's cytosine model; $p$: paired Wilcoxon signed-rank test against it. In-region windows: the three windows inside 52.6–72.5 kb, which the region-holdout model never saw. Null: 0.52.**

| Model | Trained on SPO1 | Median AUROC | Ahead of Dorado | $p$ | In-region windows |
|---|---|---|---|---|---|
| All data (released) | all reads | 0.811 | 17/21 | 0.002 | 0.78, 0.84, 0.81 |
| Region holdout (main text) | outside 52.6–72.5 kb | 0.796 | 17/21 | 0.002 | 0.79, 0.90, 0.83 |
| Without 5hmU (LOCO) | no | 0.727 | 14/21 | 0.27 | 0.70, 0.77, 0.71 |
| Without SPO1 (leave-out) | no | 0.735 | 12/21 | 0.79 | 0.64, 0.67, 0.73 |
| Dorado, cytosine model | – | 0.688 | – | – | 0.69, 0.75, 0.81 |

*Dorado without a thymine model.* Dorado separates these reads without calling the modification itself. On these reads, it emits 5mC/5hmC tags at cytosines only, and it makes no call at thymines. A thymine receives a probability only when the basecaller miscalls it as a cytosine, which then inherits a 5hmC score. This happens at 0.45–0.56% of the native and 0.18–0.26% of the amplicon read and T site pairs. We score a thymine without a call as unmodified (probability 0). At the same T sites, RawMod reaches an AUROC of 0.80 and Dorado 0.59, which is barely above chance, and RawMod is ahead in 19 of the 21 windows ($p = 3{\times}10^{-5}$).

RawMod also separates the arms at adenines. Scored at up to 200 adenines per window, it reaches 0.80, the same as at T sites, although the T sites are ahead in 15 of the 21 windows ($p = 0.03$). This adenine score is consistent with a response to the 5hmU at the neighboring bases. We therefore apply one rule to both methods and score each at the base its model reads: RawMod at thymines, where the 5hmU sits, and Dorado at cytosines, the base of its hydroxymethylation model.

*Identical CpG sites.* When both methods are scored at the same positions, the order between them depends on the base. At the CpG sites of the same windows (3–32 per window; one window has too few covered CpGs and is dropped), RawMod reaches 0.80 against 0.63 for Dorado and is ahead in 18 of the 20 windows ($p = 3{\times}10^{-5}$). As described above, RawMod is also ahead at thymines. Combining the two feature blocks at the CpG sites does not improve on RawMod alone.

*Basecalling 5hmU DNA.* 5hmU also lowers the accuracy of the basecall from which every method, including RawMod, starts. The super-accuracy basecaller of Dorado reads native SPO1 DNA with a median identity of 95.2%, against 98.9% for PCR amplicons of the same genome. Native reads carry about twice as many insertions and deletions, and their error rate is 2 to 5 times higher at every offset from the nearest T, without a peak at the T itself, as Fig. S7b shows. Because every T of this genome is 5hmU, these data cannot separate a local effect from a global one. Read identity alone separates native from PCR reads with a median per-window AUROC of 0.827 over 16 windows, against 0.589 for the native-versus-native and PCR-versus-PCR controls and 0.796 for RawMod at T sites. On this dataset, the difference between the arms is therefore a shift across the whole read, which a read-level identity score also detects. RawMod places the signal of each read on the reference sequence rather than on its basecall, as Section 2 describes.

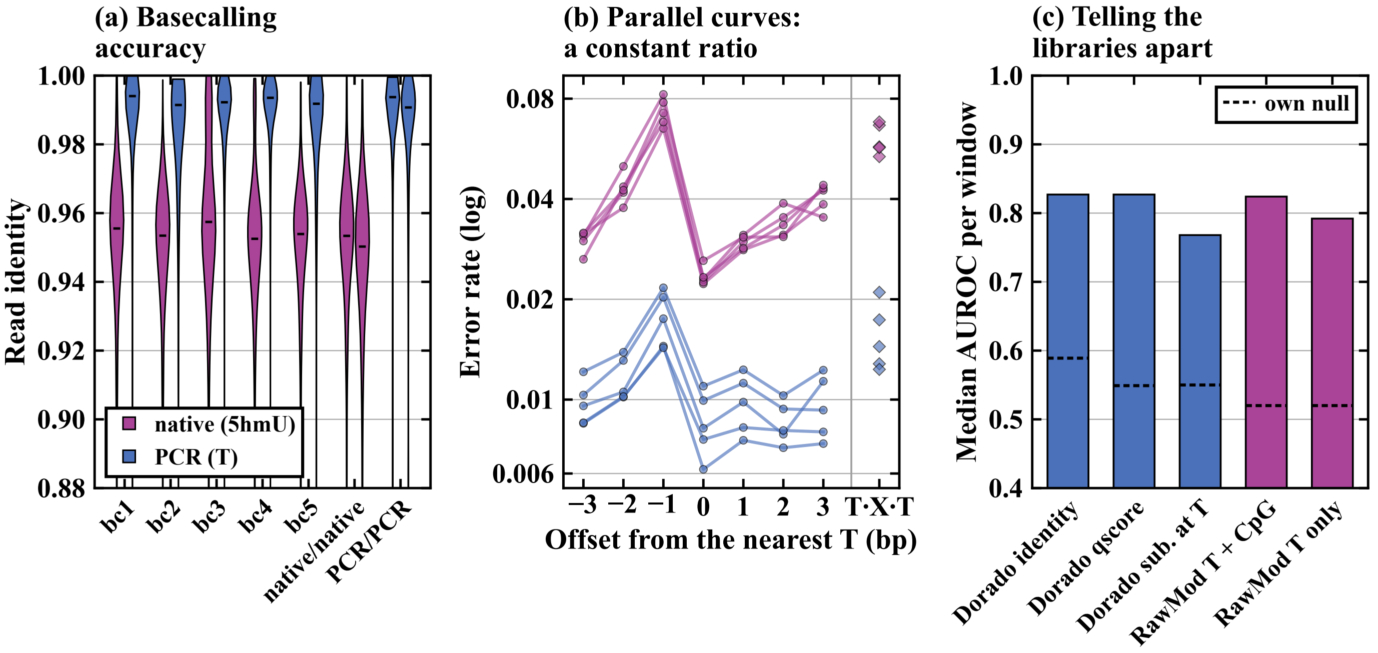


**Figure S7: Dorado basecalling of native (5hmU) and PCR SPO1 reads. (a) Read identity per library and in the two control contrasts. (b) Error rate against offset from the nearest T on the read's strand, one line per library, on loci that both libraries cover. A negative offset means the T lies before (5′ of) the base. T·X·T: bases with a T on both sides. (c) Median per-window AUROC for separating native from PCR reads; dashed: each score's own null.**

#### S7. Extended discussion

##### S7.1. Design for a genome scan

RawMod is small by design, because its intended use is a scan of every position of a genome. Its input is compact: the basecall and the refined segmentation have already placed the signal of each read on the reference, so every read contributes ten resampled current values per base over 21 bases. What remains for the model is to compare the reads at a site with the expected current, which requires only 0.23 million parameters. The cost of this design is its output. RawMod gives one score per site and strand, whereas UniMeth and Rockfish give a call for every read. The only per-read output of RawMod is the read embedding that Section 3.6 uses. One score per site fits a genome scan, in which each position is scored once from the reads that cover it. For example, RawMod scores the 2.17 million forward-strand thymines of the *Amphidinium carterae* contig in 4.8 min on one GPU, as Section S10 reports. Our code also supports running it without a GPU.

The design also lowers the computational cost. A map of each position of a bacterial genome takes RawMod less wall time than the four modification models of Dorado (8.6 against 9.9 min, basecalling included) and about a tenth of their GPU time. In exchange, our featurization uses more memory, as Section S10.4 shows. At the measured 7,500 sites per second, every position of both strands of a human genome would take about 230 GPU-hours on one RTX A5000. The human analyses here score CpG sites in selected windows.

##### S7.2. Why a label-free consensus works

The read-origin results show why a label-free consensus succeeds where single sites fail. At any one site, the split of the reads by the first principal component of their embeddings is noisy, because the signal of a single site is too weak to resolve the origin of a read. Combined over the sites of a window, the splits agree and the partition follows the sample, whereas two flow cells of one HS578 line stay at chance for any number of sites, as Section S6.5 shows. The gain therefore comes from methylation differences that accumulate along a read. This motivates read-origin methods that learn to pool evidence across the sites of a read and across reads, controlled for batch, rather than average a fixed per-site score.

##### S7.3. Limitations

RawMod has six limitations that future work can address. First, RawMod scores are not calibrated modification fractions. The model is trained to classify sites, so absolute estimates require orthogonal quantification. On the held-out regions, the predicted probability overstates the modified fraction in every bin, with an expected calibration error of 0.058, as Fig. S4 shows. Second, RawMod detects modifications but does not identify them, because its site embeddings do not separate modification types, as Section S3 shows. Naming a modification therefore requires an orthogonal assay or a modification-specific method. Third, RawMod trains on several public datasets that annotate modifications by motif presence rather than by measured methylation. Fourth, our read-origin analysis is preliminary, because it covers one chromosome of each of two cell-line pairs with complete tumor purity. Fifth, pairwise labels from matched modified and unmodified sites reduce, but do not remove, shortcuts through the organism and the library, and the site embedding still carries the dataset of origin. Sixth, outside the cytosine modifications, the accuracy of RawMod on the benchmark depends on having seen the library of the modification in training, as Section S9.8 shows. Training on more libraries per modification should reduce this dependence.

#### S8. Datasets and implementation

**Table S11: Datasets and their applications. Stages indicate training time usage.**

| Dataset | Modifications | Unmodified reference or ground truth | Use | Pileups |
|---|---|---|---|---|
| ONT synthetic 5-mer constructs [7] | 5mC, 5hmC, 6mA | Canonical construct (pairwise) | Core pool, stages 1–2, benchmark | 88,813 |
| SPO1 native genome | 5hmU | PCR amplicons (pairwise) | Core pool, stages 1–2, benchmark, application | 259,452 |
| *H. pylori* 26695 native [8] | 6mA, 4mC | Whole-genome amplification (pairwise) | Core pool, stages 1–2, benchmark | 89,312 |
| *E. coli* WT, DM, DM + M.SssI [8] | 6mA, 5mC | Motif annotation (training); Δ*dam*Δ*dcm* knockout at the same positions (benchmark) | Stage 2, benchmark | 179,920 |
| *Anabaena*, *T. denticola*, *H. pylori* J99 [8] | 6mA, 4mC | Motif annotation | Stage 2, benchmark (*Anabaena* only) | 56,381 |
| *A. thaliana*, HG001, HG002 [8, 9] | 5mC | Bisulfite or EM-seq | Stage 2, benchmark (*A. thaliana*, HG002) | 12,580 |
| Mouse brain, *O. sativa* [8] | 5mC | Bisulfite or EM-seq | Benchmark only | – |
| *Amphidinium carterae* contig_3762 | 5hmU | None | Application | 2.17 M sites |
| COLO829 / COLO829BL | 5mC | None | Application | — |
| HS578T / HS578Bst | 5mC | None | Application | — |

##### S8.1. Pairwise contrasts

Three datasets sequence the same positions in a modified and an unmodified state, so that only the modification separates the two libraries. The ONT synthetic constructs are 32 random sequences that cover every 5-mer, each sequenced fully substituted with 5mC, 5hmC or 6mA and again unmodified [7]. The *H. pylori* 26695 libraries compare native DNA, which carries 6mA and 4mC at methyltransferase motifs, with a whole-genome amplification of the same DNA [8]. The SPO1 libraries compare native DNA, in which every thymine is 5hmU, with PCR amplicons of the same 132 kb genome; we sequenced both for this study (Section S8.3).

All datasets were sequenced with the R10.4.1 chemistry at 5 kHz. Training uses at most $N$=15 reads per pileup and up to five pileups per site.

#### S8.2. Held-out genomic regions

For each organism, a contiguous region of about 15% of the genome is removed from training, together with every read that overlaps it. Section S9.7 lists the region of every organism.

#### S8.3. SPO1 sequencing data

We sequenced SPO1 to provide a public nanopore dataset of 5hmU-containing DNA together with an unmodified copy of the same genome. SPO1 is a *Bacillus subtilis* phage with 5hmU in place of every thymine [10]. No released modification caller includes a 5hmU model.

**S8.3.1. Sample preparation.** SPO1 (ATCC 27370-B1) was obtained from the American Type Culture Collection, and genomic DNA was extracted with the Norgen Biotek DNA Isolation Kit. Five primer sets were designed from the reference genome to amplify select regions with sizes from 6.4–19.3 kb, as shown in Table S12. PCR amplification was performed on these regions with Takara One Shot LA PCR Mix Ver.-2.0, and the products were verified on a Lonza FlashGel system. Amplicons were purified with AMPure XP beads at a 0.8× bead-to-sample ratio, and amplicons and native DNA were quantified with a Qubit fluorometer. Because PCR amplifies unmodified nucleotides, the amplicons carry thymine in place of 5hmU, so they serve as unmodified controls.

**S8.3.2. Sequencing.** Libraries were prepared with the Oxford Nanopore Native Barcoding Kit 24 V14 (SQK-NBD114.24). Barcodes 01–05 were assigned to the five amplicons, and barcodes 06 and 07 to native DNA. The pooled libraries were sequenced on a single R10.4.1 flow cell (FLO-MIN114, FBA96731) on a MinION Mk1B platform running MinKNOW (v24.11.10). Operations used a sampling rate of 5 kHz and a 400 bps translocation speed. Two runs on this flow cell produced 5,761,466 reads.

**S8.3.3. Demultiplexing and mapping.** All reads were basecalled with Dorado 1.4.0 `sup@v5.2.0` and demultiplexed with `dorado demux`, which requires a barcode at either end of a read. Of the 5,761,466 reads, 4,359,197 were assigned to one of the seven barcodes, 1,402,253 remained unclassified, and 16 were absent from the demultiplexed output (Table S12). Reads were aligned to the SPO1 reference genome (GenBank FJ230960.1, RefSeq NC_011421.1, 132,562 bp) [10] with minimap2 2.26. Alignments were sorted and indexed with SAMtools. Each amplicon library is enriched 5.5× to 179× its target region, and both native libraries cover the entire genome. The benchmark retains only reads whose library of origin agrees with the alignment (Section S9.1), which removes reads whose barcode disagrees with their alignment. All analyses in this study use the reads of the first run.

**Table S12: SPO1 sequencing libraries. Regions are given in coordinates of reference GenBank FJ230960.1.**

| Barcode | Library | Region | Size (bp) | # Reads |
|---|---|---|---|---|
| 01 | PCR amplicon 1 | 40,727–47,100 | 6,374 | 483,945 |
| 02 | PCR amplicon 2 | 59,437–77,340 | 17,904 | 865,851 |
| 03 | PCR amplicon 3 | 1,381–20,644 | 19,264 | 493,395 |
| 04 | PCR amplicon 4 | 123,573–130,680 | 7,108 | 383,470 |
| 05 | PCR amplicon 5 | 87,767–105,951 | 18,185 | 837,050 |
| 06 | Native DNA | whole genome | 132,562 | 872,386 |
| 07 | Native DNA | whole genome | 132,562 | 423,100 |
| – | Unclassified | – | – | 1,402,253 |

#### S8.4. *Amphidinium carterae* sequencing data

*Amphidinium carterae* strain NCMA 1314 was obtained from the National Center for Marine Algae and Microbiota. Its culturing, DNA isolation and sequencing are described in [11, 12]. We summarize them here.

**S8.4.1. Sample preparation.** After DNA isolation, fragments were size-selected with the SRE or SRE-XL kit (Pacific Biosciences). Libraries were prepared with the Ligation Sequencing Kit (SQK-LS114, Oxford Nanopore).

**S8.4.2. Sequencing.** Libraries were sequenced on R10.4.1 MinION and PromethION flow cells. Reads were basecalled in duplex mode with Dorado and the `dna_r10.4.1_e8.2_400bps_sup@v4.3.0` model. The assembly was built in two stages. First, all reads were assembled with Flye 2.9.1 [13]. Bacterial and mitochondrial contigs were identified by BLASTN [14] hits, coverage and AT content, and removed. Nuclear contigs had about 60× coverage, a slight GC bias and often high RNA-seq mapping. Mitochondrial contigs were AT-rich and carried fragments of the *coxI*, *coxIII* and *cob* genes. Bacterial genomes assembled as single contigs without RNA-seq mapping [11]. Second, reads that mapped to non-nuclear contigs with minimap2 [15] were removed, and the remaining reads were assembled again with Flye. The transcriptome [16] was placed on the contigs with BLASTN, and RNA-seq reads were mapped with HISAT2 [17].

**S8.4.3. Preprocessing.** Two PromethION flow cells (11,493 and 17,377 reads) were re-basecalled from their raw signal with Dorado 0.9.2 `sup@v5.0.0`. Output move tables were refined against the R10.4.1 9-mer level table, and every reference T on the forward strand with ≥10 reads was featurized and queried with RawMod. The reference is contig JBNFNT010000839.1 (GenBank), the largest contig of the assembly described in [12]. Exonerate [18] was used in est2genome mode with the transcriptome to create initial gene models, which were corrected manually using Apollo [19] with RNA-seq reads mapped by HISAT2 (SRA SRR5220732–SRR5220737, SRR1294391–SRR1294394, SRR1296757–SRR1296758, SRR8513682–SRR8513683 and SRR1610335). The contig comes from a draft long-read assembly of a 1.25 Gb repeat-rich genome. Interspersed repeats cover 54% of it, and only 237 of its genes are annotated. 69% of the contig aligns to the strain used for previous 5hmU mapping, at 1.4% divergence. Our reads align to the contig with a median per-read error of 5.4%. Part of this error likely comes from 5hmU itself, which lowers basecalling accuracy on SPO1 DNA (Section S6.5), and part from errors in the draft assembly. Assembly and annotation errors can therefore shift the scores of individual regions.

**Table S13: Tool versions.**

| Tool | Version | Use |
|---|---|---|
| RawMod | 1.0 | main method |
| Dorado | 0.9.2, `dna_r10.4.1_e8.2_400bps_sup@v5.0.0` | basecalling, refinement |
| minimap2 | bundled with Dorado | alignment |
| Remora | 3.3.0 | move-table refinement against the level table |
| samtools | 1.22 | BAM processing |
| PyTorch | 2.12.0 (CUDA 13.0) | model training |
| level table | R10.4.1 9-mer means (Uncalled4) | expected current, move refinement |
| Dorado (benchmark) | 1.4.0, `sup@v5.0.0`, `6mA@v3`, `4mC_5mC@v3`, `5mC_5hmC@v3`, `5mCG_5hmCG@v3` | benchmark (Section 3.2) |
| Dorado (SPO1 demultiplexing) | 1.4.0, `sup@v5.2.0` | barcode assignment |
| UniMeth | 0.3.3 (released models); 0.3.1, commit 11215d4 (fine-tuned) | benchmark |
| Rockfish | 0.0.2, `rf_5kHz.ckpt` | benchmark |
| DeepMod2 | commit ff910f6, `bilstm_r10.4.1_5khz_v5.0` | benchmark |
| MethyNano | commit 4f66a60 | benchmark |

#### S8.5. Data availability

The RawMod code, trained models, and scripts that reproduce every table and figure are available at `https://github.com/STORMgroup/RawMod`. The benchmark scripts download each public dataset from its original source. The ONT synthetic oligonucleotides are from the ONT modification validation data [7]. The bacterial, plant, mouse and human datasets are from Kulkarni et al. [8] and ONT open data [9]. The COLO829 and COLO829BL cell lines are described by Valle-Inclán et al., and their reads are from ONT open data [20, 21]. The HS578T and HS578Bst reads are from the CASTLE panel [5, 6] (NCBI SRA BioProject PRJNA1086849). The SPO1 reads that we sequenced for this study under Section S8.3 are deposited under BioProject PRJNA1549975. The SPO1 reads are also in SquiDBase [22] under accession SQB000031. The *Amphidinium carterae* reference is contig JBNFNT010000839.1 in GenBank [12].

The *Amphidinium carterae* reads and assembled contigs are available under NCBI BioProject PRJNA1110228. The raw signal (POD5) of the two flow cells used here is in SquiDBase [22] under accession SQB000028.

#### S8.6. Where a modification changes the signal

Different modification types perturb the output signal in unique ways (Fig. 1b). Across pairwise sites of a modified library and its unmodified control, 5mC and 5hmC change the read-median current most, 6mA one base downstream, and 4mC split between one base upstream and again two bases downstream. In SPO1, where every thymine carries 5hmU, neither the read-median current nor the dwell time of the native library differs from that of its PCR copy at a single position. The modified base therefore does not always sit where its signal is strongest.

#### S8.7. Model and training details

This section gives the sizes and settings that Section 2 leaves out. The model has three parts: a convolutional encoder that embeds each read, a cross-read transformer that compares the reads of a site, and a head that scores the site.

**S8.7.1. RawMod methods implementation.** A shared convolutional encoder embeds each read (Fig. 2). The encoder applies a three-layer convolutional stem inspired by ConvFormer [23], a 32–32–48 channel projection whose last layer has stride 2. Then, an Inception-style [24, 25] multi-branch block with kernel sizes 1, 3, and 5 and pooling across 64 channels feeds a grid-reduction block of stride 2 and 184 channels. The last part of the encoder consists of a $d$=96 projection and mean pool, resulting in a $\mathbf{e}_r \in \mathbb{R}^{96}$ embedding per read.

RawMod uses $N$=15 reads per pileup ($H$=16 rows), a window of $W$=21 bases and $L$=10 samples per base, so each pileup has $C \times H \times W \cdot L$ = 11×16×210 values. Read embeddings have $d$=96 dimensions. The cross-read transformer has two pre-norm layers with four heads, a feed-forward width of 192, GELU activations, and a dropout of 0.4. The SupCon projection head is a two-layer MLP with 128 outputs and temperature $\tau$=0.2, and the Deep SAD map $h$ is a linear layer without bias with 16 outputs, with margin $m$=2 and weight $\eta$=1. The first training stage runs for 15 epochs. The second runs for at most 50 epochs and stops after 15 epochs without improvement on the validation set.

We use AdamW (learning rate $1.2\times10^{-3}$, weight decay $10^{-3}$) with batch size 512, 500 warm-up steps, reduce-on-plateau scheduling (factor 0.5, patience 7), and gradient clipping at 1.0. Data augmentation randomly masks 0–30% of read rows for generalization to low coverage and adds Gaussian noise ($\sigma$=0.05) to the current channel for generalization to noisy data.

#### S9. Benchmark procedure

This section describes how we produce Table 1, Table S21, Table S22 and Fig. 3. The goal of the benchmark is to score every tool on the same reads, at the same sites, with the same evaluation protocol, so that a difference between two columns comes from the tools and not from the evaluation. The pipeline is a set of six steps: read subsetting, basecalling, read selection, one job per tool and sample, aggregation, and scoring. Section S9.10 describes the released scripts that run these steps.

##### S9.1. Read sets and basecalling

**Read subsetting.** To ensure a consistent comparison, we first select a fixed set of reads for each sequencing sample before running the benchmarked tools. The benchmark uses 16 sequencing samples, each drawn from a predefined subset of its source data. We select the reads using the following criteria:

- **ONT synthetic oligonucleotides (5mC, 5hmC, 6mA and unmodified control)**: We use the held-out 20% of reads from each library (31,453–31,989 reads).
- **SPO1 (native barcode 07 and PCR barcode 01), *H. pylori* 26695 (native and whole-genome-amplified), *Anabaena* and HG002**: We select up to 15,000 alignments per sample (11,908–14,905 reads). For the whole-genome-amplified *H. pylori* sample, selection is taken only over the region covered by the native sample.
- **Mouse brain**: We select the 39,622 reads that overlap the chromosome 1 EM-seq sites.
- ***O. sativa***: We select up to 15,000 reads whose alignment starts in the first 13 Mb of chromosome 1.
- ***E. coli* K-12 samples (wild type, Δ*dam*Δ*dcm*, and Δ*dam*Δ*dcm* treated with M.SssI)**: We select the reads whose primary alignment starts in the held-out region (1.74–2.44 Mb). Using the same genomic region across the three samples ensures that the pairwise rows compare the same region.
- ***A. thaliana***: We select up to 100,000 reads whose alignment starts on chromosome 4, the held-out chromosome.

In cases where the number of eligible reads exceeds the specified limit, we keep those with the lowest seeded hash ranks. This ensures that the benchmark does not depend on the order in which reads are processed or on the tool being evaluated.

**Basecalling and preprocessing.** We basecall every read set once with Dorado 1.4.0 and the `dna_r10.4.1_e8.2_400bps_sup@v5.0.0` model. We emit move tables (`-emit-moves`) and align reads to the sample's reference (`-reference`). We basecall ourselves rather than using any released BAM so that every tool sees identical reads, alignments, and signal scaling. All tools except MethyNano use the resulting BAM files. MethyNano's feature extraction step requires move tables from Dorado 0.9.2 with the `hac@v4.2.0` model. We therefore re-basecall the same POD5 reads with that model for MethyNano only. The read identifiers are unchanged, so MethyNano's calls are restricted to the same selected reads (Section S9.2). RawMod refines the move tables using Remora against the Uncalled4 R10.4.1 9-mer level table (Section 2.1).

##### S9.2. Read selection and site definition

**Read selection.** We only use primary alignments with a mean read quality of at least 10, keeping low-quality reads out of every site. To ensure that differences in sequencing coverage do not affect comparisons between tools, we rank eligible reads using a seeded BLAKE2b hash of their read identifiers (seed 0) and process them in increasing hash order. Each read contributes to an aligned reference position (including matches and mismatches) on its respective strand only if fewer than 10 reads have already been selected for that position. This procedure retains the 10 lowest-ranked eligible reads that cover each site, and the selection is the same for every tool. Sites with fewer than 10 eligible reads on a strand are excluded from evaluation. With this protocol, no tool can gain from deeper sites. Tools that skip reads due to additional filtering criteria, such as read length, mapping quality, or signal quality, are charged for the skipped reads.

*SPO1 read filtering.* The SPO1 libraries require additional filtering, because their labels are otherwise contaminated. The PCR libraries of SPO1 are single amplicons (barcode 01 spans 40,727–47,100, barcode 02 59,437–77,340). They were demultiplexed on one end only, so some reads were assigned to the wrong barcode. This contamination is particularly important in the held-out region, which is not covered by the barcode 01 amplicon. Inside this region, 72% of the reads assigned to barcode 01 carry over from native DNA, and 11% of reads assigned to native barcode 07 come from the PCR product of barcode 02. We therefore keep only the reads whose origin the alignment itself supports:

- **Unmodified (PCR) reads.** We retain the reads of barcode 01 whose primary alignment starts or ends within 30 bp of a barcode-02 primer or product end. These reads must also have a mean quality score of at least 18 and an alignment identity of at least 0.98. This filtering retains 1,870 of 7,561 reads.
- **Modified (native) reads.** We retain the reads of barcode 07 that are not anchored at any primer or product end of barcodes 01–05. This filtering retains 12,334 of 14,930 reads.

For the SPO1 benchmark, we evaluate thymine positions within the interval 59,437–72,503 that have at least 10 selected reads in both the filtered native and PCR datasets. Every tool is evaluated using these selected reads and positions. To check if the comparison measures the modification rather than library identity, we compare libraries with the same expected modification status at the same sites. Two native libraries (barcodes 06 and 07) separate at an AUROC of 0.49–0.53, and two PCR libraries (barcodes 01 and 02) separate at 0.56–0.57. These controls show little library effect within an arm but cannot exclude a difference between native and PCR preparation.

##### S9.3. Tools and models

This subsection lists how we run each tool, so that every tool is used as its documentation intends. Table S14 lists every tool, version, model and non-default setting. Before scoring, we identify which modifications and sequence contexts each tool is expected to call. This record decides which cells are bold in Table 1 and which models enter the matched view (Table S22).

*UniMeth.* We run the released UniMeth models with UniMeth 0.3.3 at `-frequency 5khz`. We use this version because version 0.3.1 skipped POD5 calibration on its 5 kHz path, which version 0.3.2 fixed.

*UniMeth-FT*$^\dagger$. UniMeth releases no model for three of our modifications. UniMeth-FT$^\dagger$ fills this gap by giving UniMeth the same training labels as RawMod for these three modifications. We fine-tune one model each for 5hmC (oligonucleotide 5hmC library against the control) and 4mC (*H. pylori* 26695 native against whole-genome amplified), starting from the released UniMeth R10.4.1 5 kHz 5mC model. We fine-tune one model for 5hmU (SPO1 native against PCR), starting from the released R10.4.1 5 kHz 6mA model. Each model trains for 3,000 steps with batch size 32 on the benchmark read sets of these libraries. A base is labeled only if its position is a training position of the benchmarked RawMod checkpoint: modified at its positives and unmodified at its negatives. Every other base is masked from the loss, so the held-out regions of the benchmark stay unseen. UniMeth-FT 5hmU reaches 0.965 on SPO1, the highest 5hmU AUROC of any model, because it is fine-tuned on the same SPO1 training positions as RawMod as shown in Table 1.

These models need a 5hmU token, so we use the patched UniMeth 0.3.1 for training and inference. In that version, `-frequency` selects only the signal normalization. For basecalls from Dorado 0.9.2 or later, its `4khz` setting applies exactly the normalization that the released 0.3.3 applies at `5khz`, and its own `5khz` setting assumes older basecalls. We therefore run it with `-frequency 4khz` on our 5 kHz data.

*MethyNano.* MethyNano's default minimum read length of 500 bases discards every oligonucleotide read (about 112 bases). We therefore run it with `-min-read-bases 50` on all samples. MethyNano's released calling script does not report a reference coordinate for each call. We therefore call it with `predict_aln.py`, a script adapted from the released one that maps each called read position to the reference through the read alignment. We release this script with our benchmark code as a patch to MethyNano commit 4f66a60.

*Why MethyNano is near chance.* We test whether MethyNano's low scores come from its model or from its released pipeline. The released checkpoints are sound: on the authors' pre-extracted test features, they reach an AUROC of 0.915–0.921, matching the reported 0.92. The features that the released calling script builds from Dorado move tables differ from these training features in segment length and variance, and the script gives near-flat calls even on the authors' own example reads. MethyNano's benchmark results therefore measure its released pipeline, not the accuracy its model could reach with features extracted as in training.

*Uncalled4.* Uncalled4 [26] detects a modification as a difference between the signal of a sample and that of an unmodified control, so we run it against the control library of each pairwise row (Table S18). We align every read with `uncalled4 align` (version 4.1.1) and the released R10.4.1 9-mer pore model, with a minimum alignment length of 50 bases, because the default of 100 discards about 45% of the oligonucleotide reads. The score of a site is the two-sample Kolmogorov–Smirnov statistic between the currents of the site's 10 selected reads and the currents of every other control read at the same position and strand. On rows without a control library, the statistic is taken against the pore model's distribution for the 9-mer. Uncalled4 is not trained on modified DNA and is not held out: as in normal use, its control includes the control library's reads in the held-out regions. It is scored at the same sites as every other column.

*NovaMod.* NovaMod [27] learns the signal of unmodified DNA with a variational autoencoder over 9-mer windows of per-base signal statistics and base qualities, and scores each base of a read by its reconstruction error. We run the released code (commit

**Table S14: Tools in the benchmark. "Trained for" is taken from each tool's documentation before scoring; unlisted settings are defaults. †Fine-tuned by us (Section S9.3). *Not held out: uses the unmodified control library's reads, including those in the held-out regions. "Trained for" decides the bold cells of Table 1 and the models used in Table S22. UniMeth-FT† is fine-tuned on the training positions of RawMod in the oligonucleotide, *H. pylori* 26695 and SPO1 libraries.**

| Tool | Version | Model(s) | Trained for | Settings |
|---|---|---|---|---|
| RawMod | this work | `mixed` checkpoint (final model) | 5mC, 5hmC, 6mA, 4mC, 5hmU; any context | one pileup per (site, strand) built from exactly the 10 selected reads; moves refined with Remora against the Uncalled4 R10.4.1 9-mer level table; reverse-strand pileups in read orientation |
| Dorado | 1.4.0 | `sup@v5.0.0` with `6mA@v3`, `4mC_5mC@v3`, `5mC_5hmC@v3`, `5mCG_5hmCG@v3` | 6mA; 4mC+5mC; 5mC+5hmC; 5mC+5hmC at CpG | one run per modification model with `–reference`; per-read probability = sum of the ML probabilities over every code the model emits |
| UniMeth | 0.3.3 | released R10.4.1 5 kHz 5mC and 6mA models | 5mC (CpG, CHG, CHH); 6mA | `infer –frequency 5khz –pore_type R10.4.1`, batch size 128; per-read probability = `prob_1` |
| UniMeth-FT† | 0.3.1 (commit 11215d4) with the collaborators' 5hmU patch | 5hmC and 4mC models fine-tuned by us from the released 5mC model, and a 5hmU model from the released 6mA model, on the training positions of RawMod | 5hmC; 4mC; 5hmU | `–frequency 4khz` (training and inference), 3,000 steps, batch size 32; 5hmU with `–hmU 1` |
| UniMeth-Agn | 0.3.1 (commit 11215d4) with the 5hmU patch and our modification-agnostic patch | one model fine-tuned by us from the released 5mC model on every training image of RawMod (Section S9.4) | any modification | `–frequency 4khz`, 12,000 steps, batch size 32, class-balanced loss; a question at every C, A and T |
| DeepMod2 | commit ff910f6 | `bilstm_r10.4.1_5khz_v5.0` | 5mC at CpG | `detect`, default thresholds, batch size 1024 |
| Rockfish | 0.0.2 | `rf_5kHz.ckpt` | 5mC at CpG | `inference`, batch size 4096 |
| MethyNano | commit 4f66a60 with our `predict_aln.py`, which maps each call to the reference through the read alignment | *H. sapiens*, *A. thaliana* and *O. sativa* checkpoints | 5mC | reads re-basecalled with Dorado 0.9.2 `hac@v4.2.0` with moves, as the tool requires; `–min-read-bases 50` (the default of 500 discards every synthetic-oligonucleotide read); site score = maximum over the three checkpoints |
| Uncalled4* | 4.1.1 | released `dna_r10.4.1_400bps_9mer` pore model | any modification, any context (against an unmodified control) | `align –min-aln-length 50` on every read; site score = two-sample Kolmogorov–Smirnov statistic of the 10 selected reads against the control library's other reads at the site, or against the pore model without a control (Section S9.3) |
| NovaMod* | commit fbe72bc | one model per unmodified control library, trained by us with the released recipe (`train.example.json`) | any modification, any context (against unmodified DNA) | `val.py` site mode on every read at the candidate sites; site score = mean reconstruction error of the 10 selected reads (Section S9.3) |

fbe72bc) with its recipe (9-mers, 20 epochs, last epoch used) on raw signal BAM files built from our Dorado basecalls. The documented checkpoints are not in the repository, so we train one model per unmodified control library, on every read of that library with mean quality ≥10 that is in no benchmark read set (37,000–319,000 reads per library). Each model is scored on the pairwise rows that use its library and is blank elsewhere. The training reads include the control library's reads in the held-out regions, so NovaMod is not held out. A site's score is the mean reconstruction error of its 10 selected reads.

*Library identity.* A pairwise row compares two libraries. A method can therefore score the row highly by recognizing the library rather than the modification. To test this, we score 1,500 positions inside the held-out *E. coli* region, on both strands, each from the benchmark's 10 selected reads. These positions have no GATC, CCWGG, CG or EcoKI site within 8 bases, so they are unmodified in the wild-type, M.SssI and Δ*dam*Δ*dcm* libraries alike. A method that sees only modifications should not separate the libraries at these positions. RawMod does not (AUROC 0.502 for wild type against Δ*dam*Δ*dcm* and 0.538 for M.SssI against Δ*dam*Δ*dcm*). Uncalled4 does not either for the wild type (0.521; 0.736 for M.SssI). NovaMod, trained on the Δ*dam*Δ*dcm* library, separates both libraries from it almost perfectly (0.998 and 0.998). Its near-perfect *E. coli* rows therefore measure the library it was trained on, not the modification.

We extend this test to every pairwise row and every modification caller (Table S15a). For each row, we take up to 5,000 background positions of the row's base inside the held-out region, scored from the 10 selected reads in both libraries. A background position is not a site of the row and lies more than 10 bases from every known modified site of the organism, so that no known modified base falls inside RawMod's 21-base window. The designed sites of the oligonucleotides are dense, so there we keep positions more than 3 bases from every designed site, which leaves 81–98 positions. We make three observations:

- On the *H. pylori* and oligonucleotide rows, RawMod and the released callers stay between 0.34 and 0.59, except UniMeth on the 5mC oligonucleotides (0.79).
- UniMeth-Agn, which is fine-tuned on reads of these libraries, separates the native library from its control on all five of these rows (0.75–0.88). UniMeth-FT† does the same on its 5hmC row (0.99) and, more weakly, on its 4mC row (0.69).
- On the M.SssI row, RawMod (0.336, scoring the M.SssI library lower) and UniMeth-Agn (0.776) separate the libraries at cytosines outside CpG. The positions of any base far from every motif above do not show this separation (0.538). The AUROC of this row is 1.000 for both tools, with or without the correction below.

We then remove what a tool gains from telling the libraries apart (*baseline-corrected AUROC*, Table S15b). We replace each site score by its percentile among the same tool's scores at the background positions of the same library, and we recompute the AUROC of the row. A shift of all scores of one library cancels, whereas the modification signal at the row's sites does not. One-library rows compare sites of one library, so the correction leaves them unchanged. The SPO1 row has no background, because every thymine of the native phage carries 5hmU. With each tool's matched models (Table S22), the correction moves RawMod's mean over the five modifications from 0.964 to 0.948 and Dorado's from 0.896 to 0.895. It moves UniMeth-Agn's mean from 0.960 to 0.917, mostly through the *H. pylori* 4mC row (0.994 to 0.787). RawMod's largest change is on the 6mA oligonucleotides (0.927 to 0.828, from 98 background positions). Library recognition therefore inflates UniMeth-Agn's pairwise rows more than RawMod's.

**Table S15: Library control of the pairwise rows inside the held-out regions. (a) AUROC between the native and the control library at background positions without a known modification; 0.5: the libraries are indistinguishable. (b) Baseline-corrected AUROC, with the matched AUROC below it in brackets. –: no matched model or <50 background positions. †Fine-tuned by us. Tool abbreviations as in Table 1; DM2: DeepMod2, MN: MethyNano. Background positions are positions of the row's base, scored from the 10 selected reads in both libraries, more than 10 bases from every known modified site (3 bases from every designed oligonucleotide site). The baseline-corrected AUROC replaces each site score by its percentile among the same tool's background scores in the same library; the matched AUROC of Table S22 is in brackets. The bottom rows give the mean and the lowest of the five per-modification means over all 15 rows, uncorrected and with the pairwise rows corrected (one-library rows and the SPO1 row unchanged). The CpG-only callers have fewer than 50 background positions on the oligonucleotides. The comparative methods are not shown, because they score a sample against its control by design. UniMeth-FT† is fine-tuned on RawMod's training positions outside the held-out regions.**

| Row | RawMod | Dor. | UM | UM-FT† | DM2 | RF | MN | UM-Agn |
|---|---|---|---|---|---|---|---|---|
| *(a) Native vs. control at unmodified background positions* | | | | | | | | |
| *E. coli* M.SssI vs. Δ*dam*Δ*dcm* (*n*=5,000) | 0.336 | 0.513 | 0.336 | – | 0.480 | 0.481 | 0.520 | 0.776 |
| *E. coli* Dcm vs. Δ*dam*Δ*dcm* (*n*=5,000) | 0.476 | 0.459 | 0.511 | – | – | – | 0.505 | 0.667 |
| ONT Benchmark – 5mC vs. control (*n*=81) | 0.460 | 0.365 | 0.794 | – | – | – | 0.332 | 0.747 |
| ONT Benchmark – 5hmC vs. control (*n*=87) | 0.476 | 0.383 | – | 0.990 | – | – | – | 0.800 |
| *E. coli* Dam vs. Δ*dam*Δ*dcm* (*n*=5,000) | 0.472 | 0.559 | 0.538 | – | – | – | – | 0.509 |
| *H. pylori* 26695 vs. WGA (GCATG) (*n*=5,000) | 0.512 | 0.500 | 0.457 | – | – | – | – | 0.844 |
| ONT Benchmark – 6mA vs. control (*n*=98) | 0.585 | 0.507 | 0.337 | – | – | – | – | 0.875 |
| *H. pylori* 26695 vs. WGA (TCTTC) (*n*=5,000) | 0.498 | 0.500 | – | 0.694 | – | – | – | 0.866 |
| *(b) Baseline-corrected AUROC (matched AUROC in brackets)* | | | | | | | | |
| *E. coli* M.SssI vs. Δ*dam*Δ*dcm* | 1.000 (1.000) | 1.000 (1.000) | 1.000 (1.000) | – | 0.895 (1.000) | 0.996 (1.000) | 0.510 (0.529) | 1.000 (1.000) |
| *E. coli* Dcm vs. Δ*dam*Δ*dcm* | 0.999 (0.999) | 1.000 (1.000) | 0.995 (0.990) | – | – | – | 0.492 (0.498) | 0.999 (1.000) |
| ONT Benchmark – 5mC vs. control | 0.925 (0.936) | 1.000 (1.000) | 0.781 (0.949) | – | – (0.837) | – (0.857) | 0.571 (0.337) | 1.000 (1.000) |
| ONT Benchmark – 5hmC vs. control | 0.932 (0.953) | 1.000 (1.000) | – | 0.950 (1.000) | – | – | – | 1.000 (1.000) |
| *E. coli* Dam vs. Δ*dam*Δ*dcm* | 0.981 (0.978) | 1.000 (1.000) | 1.000 (1.000) | – | – | – | – | 0.980 (0.953) |
| *H. pylori* 26695 vs. WGA (GCATG) | 0.980 (0.995) | 0.992 (0.995) | 0.991 (0.994) | – | – | – | – | 0.966 (1.000) |
| ONT Benchmark – 6mA vs. control | 0.828 (0.927) | 1.000 (1.000) | 0.994 (0.982) | – | – | – | – | 0.983 (1.000) |
| *H. pylori* 26695 vs. WGA (TCTTC) | 0.951 (0.981) | 0.982 (0.984) | – | 0.927 (0.988) | – | – | – | 0.787 (0.994) |
| Mean over the five modifications | 0.964 | 0.896 | 0.697 | 0.791 | 0.571 | 0.571 | 0.503 | 0.960 |
| baseline-corrected | 0.948 | 0.895 | 0.694 | 0.768 | – | – | 0.508 | 0.917 |
| Worst modification | 0.930 | 0.500 | 0.500 | 0.500 | 0.500 | 0.500 | 0.500 | 0.825 |
| baseline-corrected | 0.930 | 0.500 | 0.500 | 0.500 | – | – | 0.500 | 0.787 |

*Each library on its own.* The same background also tests each library separately (Table S16). This test asks whether a caller reads the modification or the sequence context of the site. In the native library, we compare the scores at the row's modified sites with the scores at the background positions of that library. This comparison measures detection with no difference between libraries to exploit. In the control library, we compare the scores at the same sites, which are unmodified there, with the background of the control library. A caller that reads the modification gives 0.5 at the control sites. A caller that scores the sequence context of the site, such as its motif, gives more or less than 0.5. The bottom of Table S16 gives the mean distance from 0.5 over the rows that each tool covers. A tool that detects nothing also gives 0.5, so we read this measure together with the AUROC on the modified datasets. For example, MethyNano is closest to 0.5 and also has the lowest AUROC (Table S21).

**Table S16: Within-library AUROC (native above, control below) of the pairwise rows inside the held-out regions. Scored against the background positions of Table S15. A control value of 0.5 means the tool does not score the site's sequence context as modified. Bottom: mean |control − 0.5| and number of rows. Tool abbreviations as in Table 1; DM2: DeepMod2, MN: MethyNano. Native: the modified sites of the row against the background of the native library. Control: the same sites, unmodified, against the background of the control library. Matched models only, as in Table 1; UniMeth-Agn as in Table S17.**

| Row | RawMod | Dor. | UM | UM-FT† | DM2 | RF | MN | UM-Agn |
|---|---|---|---|---|---|---|---|---|
| *E. coli* M.SssI vs. Δ*dam*Δ*dcm* | 1.000 | 1.000 | 1.000 | – | 1.000 | 1.000 | 0.538 | 1.000 |
| | 0.214 | 0.721 | 0.555 | | 0.992 | 0.985 | 0.528 | 0.587 |
| *E. coli* Dcm vs. Δ*dam*Δ*dcm* | 1.000 | 1.000 | 0.999 | – | – | – | 0.519 | 1.000 |
| | 0.737 | 0.699 | 0.730 | | | | 0.526 | 0.735 |
| ONT Benchmark – 5mC vs. control | 0.964 | 0.999 | 0.766 | – | 0.994 | 0.860 | 0.603 | 1.000 |
| | 0.624 | 0.515 | 0.467 | | 0.814 | 0.528 | 0.544 | 0.457 |
| ONT Benchmark – 5hmC vs. control | 0.970 | 0.999 | – | 1.000 | – | – | – | 1.000 |
| | 0.647 | 0.405 | | 0.644 | | | | 0.444 |
| *E. coli* Dam vs. Δ*dam*Δ*dcm* | 0.998 | 1.000 | 1.000 | – | – | – | – | 1.000 |
| | 0.857 | 0.584 | 0.640 | | | | | 0.944 |
| *H. pylori* 26695 vs. WGA (GCATG) | 0.998 | 0.988 | 0.990 | – | – | – | – | 0.988 |
| | 0.661 | 0.430 | 0.551 | | | | | 0.651 |
| ONT Benchmark – 6mA vs. control | 0.938 | 1.000 | 0.980 | – | – | – | – | 0.999 |
| | 0.700 | 0.312 | 0.458 | | | | | 0.768 |
| *H. pylori* 26695 vs. WGA (TCTTC) | 0.994 | 0.973 | – | 0.997 | – | – | – | 0.959 |
| | 0.663 | 0.472 | | 0.962 | | | | 0.801 |
| Mean \|control − 0.5\| | 0.209 | 0.113 | 0.092 | 0.303 | 0.403 | 0.257 | 0.033 | 0.198 |
| Rows | 8 | 8 | 6 | 2 | 2 | 2 | 3 | 8 |

#### S9.4. A modification-agnostic UniMeth control

RawMod scores every modification with one shared model. UniMeth-Agn tests whether this ability comes from RawMod's design or only from training one model on every modification. It applies the same training to UniMeth. UniMeth-Agn is fine-tuned from the released UniMeth R10.4.1 5 kHz 5mC model on the training pileups of RawMod, whatever their modification. Each training pileup (library, position, strand and label) labels the reads of that library's benchmark read set at that base: modified at a positive pileup and unmodified at a negative one. A control library carries no modification, so it is labeled unmodified at every training position of its organism, as for UniMeth-FT†. Every other base is masked from the loss. As for UniMeth-FT†, every read that overlaps the held-out region of its organism is removed. UniMeth-Agn, like RawMod, therefore never sees the test regions or their reads (Section S9.7). Training uses all 16 benchmark libraries, 12,000 steps with batch size 32, and UniMeth's class-balanced loss.

UniMeth places a question token after each candidate base of a read and predicts whether that base is modified. UniMeth-Agn places such a token after every C, A and T, so that one model covers every modification. It is run on the same reads and sites as every other model (Table S17). We train two versions:

- one asks UniMeth's context tokens (CpG, CHG and CHH at C, m6A at A, and a new 5hmU token at T);
- one asks a single new [MOD] token at every C, A and T, whose embedding starts as the mean of the four released question tokens.

We report the [MOD] version, which has the higher pooled AUROC (0.972 against 0.963). The two versions differ by at most 0.002 in the mean over the rows and over the modifications, and by 0.017 in the worst modification.

UniMeth-Agn starts from a UniMeth model pretrained on large native datasets. Averaged over the five modification types, it reaches an AUROC of 0.960, against 0.964 for RawMod (Table S17). Where it is ahead on individual rows, the largest leads come from the pairwise rows. On the six rows whose positives and negatives come from one library, it is close to RawMod (mean 0.992 against 0.985). On the oligonucleotide and *H. pylori* rows, it leads by 0.047–0.073 and 0.005–0.013, which are the rows where it also recognizes the library (Section S9.3).

**Table S17: One modification-agnostic UniMeth against RawMod and the UniMeth models of Table S21, in the held-out regions. UniMeth and UniMeth-FT† give the maximum over their models. Mean: over the 15 rows; per mod.: mean of the five per-modification means; harmonic: their harmonic mean; worst: the lowest per-modification mean; pooled: one threshold across all rows (at most 1,000 positives and 1,000 negatives per row; each modification type weighted equally, split equally between its modified and unmodified sites). Parameters are those of each model, counted from the checkpoints we run (Table 1).**

| Model | Mean | Per mod. | Harmonic | Worst | Pooled | Parameters |
|---|---|---|---|---|---|---|
| RawMod (one model) | 0.974 | **0.964** | **0.964** | **0.930** | **0.968** | 0.23M |
| UniMeth (2 released models) | 0.927 | 0.796 | 0.734 | 0.525 | 0.756 | 2×100.3M |
| UniMeth-FT† (3 models) | 0.859 | 0.890 | 0.826 | 0.500 | 0.782 | 3×100.3M |
| UniMeth-Agn (one model) | **0.982** | 0.960 | 0.955 | 0.825 | 0.941 | 100.3M |

#### S9.5. From per-read calls to site scores

The tools report their calls in different formats and coordinates. The third step converts them into one site score with one rule. A small adapter per tool converts the tool's per-read output into one table of (read, reference position, probability):

- Dorado: the probability at a read base is the sum of the modification probabilities (MM/ML tags) over every code that the model emits (for example 5mC + 5hmC for the 5mC_5hmC model), mapped to the reference through the read's alignment. A base without an MM entry counts as unmodified.
- UniMeth: `prob_1` at the reported reference position.
- DeepMod2: the methylation score at the reference position.
- Rockfish: the read-coordinate calls, projected onto the reference through the read's alignment, with a sigmoid where the output is a logit.
- MethyNano: the positive-class probability of each checkpoint.

The aggregation step keeps a call only if its read is one of the 10 selected reads at that site and strand. If a tool reports one read twice at one site, it keeps the maximum. For each site, it records the number of calls, their summed probability and the number of calls above 0.5. A selected read without a call counts as unmodified. The site score is therefore $\bar{p} = \frac{1}{10} \sum_r p_r$, with $p_r$=0 for uncalled reads, and the call frequency is the fraction of the 10 reads with $p_r$>0.5. AUROC is computed on $\bar{p}$. We use this rule so that every tool is scored on the same reads. A model that does not call a row's base scores all of the row's sites 0 and obtains an AUROC of exactly 0.5, which is how Table 1 reports models off their target base. A tool that calls only some reads at a site is penalized only for the reads it leaves out. RawMod is not a per-read caller. For each site and strand, it builds one pileup from exactly the 10 selected reads (reverse-strand pileups in read orientation) and outputs one site score, which we use as $\bar{p}$.

After every aggregation, we check the coordinates by tabulating the reference base under the kept calls. Cytosine models must fall on C for forward reads and on G for reverse reads, adenine models on A and T, the 5hmU model on T and A, and RawMod on the base of each row. On every sample and model, the expected base lies under 82–100% of the kept calls. The range is 87–100% for every tool except MethyNano, and 100% for RawMod, which scores only the row's positions. The remaining calls sit on read bases that do not match the reference. These are most frequent in the SPO1 and oligonucleotide reads, and for MethyNano, whose calls come from its own basecall, also in *H. pylori* 26695. A coordinate error would instead put the calls on the complementary base or spread them over all four bases. Duplicate site keys stop the scoring with an error.

#### S9.6. Ground-truth rows

Each row of the benchmark is one ground-truth set of modified and unmodified sites. The goal of the row design is labels that the reference sequence alone cannot predict. Table S18 lists the 15 rows. The reference base at a position sets the strand of a site: the row's modified base gives the forward strand, its complement the reverse strand, and both strands are scored. No row takes its per-site labels from a modification caller. The rows fall into three kinds.

*Pairwise rows.* A pairwise (two-library) row scores the same positions in a modified and an unmodified sample. The reference sequence therefore cannot predict the label (11-mer null 0.50–0.52).

- The ONT oligonucleotide rows use the 256 modified positions of the 32 constructs (8 per construct) against the same positions in the canonical control.
- The *E. coli* rows use every GATC (6mA, Dam), CCWGG (5mC at the internal C, Dcm) or CG (5mC, M.SssI) on either strand, inside the held-out region (1.74–2.44 Mb). They are scored in the wild-type or M.SssI sample against the Δ*dam*Δ*dcm* knockout.
- The two *H. pylori* 26695 rows score the native sample against its whole-genome amplification at the C of TCTTC (4mC; position 1, both strands) and at the A of GCATG (6mA; position 2, both strands).
- The SPO1 row uses every thymine of the native genome against the PCR amplicons, because 5hmU replaces thymine throughout this genome.

We used Dorado only to decide which motif class a row tests. The dataset's motif annotation also labels the C of GCATG as 4mC. However, Dorado separates native from amplified reads at the adjacent A with its 6mA model (AUROC 0.99) and only weakly at that C with its 4mC model (0.72–0.82). We therefore treat GCATG as a 6mA motif. For the same reason, we do not include an *H. pylori* J99 4mC row, whose annotated 4mC sites Dorado does not call as modified.

*Bisulfite and EM-seq rows.* These rows use the per-strand counts of a bisulfite (HG002) or enzymatic methyl-seq (EM-seq; *A. thaliana*, *O. sativa*, mouse brain) experiment on the same organism: *A. thaliana* chromosome 4 (CpG, and CHG plus CHH as a separate non-CpG row), HG002 (ONT whole-genome bisulfite sequencing of GM24385, Bismark CpG coverage), *O. sativa*, and mouse brain chromosome 1. We use one rule for all of them. A cytosine is assayed if it has at least 20 bisulfite or EM-seq reads on its strand. It is positive at a methylated fraction of at least 0.9, negative at a fraction of at most 0.1, and dropped in between. Only assayed sites of the row's context are scored, so an unassayed cytosine is never a negative.

*Motif row.* The motif row (*Anabaena* GATC for 6mA) has no control sample. Every motif occurrence is positive, and every other adenine in the genome is negative. These labels are a property of the sequence, so the 11-mer null reaches 1.00 on this row. A high AUROC there therefore does not show that a tool models the signal. We keep the row because it is the only 6mA data from this organism, and we mark it in every table. We removed an *H. pylori* J99 4mC motif row ($TCN_7GC$), because Dorado finds no 4mC signal at that motif in this sample.

On every one-library row, we keep all positives and at most 50,000 negatives, chosen by a seeded hash of the site. The same subsample is used for every tool and is the exact set of sites that RawMod featurizes.

**Table S18: Ground-truth rows of the benchmark. Pos., Neg.: scored positive and negative sites inside the held-out regions; Null: AUROC of the reference 11-mer alone on those sites. Controls: KO, Δ*dam*Δ*dcm* knockout; WGA, whole-genome amplification; PCR, PCR amplicons; control, canonical oligonucleotides. Sites are counted after read selection (≥10 reads on the site's strand), the 50,000-negative cap of one-library rows, and restriction to the held-out regions (Section S9.7). The null is the AUROC of a logistic regression on the reference 11-mer with position-grouped five-fold cross-validation on the scored sites; pairwise rows are ≈0.5 by construction. Pairwise rows name their unmodified control library, which Uncalled4 and NovaMod use as their control.**

| Mark | Dataset | Design | Positive | Negative | Pos. | Neg. | Null |
|---|---|---|---|---|---|---|---|
| 5mC CpG | *A. thaliana* chr4 | EM-seq | CpG, fraction ≥0.9 | CpG, fraction ≤0.1 | 3,956 | 20,859 | 0.65 |
| 5mC CpG | HG002 | bisulfite | CpG, fraction ≥0.9 (WGBS) | CpG, fraction ≤0.1 | 640 | 428 | 0.67 |
| 5mC CpG | *O. sativa* chr1:0–13 Mb | EM-seq | CpG, fraction ≥0.9 | CpG, fraction ≤0.1 | 2,667 | 3,522 | 0.59 |
| 5mC CpG | Mouse brain chr1 | EM-seq | CpG, fraction ≥0.9 | CpG, fraction ≤0.1 | 77,301 | 8,012 | 0.68 |
| 5mC CpG | *E. coli* 1.74–2.44 Mb | pairwise (KO) | CG in KO + M.SssI | same CG in KO | 101,197 | 101,125 | 0.49 |
| 5mC non-CpG | *E. coli* 1.74–2.44 Mb | pairwise (KO) | CCWGG internal C in wild type | same C in KO | 3,531 | 3,526 | 0.50 |
| 5mC non-CpG | *A. thaliana* chr4 | EM-seq | CHG/CHH, fraction ≥0.9 | CHG/CHH, fraction ≤0.1 | 87 | 48,571 | 0.93 |
| 5mC | ONT oligonucleotides | pairwise (control) | modified position of each construct | same position in control | 40 | 40 | 0.50 |
| 5hmC | ONT oligonucleotides | pairwise (control) | modified position of each construct | same position in control | 40 | 40 | 0.50 |
| 6mA | *E. coli* 1.74–2.44 Mb | pairwise (KO) | GATC in wild type | same A in KO | 5,597 | 5,581 | 0.50 |
| 6mA | *Anabaena* | motif | GATC | every other A | 2,921 | 50,000 | 1.00 |
| 6mA | ONT oligonucleotides | pairwise (control) | modified position of each construct | same position in control | 40 | 40 | 0.50 |
| 6mA | *H. pylori* 26695 | pairwise (WGA) | GCATG in native | same A after WGA | 846 | 852 | 0.50 |
| 4mC | *H. pylori* 26695 | pairwise (WGA) | TCTTC in native | same C after WGA | 694 | 694 | 0.50 |
| 5hmU | SPO1 | pairwise (PCR) | every T, native genome (reads anchored at no PCR primer) | same T in PCR reads anchored at a primer / product end | 2,701 | 2,701 | 0.50 |

#### S9.7. Test-set design and data leakage

This subsection explains how we keep the test data unseen and what this design cannot control.

A test set can be separated from the training data at four levels:

(i) Positions: no test position is a training position, but test positions can lie next to training positions and come from the same reads.
(ii) Genomic regions: a contiguous part of each genome is removed from training, together with every read that overlaps it, and only sites inside that part are tested. No test read and no test position was seen in training.
(iii) Organisms or libraries: every sample of an organism or library is removed from training (Section S9.8).
(iv) Modifications: every positive of one modification is removed from training (the LOCO folds of Table S1).

Levels (i) and (ii) measure accuracy on data like the training data. Levels (iii) and (iv) measure transfer to new data, and we report them separately. With positions held out alone, the test sites of 9 of the 15 rows would lie 1–10 bp from training positions and come from the same reads. For this reason, no result in this paper uses a position split.

*The benchmark holds out genomic regions.* Table 1 uses level (ii) for both models that we train. The RawMod column is the final configuration retrained without a contiguous ~15% of every genome and without every read that overlaps it (Table S1).

UniMeth-FT$^{\dagger}$ is fine-tuned on the same training positions, with every read that overlaps a held-out region removed from its fine-tuning data. Every model, including the released ones, is then scored only at the benchmark sites inside the held-out regions:

- *E. coli* NC_000913.3:1.74–2.44 Mb;
- *H. pylori* 26695 NC_018939.1:1.08–1.33 Mb;
- SPO1 52.6–72.5 kb;
- five of the 32 oligonucleotide constructs;
- the six *Anabaena* plasmids;
- chromosome 4 of *A. thaliana*;
- chromosome 1 of the human genome.

For *E. coli*, *Anabaena*, *H. pylori* 26695, SPO1 and *A. thaliana*, we draw new test reads: those whose primary alignment starts in the held-out region (read q-score ≥10, at most 15,000 reads per sample; for *A. thaliana*, the held-out region is chromosome 4 and all its 40,548 reads are kept). Every site is still scored from exactly 10 reads per strand. Mouse brain and *O. sativa* were never used in training and keep all their sites.

*What region holdout does not control.* Holding out regions controls what RawMod and UniMeth-FT$^{\dagger}$ have seen. It does not control what the released models were trained on, and several of our test datasets are public (Table S19).

- Rockfish's R10.4.1 model is trained on chromosomes 1–21, X and Y of HG002, with bisulfite labels from the same sample [28]. This training set includes our HG002 test sites on chromosome 1.
- DeepMod2's earlier models are trained on chromosomes 2–21 of HG002, HG003 and HG004 and exclude chromosome 1 [29]. The 5 kHz model we run (v5.0) was released later, and its training data are not documented.
- ONT does not publish the training data of the Dorado modification models. ONT releases the all-5-mer oligonucleotides as validation data [7], so the oligonucleotide rows may be in distribution for Dorado.
- The released UniMeth 5mC model is trained on three other plant species and evaluated on nine plant species, including *A. thaliana* and *O. sativa* [30].
- MethyNano is trained on other *A. thaliana*, *O. sativa* and human (GM12878) libraries [31].

The bacterial, plant and mouse datasets of Kulkarni et al. [8] were released after the Rockfish model, and the SPO1 libraries are unpublished. Any such overlap can only raise the scores of the released callers.

**Table S19: Documented training data of every benchmarked model and its overlap with the benchmark's test sites. "None by construction": trained without the held-out regions and every read overlapping them. Sources: the publications and repositories cited in Section S9.7.**

| Model | Documented training data | Overlap with our test sites |
|---|---|---|
| RawMod | our training samples outside the held-out regions | none by construction |
| UniMeth-FT$^{\dagger}$ (5hmC, 4mC, 5hmU) | UniMeth 5mC / 6mA, fine-tuned at RawMod's training positions outside the held-out regions | none by construction |
| UniMeth-Agn (one model) | UniMeth 5mC, fine-tuned at RawMod's training positions of the 16 libraries that carry them (mouse and *O. sativa* have none), outside the held-out regions | none by construction |
| Dorado 6mA, 4mC+5mC, 5mC+5hmC, 5mCG+5hmCG (v3) | not published; all-context models trained on synthetic DNA | unknown; the oligonucleotide rows may overlap |
| UniMeth 5mC (5 kHz) | *S. miltiorrhiza*, *S. tuberosum*, *R. communis*; pretraining on human data of an unnamed sample | none documented; *A. thaliana* and *O. sativa* used for evaluation only |
| UniMeth 6mA (5 kHz) | HG002 Fiber-seq, chromosome 1 excluded | none documented; *E. coli* used for evaluation only |
| DeepMod2 5mCG (5 kHz) | not documented for v5.0; earlier models: HG002, HG003, HG004, chromosomes 2–21 | unknown; earlier models exclude chromosome 1 |
| Rockfish 5mCG (5 kHz) | HG002, chromosomes 1–21, X, Y; bisulfite labels of the same sample | HG002 row |
| MethyNano 5mC | *A. thaliana*, *O. sativa* (other libraries), GM12878 | other libraries of two of our species |

#### S9.8. In-distribution and leave-out evaluation

The benchmark of Table 1 is in distribution for RawMod. On 13 of the 15 rows, other regions of the same library are in the training data of the RawMod model. Only the held-out regions and their reads are new. UniMeth-FT$^{\dagger}$ is trained on the same data with the same regions held out, so the two are compared on equal terms. The training data of the released tools are documented in Table S19. Here we measure how much of the accuracy of RawMod depends on having seen the library. We score every row with a RawMod model that is trained with the same configuration but without any sample of the row's organism or library. We score it at the same sites inside the held-out regions (Table S20).

Without the library, the mean AUROC drops from 0.974 to 0.825, and the number of rows with AUROC ≥0.9 drops from 15 to 9. CpG methylation in plants and mammals transfers (≥0.97). So do *Anabaena* 6mA (0.979) and *O. sativa* and mouse brain, which are never in training. *E. coli* M.SssI and Dcm 5mC (0.921 and 0.924), *A. thaliana* non-CpG 5mC (0.902) and the 5mC and 5hmC oligonucleotides (0.902 and 0.863) transfer with a loss of at most 0.09. *E. coli* Dam 6mA (0.707), *H. pylori* 26695 6mA and 4mC (0.659 and 0.521), the 6mA oligonucleotides (0.516) and SPO1 5hmU (0.551) do not transfer. For 4mC, 5hmC and 5hmU, the withheld library is the only training source of the modification. Their leave-out scores are therefore zero-shot detection of an unseen modification, as in LOCO, but on a different test set.

The size of the loss depends on how the row is labeled. The loss is largest on the pairwise rows, where only the signal separates the same position in a modified and an unmodified library: *E. coli* (0.07–0.27), the 5hmC and 6mA oligonucleotides (0.09 and 0.41), *H. pylori* 26695 6mA and 4mC (0.34 and 0.46) and SPO1 (0.38). The rows whose positives and negatives come from one library lose at most 0.06. The 5mC oligonucleotides are also pairwise but lose only 0.03.

The per-site scores show two ways in which the leave-out models fail. Each per-site score is the sum of the per-read probabilities over the 10 reads. First, the model trained without *E. coli* scores every GATC adenine as modified, in both the wild type and the Δ*dam*Δ*dcm* knockout (median site score 10.0 and 9.8 of 10). It has learned the GATC motif from the other Dam-methylated bacteria, not the 6mA signal. The same motif solves the *Anabaena* row, whose sequence-only null is 1.00, so this row is not evidence of transfer. Second, on *H. pylori* 26695 6mA and 4mC, the 6mA oligonucleotides and SPO1, the leave-out models score the modified and the unmodified library alike, near zero (median 0.00–0.01 of 10). Without its library, RawMod does not detect these modifications. The cytosine modifications transfer because several libraries contribute modified and unmodified cytosines across sequence contexts. Outside the cytosine modifications, the accuracy of RawMod on the benchmark therefore depends on having seen the library. We list this dependence among the limitations (Section S7.3).

**Table S20: RawMod with genomic regions held out against RawMod with the row's organism or library withheld from training, at the sites of Table 1. Region held out: the RawMod model of Table 1. Leave-out: one model per withheld group, trained with the same configuration. Last column: the best model of Table 1 trained for the row's modification. This reference column is in distribution only for UniMeth-FT† (*H. pylori* 26695 4mC, SPO1), and the training data of the released tools are not public. ¶The withheld library is the only training source of the modification. ‡Sequence alone gives AUROC ≥0.9. Colors as in Table 1.**

| Dataset | Withheld from training | RawMod region held out | RawMod leave-out | Best model trained for the modification |
|---|---|---|---|---|
| **5mC, CpG** | | | | |
| *A. thaliana* | *A. thaliana* | 0.994 | 0.973 | 0.999 |
| HG002 | HG001, HG002 | 0.990 | 0.982 | 0.999 |
| *O. sativa* | none (never in training) | 0.999 | 0.999 | 1.000 |
| Mouse brain | none (never in training) | 0.977 | 0.977 | 1.000 |
| *E. coli* M.SssI vs. Δ*dam*Δ*dcm* | *E. coli* (all samples) | 1.000 | 0.921 | 1.000 |
| **5mC, non-CpG** | | | | |
| *E. coli* Dcm vs. Δ*dam*Δ*dcm* | *E. coli* (all samples) | 0.999 | 0.924 | 1.000 |
| *A. thaliana*‡ | *A. thaliana* | 0.957 | 0.902 | 1.000 |
| **5mC, all contexts** | | | | |
| ONT Benchmark – 5mC vs. control | 5mC oligonucleotides, control | 0.936 | 0.902 | 1.000 |
| **5hmC** | | | | |
| ONT Benchmark – 5hmC vs. control¶ | 5hmC oligonucleotides, control | 0.953 | 0.863 | 1.000 |
| **6mA** | | | | |
| *E. coli* Dam vs. Δ*dam*Δ*dcm* | *E. coli* (all samples) | 0.978 | 0.707 | 1.000 |
| *Anabaena*‡ | *Anabaena* | 0.995 | 0.979 | 0.999 |
| *H. pylori* 26695 vs. WGA (GCATG) | *H. pylori* 26695 (native, WGA) | 0.995 | 0.659 | 1.000 |
| ONT Benchmark – 6mA vs. control | 6mA oligonucleotides, control | 0.927 | 0.516 | 1.000 |
| **4mC** | | | | |
| *H. pylori* 26695 vs. WGA (TCTTC)¶ | *H. pylori* 26695 (native, WGA) | 0.981 | 0.521 | 0.994 |
| **5hmU** | | | | |
| SPO1 native vs. PCR¶ | SPO1 (all barcodes) | 0.930 | 0.551 | 0.965 |
| **Average AUROC** | | 0.974 | 0.825 | |
| Rows with AUROC ≥0.9 | | 15/15 | 9/15 | |

#### S9.9. Evaluation views and summary statistics

The benchmark scores every tool in several views, because one view cannot show both how good each model is on its own modification and how well a tool works when the modification is unknown. This subsection defines the views, gives the notes to Table 1 and Fig. 3, and reports the results that do not fit in the main text.

*Per model* (Table 1). This view shows every model on its own. Each Dorado and UniMeth model is a separate column. Every model is scored on all sites of every row. Bold marks the rows of the modifications that the model is trained for. Table 1 leaves out MethyNano, which Tables S21 and S22 include as the per-site maximum over its three species checkpoints. The footer reports the mean AUROC over the five modifications (the rows of each modification averaged first), the mean over the 15 rows, the worst modification, and the parameters of the model.

*Modification-agnostic* (Table S21). This view is the setting of a user who does not know which modification a sample carries. Each tool gives one score per site, the maximum over all of its models, without being told which modification the row carries. Every tool is scored on all sites of every row. The footer reports the mean AUROC over all rows, the same mean over the rows whose 11-mer null is below 0.75, and a pooled AUROC. For the pooled AUROC, each row is subsampled to at most 1,000 positives and 1,000 negatives (seed 0), all rows are concatenated, and one AUROC is computed with a single threshold. The pooled AUROC is high only if the score of a modified site is comparable across modifications, organisms and sequence contexts.

**Table S21: Modification-agnostic site-level AUROC inside the held-out regions. Each tool gives one score per site, the maximum over all of its models. Every tool is scored on every row, RawMod with its single score. Bottom rows: mean AUROC over all rows, the same over rows whose reference-sequence null is <0.75, the mean, harmonic mean and minimum of the tool's five per-modification AUROCs (rows of each modification averaged first), and a pooled AUROC with one threshold across all rows (each row subsampled to at most 1,000 positives and 1,000 negatives; each modification type weighted equally). UniMeth-FT†: fine-tuned by us (Section S9.3). NovaMod's *E. coli* cells measure the library, not the modification (Section S9.3). Colors, bold and footnotes as in Table 1. Tool abbreviations as in Table 1; DM2: DeepMod2, MN: MethyNano.**

| | *Modification callers* | | | | | | | | *Comparative* | |
|---|---|---|---|---|---|---|---|---|---|---|
| **Dataset** | **RawMod** | **Dor.** | **UM** | **UM-FT†** | **DM2** | **RF** | **MN** | **UM-Agn†** | **Unc4*** | **NM*** |
| **5mC, CpG** | | | | | | | | | | |
| *A. thaliana* | 0.994 | 0.998 | **0.999** | 0.999 | 0.990 | **0.999** | 0.496 | **0.999** | 0.829 | |
| HG002 | 0.990 | 0.987 | **0.999** | 0.993 | 0.997 | **0.999** | 0.687 | 0.997 | 0.903 | |
| *O. sativa*° | 0.999 | 0.991 | **1.000** | 0.999 | 0.999 | **1.000** | 0.517 | **1.000** | 0.894 | |
| Mouse brain° | 0.977 | 0.994 | **1.000** | 0.985 | 0.999 | **1.000** | 0.594 | 0.977 | 0.837 | |
| *E. coli* M.SssI vs. Δ*dam*Δ*dcm* | **1.000** | **1.000** | **1.000** | **1.000** | **1.000** | **1.000** | 0.529 | **1.000** | 0.990 | **1.000** |
| **5mC, non-CpG** | | | | | | | | | | |
| *E. coli* Dcm vs. Δ*dam*Δ*dcm* | 0.999 | **1.000** | 0.990 | 0.955 | 0.479 | 0.479 | 0.498 | **1.000** | 0.960 | **1.000** |
| *A. thaliana*‡ | 0.957 | 0.999 | **1.000** | 0.999 | 0.516 | 0.516 | 0.460 | 0.983 | 0.756 | |
| **5mC, all contexts** | | | | | | | | | | |
| ONT Benchmark – 5mC vs. control | 0.936 | **1.000** | 0.948 | **1.000** | 0.500 | 0.490 | 0.337 | **1.000** | 0.971 | 0.594 |
| **5hmC** | | | | | | | | | | |
| ONT Benchmark – 5hmC vs. control | 0.953 | **1.000** | 0.913 | **1.000** | 0.527 | 0.532 | 0.325 | **1.000** | 0.988 | 0.611 |
| **6mA** | | | | | | | | | | |
| *E. coli* Dam vs. Δ*dam*Δ*dcm* | 0.978 | **1.000** | **1.000** | 0.495 | 0.500 | 0.500 | 0.499 | 0.953 | 0.624 | **1.000** |
| *Anabaena*‡ | 0.995 | 0.999 | **0.999** | 0.497 | 0.499 | 0.499 | 0.492 | 0.996 | 0.409 | |
| *H. pylori* 26695 vs. WGA (GCATG) | 0.995 | 0.995 | 0.994 | 0.497 | 0.500 | 0.500 | 0.511 | **1.000** | 0.961 | 0.535 |
| ONT Benchmark – 6mA vs. control | 0.927 | **1.000** | 0.982 | 0.512 | 0.512 | 0.512 | 0.500 | **1.000** | 0.815 | 0.492 |
| **4mC** | | | | | | | | | | |
| *H. pylori* 26695 vs. WGA (TCTTC) | 0.981 | 0.981 | 0.556 | 0.993 | 0.505 | 0.505 | 0.309 | **0.994** | 0.739 | 0.594 |
| **5hmU** | | | | | | | | | | |
| SPO1 native vs. PCR | 0.930 | 0.521 | 0.525 | **0.965** | 0.500 | 0.500 | 0.514 | 0.825 | 0.745 | 0.517 |
| Mean AUROC, all rows | 0.974 | 0.964 | 0.927 | 0.859 | 0.668 | 0.669 | 0.485 | **0.982** | 0.828 | |
| Mean AUROC, null < 0.75 | 0.974 | 0.959 | 0.916 | 0.876 | 0.693 | 0.694 | 0.486 | **0.980** | 0.866 | |
| Mean over the five modifications | **0.964** | 0.899 | 0.796 | 0.890 | 0.569 | 0.570 | 0.433 | 0.960 | 0.813 | 0.652 |
| Harmonic mean over the five modifications | **0.964** | 0.841 | 0.734 | 0.826 | 0.550 | 0.551 | 0.410 | 0.955 | 0.800 | 0.634 |
| Worst modification | **0.930** | 0.521 | 0.525 | 0.500 | 0.500 | 0.500 | 0.309 | 0.825 | 0.702 | 0.517 |
| Pooled AUROC, one threshold, types weighted equally | **0.968** | 0.818 | 0.756 | 0.782 | 0.534 | 0.537 | 0.466 | 0.941 | 0.806 | |

*Matched* (Table S22). This view gives each tool its best case. Each tool is scored on a row only with the models that it provides for the row's modification and context. If several models qualify, the site score is their maximum, a rule fixed before scoring. For example, Dorado 5mC is the maximum of the 5mCG_5hmCG, 5mC_5hmC and 4mC_5mC models. A CpG-only model on a row of mixed context is scored on that row's CpG sites only. A tool without a model for the modification is N/A.

*ROC curves.* The per-row curves (Fig. S8) use the matched rule. Each curve is computed on the full site set. We draw it on a logarithmic false-positive-rate axis, where callers with AUROC above 0.99 still differ. The pooled curves of Fig. 3a use the modification-agnostic score on the same subsampled sites as the pooled AUROC.

*Notes to Table 1.* The SPO1 row is scored on the restricted read sets of Section S9.1, which keep only reads whose library of origin the alignment establishes. RawMod, UniMeth-FT† and UniMeth-Agn† are trained without the held-out regions and without every read that overlaps them (Section S9.7). The results are still in distribution for RawMod at the level of the library. Except for the rows marked °, every row comes from a library whose other regions are in its training data (RawMod with the row's organism or library withheld: Table S20). A model with no call at the row's base scores 0.5. Only NovaMod is blank on the rows without an unmodified control library, and its summaries are over the rows it scores. Rows labeled "vs." compare a modified library with its matched unmodified library, which the comparative methods use as their control. Cells are colored by

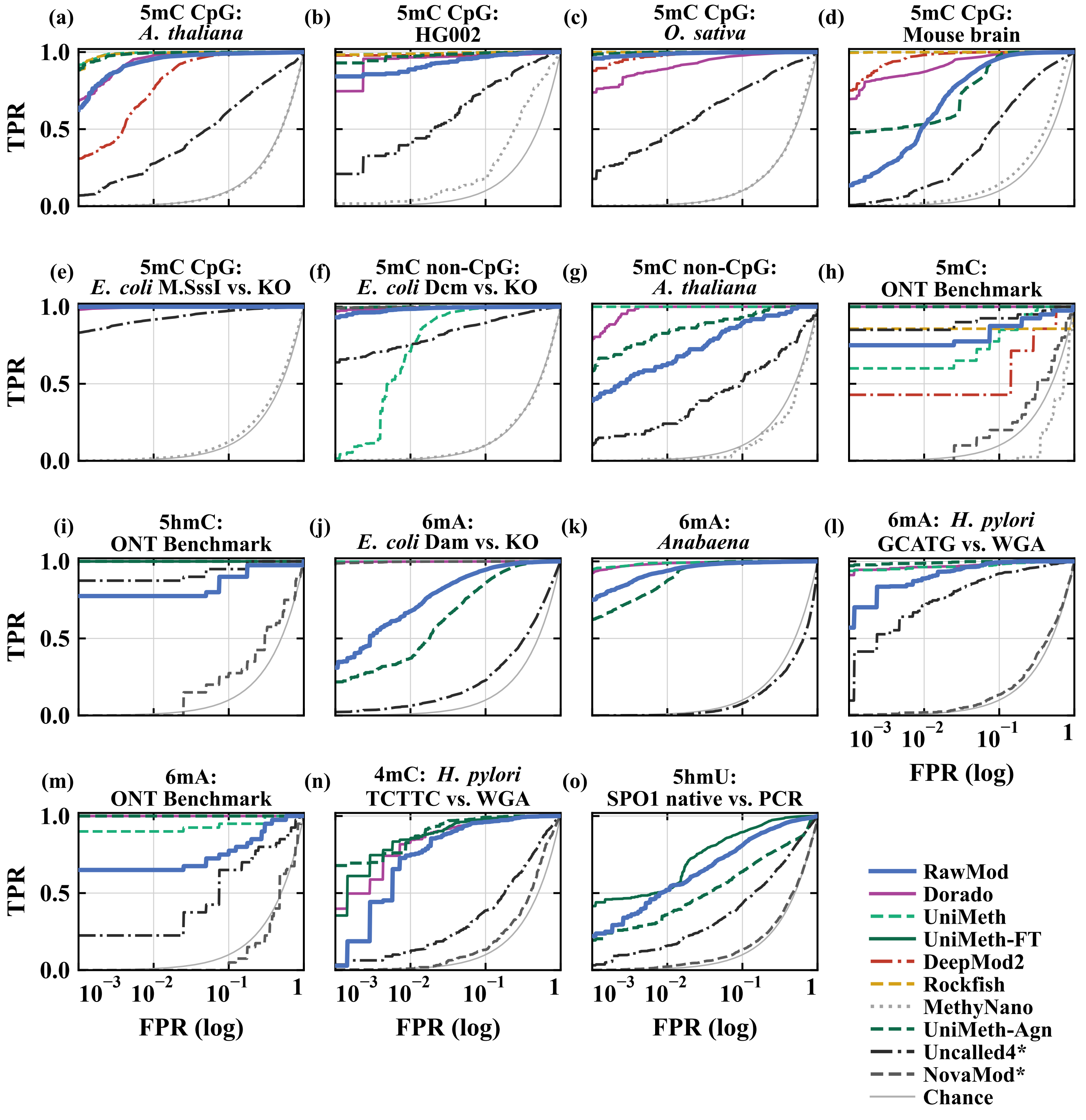


**Figure S8:** Per-row ROC curves inside the held-out regions under the matched rule. The false-positive-rate axis is logarithmic; legends give each AUROC. RawMod blue, Dorado purple, UniMeth aqua (dashed), UniMeth-FT[†] dark aqua, DeepMod2 red (dash-dot), Rockfish gold (dashed), MethyNano light grey (dotted), Uncalled4[*] near-black (dash-dot), NovaMod[*] grey (dashed). [c]CpG sites only. Each tool is drawn only on the rows of a modification it has a model for; its AUROCs are in Table S22. On the ONT 5mC row, DeepMod2 and Rockfish are scored on the CpG sites only.

**Table S22: Matched site-level AUROC inside the held-out regions. Each tool is scored only with its models for the row's modification (per-site maximum if several). N/A: no such model. $^c$CpG-only model scored at CpG sites. Bottom rows: mean AUROC over the rows a tool scores, the number of such rows, the number of the five modifications the tool covers, and the number of models it needs to cover them. Footnotes as in Table 1. Tool abbreviations as in Table 1; DM2: DeepMod2, MN: MethyNano.**

| | *Modification callers* | | | | | | | | *Comparative* | |
|---|---|---|---|---|---|---|---|---|---|---|
| **Dataset** | **RawMod** | **Dor.** | **UM** | **UM-FT**† | **DM2** | **RF** | **MN** | **UM-Agn**† | **Unc4*** | **NM*** |
| **5mC, CpG** | | | | | | | | | | |
| *A. thaliana* | 0.994 | 0.998 | **0.999** | N/A | 0.990 | **0.999** | 0.496 | **0.999** | 0.829 | |
| HG002 | 0.990 | 0.987 | **0.999** | N/A | 0.997 | **0.999** | 0.687 | 0.997 | 0.903 | |
| *O. sativa*◦ | 0.999 | 0.991 | **1.000** | N/A | 0.999 | **1.000** | 0.517 | **1.000** | 0.894 | |
| Mouse brain◦ | 0.977 | 0.994 | **1.000** | N/A | 0.999 | **1.000** | 0.594 | 0.977 | 0.837 | |
| *E. coli* M.SssI vs. Δ*dam*Δ*dcm* | **1.000** | **1.000** | **1.000** | N/A | **1.000** | **1.000** | 0.529 | **1.000** | 0.990 | **1.000** |
| **5mC, non-CpG** | | | | | | | | | | |
| *E. coli* Dcm vs. Δ*dam*Δ*dcm* | 0.999 | **1.000** | 0.990 | N/A | N/A | N/A | 0.498 | **1.000** | 0.960 | **1.000** |
| *A. thaliana*‡ | 0.957 | 0.999 | **1.000** | N/A | N/A | N/A | 0.460 | 0.983 | 0.756 | |
| **5mC, all contexts** | | | | | | | | | | |
| ONT Benchmark – 5mC vs. control | 0.936 | **1.000** | 0.949 | N/A | 0.837$^c$ | 0.857$^c$ | 0.337 | **1.000** | 0.971 | 0.594 |
| **5hmC** | | | | | | | | | | |
| ONT Benchmark – 5hmC vs. control | 0.953 | **1.000** | N/A | **1.000** | N/A | N/A | N/A | **1.000** | 0.988 | 0.611 |
| **6mA** | | | | | | | | | | |
| *E. coli* Dam vs. Δ*dam*Δ*dcm* | 0.978 | **1.000** | **1.000** | N/A | N/A | N/A | N/A | 0.953 | 0.624 | **1.000** |
| *Anabaena*‡ | 0.995 | 0.999 | **0.999** | N/A | N/A | N/A | N/A | 0.996 | 0.409 | |
| *H. pylori* 26695 vs. WGA (GCATG) | 0.995 | 0.995 | 0.994 | N/A | N/A | N/A | N/A | **1.000** | 0.961 | 0.535 |
| ONT Benchmark – 6mA vs. control | 0.927 | **1.000** | 0.982 | N/A | N/A | N/A | N/A | **1.000** | 0.815 | 0.492 |
| **4mC** | | | | | | | | | | |
| *H. pylori* 26695 vs. WGA (TCTTC) | 0.981 | 0.984 | N/A | 0.988 | N/A | N/A | N/A | **0.994** | 0.739 | 0.594 |
| **5hmU** | | | | | | | | | | |
| SPO1 native vs. PCR | 0.930 | N/A | N/A | **0.965** | N/A | N/A | N/A | 0.825 | 0.745 | 0.517 |
| **Average AUROC** | 0.974 | **0.996** | 0.993 | 0.984 | 0.970 | 0.976 | 0.515 | 0.982 | 0.828 | 0.705 |
| Rows scored | 15/15 | 14/15 | 12/15 | 3/15 | 6/15 | 6/15 | 8/15 | 15/15 | 15/15 | 9/15 |
| Modifications | 5/5 | 4/5 | 2/5 | 3/5 | 1/5 | 1/5 | 1/5 | 5/5 | 5/5 | 5/5 |
| Models | 1 | 4 | 2 | 3 | 1 | 1 | 3 | 1 | 1 | 1 |

AUROC from red (≤0.5) through yellow (0.75) to green (1.0). Bold in the bottom rows marks the best model. UniMeth-FT† and UniMeth-Agn† are the models we fine-tune on RawMod's training positions (Sections S9.3 and S9.4). Uncalled4* and NovaMod* use the row's unmodified control library, including its reads in the held-out regions (Section S9.3); NovaMod's *E. coli* cells measure the library it was trained on rather than the modification (Section S9.3). Parameters are the learned weights of each modification model, counted from the checkpoints we run. We count no basecaller and no k-mer table of expected current. Uncalled4 has no learned model, because it compares the signal with a 9-mer pore model.

*Notes to Fig. 3.* In (a) and (b), each tool gives one score per site without being told the modification: the maximum over all of its models, and the single score for RawMod. (a) Tools that cover more than one modification. Each row contributes at most 1,000 positives and 1,000 negatives, each modification type is weighted equally, and legends give the number of models and the pooled AUROC. (b) Logarithmic parameter axis. The number after a tool is its number of models. Dotted line: chance. (c) Background positions as in Section S9.3, scored with the models of the tool that claim the row's modification. The AUROC between the native and the control library is shown as the invariance $1 - 2\,|\text{AUROC} - 0.5|$, so that a tool that separates the libraries in either direction scores low. Small dots are rows and large dots the mean over the rows the tool covers. Models above the line were trained on these libraries, and models below it are released. Uncalled4* and NovaMod* compare a sample with its control library by design, so they are not in (b) or (c). Uncalled4 is in (a). NovaMod has no score on the rows without a control library, so it is in Table 1 only. Per-row ROC curves: Fig. S8.

*Per-model results.* RawMod reaches an AUROC of at least 0.9 on every one of the 15 rows; the best released model, Dorado 4mC+5mC, does so on 10. Off their base, the other models score close to chance (0.46–0.52), and within a base the Dorado cytosine models are not specific to the mark: the 4mC+5mC model reaches 1.000 on the 5hmC oligonucleotides. On their own rows, Dorado and UniMeth remain more accurate than RawMod (mean above 0.99), most on mouse brain, which RawMod never trained on (0.977 against ≥0.993), and on the ONT oligonucleotides, which keep only 40 positive and 40 negative sites each inside the held-out constructs (5mC 0.936, 95% CI 0.868–0.986; 5hmC 0.953, 0.897–0.993; 6mA 0.927, 0.871–0.967).

Figure S8 shows that these gaps are largest at low false-positive rates. At 1% false positives, RawMod detects 52% of the modified mouse CpGs and 68% of the *E. coli* Dam sites, against ≥99.9% for UniMeth. At 10% false positives, it detects 96% and 95%. At 1% false positives, it detects more *E. coli* Dcm sites than UniMeth (99% against 71%). MethyNano stays near chance even on 5mC (0.515, Table S22), for the reason given in Section S9.3.

When each tool combines all of its models into one score (Table S21), RawMod keeps a higher mean AUROC over the 15 rows than the four models of Dorado (0.974 against 0.964). It is also higher on the rows whose sequence-only null is below 0.75 (0.974 against 0.959). It leads by more on the harmonic mean over the five modifications (0.964 against 0.841).

*One threshold in operation* (Table S23). The pooled AUROC measures how well a score ranks sites across rows, but a user must also fix one threshold. Table S23 therefore fixes the threshold. We threshold the modification-agnostic score of each tool (the maximum over its models) once for all rows. The threshold is the value that gives 1% false positives on the pooled negatives of the agnostic table (RawMod 0.993, Dorado 0.760, UniMeth-Agn$^{\dagger}$ 0.927). We also report a threshold of 0.5. At the shared 1% threshold, RawMod recovers a mean of 68% of the modified sites over the 15 rows. This ranges from 18% (ONT 6mA, 40 positives) to 99% (*E. coli* M.SssI). The three oligonucleotide rows and *A. thaliana* non-CpG stay below 40%. The row mean weights 5mC by 8 of the 15 rows. Averaged over the five modification types instead, RawMod recovers 60%, Dorado 57% and UniMeth-Agn$^{\dagger}$ 57%. Over the 15 rows, the four models of Dorado recover 79% on average. They recover 16% on 4mC, 0% on 5hmU, for which no Dorado model exists, and 62–100% elsewhere. UniMeth-Agn$^{\dagger}$ recovers 61%. At 0.5, the false-positive rate of RawMod is below 1% on 7 rows. It reaches 17% on HG002 and 37% on SPO1, whose PCR negatives include native carryover (Section S9.2). The pooled lead of RawMod therefore comes from scoring every modification on one scale, not from a sharper operating point on each row.

**Table S23: Operating points at one shared threshold inside the held-out regions. TPR at the threshold giving 1% FPR on the pooled negatives (last row), and TPR and FPR at 0.5. Modification-agnostic score per tool.**

| | TPR at the shared 1% FPR threshold | | | TPR at 0.5 | | | FPR at 0.5 | | |
|---|---|---|---|---|---|---|---|---|---|
| **Row** | RawMod | Dorado | UniMeth-Agn$^{\dagger}$ | RawMod | Dorado | UniMeth-Agn$^{\dagger}$ | RawMod | Dorado | UniMeth-Agn$^{\dagger}$ |
| *A. thaliana* CpG | 0.77 | 0.89 | 0.39 | 0.92 | 0.99 | 0.99 | 0.01 | 0.02 | 0.00 |
| HG002 | 0.93 | 0.97 | 0.82 | 0.98 | 0.98 | 0.99 | 0.17 | 0.28 | 0.07 |
| *O. sativa* | 0.89 | 0.97 | 0.78 | 0.98 | 0.99 | 0.99 | 0.00 | 0.36 | 0.00 |
| Mouse brain | 0.87 | 0.97 | 0.54 | 0.96 | 1.00 | 1.00 | 0.10 | 0.33 | 0.12 |
| *E. coli* M.SssI | 0.99 | 0.99 | 0.88 | 1.00 | 1.00 | 1.00 | 0.00 | 0.08 | 0.00 |
| *E. coli* Dcm | 0.80 | 0.99 | 0.55 | 0.96 | 1.00 | 0.99 | 0.00 | 0.18 | 0.00 |
| *A. thaliana* non-CpG | 0.34 | 0.76 | 0.07 | 0.61 | 1.00 | 0.67 | 0.01 | 0.01 | 0.00 |
| ONT 5mC | 0.40 | 1.00 | 0.82 | 0.80 | 1.00 | 1.00 | 0.07 | 0.00 | 0.00 |
| ONT 5hmC | 0.33 | 0.95 | 0.80 | 0.80 | 1.00 | 1.00 | 0.07 | 0.00 | 0.00 |
| *E. coli* Dam | 0.70 | 0.91 | 0.40 | 0.96 | 0.99 | 0.98 | 0.14 | 0.00 | 0.31 |
| *Anabaena* | 0.86 | 0.62 | 0.87 | 0.98 | 0.89 | 1.00 | 0.04 | 0.00 | 0.10 |
| *H. pylori* 6mA | 0.89 | 0.68 | 0.78 | 1.00 | 0.90 | 1.00 | 0.14 | 0.00 | 0.03 |
| ONT 6mA | 0.17 | 0.95 | 0.70 | 0.85 | 1.00 | 0.97 | 0.25 | 0.00 | 0.00 |
| *H. pylori* 4mC | 0.66 | 0.16 | 0.21 | 0.95 | 0.60 | 0.91 | 0.06 | 0.01 | 0.02 |
| SPO1 5hmU | 0.58 | 0.00 | 0.56 | 0.95 | 0.00 | 1.00 | 0.37 | 0.00 | 0.98 |
| **Mean over rows** | 0.68 | 0.79 | 0.61 | 0.91 | 0.89 | 0.97 | 0.10 | 0.08 | 0.11 |
| Mean over the five types | 0.60 | 0.57 | 0.57 | 0.91 | 0.71 | 0.97 | 0.14 | 0.03 | 0.23 |
| Worst type | 0.33 | 0.00 | 0.21 | 0.80 | 0.00 | 0.91 | 0.37 | 0.16 | 0.98 |
| Threshold (1% FPR) | 0.993 | 0.760 | 0.927 | | | | | | |

*Different modifications on the same sequence* (Table S24). The pooled AUROC mixes organisms and sequence contexts. This test removes that difference: it scores different modifications on the same sequence under one threshold. Set (A) is one library. Wild-type *E. coli* carries 6mA at GATC and 5mC at CCWGG, scored against the $dam^{-}dcm^{-}$ knockout at the same positions inside the held-out region (18,235 sites). Set (B) adds the M.SssI library (5mC at CpG) and its knockout (220,557 sites). Set (C) pools the ONT oligonucleotide rows against the control library (240 sites). Their constructs carry 5mC, 5hmC or 6mA at designed positions. Each tool is scored on the same 10 reads per site as in Table 1. We report the AUROC over the pooled set, which uses one threshold for every modification in it. We also report the share of modified sites recovered at the threshold that gives 1% false positives on the pooled unmodified sites (TPR). A single released model detects one modification and is at chance on the other, so its pooled AUROC falls to 0.49–0.73. RawMod reaches 0.987, 1.000 and 0.939 with one model and one threshold. Taking the maximum over the models of a tool recovers the pooled AUROC (Dorado 0.999, 1.000 and 1.000), because 5mC and 6mA each have a released model. The sets contain no 5hmU or 4mC, where no released model exists.

*Confidence intervals* (Table S25). For RawMod against UniMeth-Agn$^{\dagger}$, a paired version of the same bootstrap (1,000 resamples shared by both tools) gives a difference in the mean over the five modifications of +0.004 (95% CI −0.009 to +0.013) and in the weakest modification of +0.105 (+0.069 to +0.117). Table S25 repeats Table 1 with a 95% confidence interval under each AUROC and the number of scored sites. We compute each interval with a cluster bootstrap over reference positions (400 resamples). A row whose two classes are two libraries scores the same reference position twice. Resampling sites on their own would therefore understate the interval, so we resample positions and keep every site of a resampled position. The intervals are within 0.005 of the estimate on all but two of the genome-wide rows. They are wide on the three oligonucleotide rows, which keep 40 modified and 40 unmodified positions each inside the held-out region: 0.868–0.986 for 5mC, 0.897–0.993 for 5hmC and 0.871–0.967 for

**Table S24: Different modifications on the same sequence under one threshold. Pooled AUROC and TPR at 1% FPR per set; see the text for the three sets.**

| | (A) *E. coli* WT library 6mA (GATC) + 5mC (CCWGG) | | (B) *E. coli* genome + M.SssI 5mC (CpG) | | (C) ONT oligonucleotides 5mC + 5hmC + 6mA | |
|---|---|---|---|---|---|---|
| **Model** | AUROC | TPR | AUROC | TPR | AUROC | TPR |
| RawMod | 0.987 | 0.79 | 1.000 | 1.00 | 0.939 | 0.67 |
| Dorado 6mA | 0.694 | 0.61 | 0.499 | 0.05 | 0.563 | 0.33 |
| Dorado 4mC+5mC | 0.575 | 0.39 | 0.950 | 0.94 | 0.730 | 0.67 |
| Dorado 5mC+5hmC | 0.576 | 0.39 | 0.952 | 0.95 | 0.734 | 0.67 |
| Dorado 5mCG+5hmCG | 0.492 | 0.01 | 0.921 | 0.92 | 0.519 | 0.12 |
| Dorado, maximum over its four models | 0.999 | 0.98 | 1.000 | 1.00 | 1.000 | 1.00 |
| UniMeth 5mC | 0.573 | 0.36 | 0.950 | 0.95 | 0.694 | 0.29 |
| UniMeth 6mA | 0.690 | 0.61 | 0.497 | 0.05 | 0.560 | 0.30 |
| UniMeth, maximum over its two models | 0.997 | 0.94 | 1.000 | 1.00 | 0.945 | 0.55 |
| UniMeth-FT†, maximum over three models | 0.565 | 0.22 | 0.950 | 0.94 | 0.729 | 0.67 |
| UniMeth-Agn† | 0.976 | 0.53 | 1.000 | 1.00 | 1.000 | 1.00 |
| Rockfish | 0.492 | 0.01 | 0.921 | 0.92 | 0.508 | 0.08 |
| DeepMod2 | 0.492 | 0.01 | 0.921 | 0.92 | 0.511 | 0.04 |

6mA. The 5hmC row is the only dataset of its modification, so the 5hmC per-modification mean of Table 1 carries the width of a 40-position estimate. Differences between tools smaller than these widths should not be read as differences in accuracy.

**Table S25: Table 1 with 95% confidence intervals. (a) RawMod with the released Dorado and UniMeth models. (b) RawMod with the fine-tuned and comparative models and Rockfish. Tool abbreviations as in Table 1.**

| *(a)* **Dataset** | **Sites** | **RawMod** | **Dor. 6mA** | **Dor. 4mC+5mC** | **Dor. 5(h)mC** | **Dor. 5(h)mCG** | **UM 5mC** | **UM 6mA** |
|---|---|---|---|---|---|---|---|---|
| **5mC, CpG** | | | | | | | | |
| *A. thaliana* | 24,815 | 0.994 [0.99,1.00] | 0.502 [0.50,0.50] | 0.998 [1.00,1.00] | 0.999 [1.00,1.00] | 0.999 [1.00,1.00] | 0.999 [1.00,1.00] | 0.503 [0.50,0.51] |
| HG002 | 1,068 | 0.990 [0.99,0.99] | 0.459 [0.44,0.47] | 0.986 [0.98,0.99] | 0.994 [0.99,1.00] | 0.997 [0.99,1.00] | 0.999 [1.00,1.00] | 0.458 [0.44,0.47] |
| *O. sativa* | 6,189 | 0.999 [1.00,1.00] | 0.500 [0.50,0.50] | 0.989 [0.99,0.99] | 0.997 [1.00,1.00] | 0.998 [1.00,1.00] | 1.000 [1.00,1.00] | 0.504 [0.50,0.51] |
| Mouse brain | 85,313 | 0.977 [0.98,0.98] | 0.504 [0.50,0.51] | 0.993 [0.99,0.99] | 0.999 [1.00,1.00] | 1.000 [1.00,1.00] | 1.000 [1.00,1.00] | 0.508 [0.51,0.51] |
| *E. coli* M.SssI vs. Δ*dam*Δ*dcm* | 202,322 | 1.000 [1.00,1.00] | 0.495 [0.50,0.50] | 0.999 [1.00,1.00] | 1.000 [1.00,1.00] | 1.000 [1.00,1.00] | 1.000 [1.00,1.00] | 0.495 [0.49,0.50] |
| **5mC, non-CpG** | | | | | | | | |
| *E. coli* Dcm vs. Δ*dam*Δ*dcm* | 7,057 | 0.999 [1.00,1.00] | 0.503 [0.50,0.50] | 1.000 [1.00,1.00] | 1.000 [1.00,1.00] | 0.479 [0.47,0.48] | 0.990 [0.99,0.99] | 0.504 [0.50,0.51] |
| *A. thaliana* | 48,658 | 0.957 [0.93,0.98] | 0.495 [0.49,0.50] | 0.999 [1.00,1.00] | 0.999 [1.00,1.00] | 0.516 [0.50,0.54] | 1.000 [1.00,1.00] | 0.501 [0.49,0.52] |
| **5mC, all contexts** | | | | | | | | |
| ONT Benchmark – 5mC vs. control | 80 | 0.936 [0.87,0.99] | 0.500 [0.46,0.54] | 1.000 [1.00,1.00] | 1.000 [1.00,1.00] | 0.515 [0.48,0.55] | 0.949 [0.91,0.98] | 0.512 [0.47,0.55] |
| **5hmC** | | | | | | | | |
| ONT Benchmark – 5hmC vs. control | 80 | 0.953 [0.90,0.99] | 0.500 [0.47,0.54] | 1.000 [1.00,1.00] | 1.000 [1.00,1.00] | 0.539 [0.51,0.58] | 0.916 [0.86,0.97] | 0.500 [0.47,0.54] |
| **6mA** | | | | | | | | |
| *E. coli* Dam vs. Δ*dam*Δ*dcm* | 11,178 | 0.978 [0.98,0.98] | 1.000 [1.00,1.00] | 0.499 [0.50,0.50] | 0.499 [0.50,0.50] | 0.500 [0.50,0.50] | 0.499 [0.50,0.50] | 1.000 [1.00,1.00] |
| *Anabaena* | 52,921 | 0.995 [0.99,1.00] | 0.999 [1.00,1.00] | 0.498 [0.50,0.50] | 0.499 [0.50,0.50] | 0.499 [0.50,0.50] | 0.498 [0.50,0.50] | 0.999 [1.00,1.00] |
| *H. pylori* 26695 vs. WGA (GCATG) | 1,698 | 0.995 [0.99,1.00] | 0.995 [0.99,1.00] | 0.501 [0.50,0.51] | 0.501 [0.50,0.51] | 0.500 [0.50,0.50] | 0.501 [0.50,0.51] | 0.994 [0.99,1.00] |
| ONT Benchmark – 6mA vs. control | 80 | 0.927 [0.87,0.97] | 1.000 [1.00,1.00] | 0.512 [0.50,0.54] | 0.512 [0.50,0.54] | 0.512 [0.50,0.54] | 0.512 [0.50,0.54] | 0.983 [0.95,1.00] |
| **4mC** | | | | | | | | |
| *H. pylori* 26695 vs. WGA (TCTTC) | 1,388 | 0.981 [0.97,0.99] | 0.700 [0.68,0.72] | 0.984 [0.98,0.99] | 0.866 [0.85,0.88] | 0.505 [0.50,0.51] | 0.521 [0.50,0.54] | 0.779 [0.76,0.80] |
| **5hmU** | | | | | | | | |
| SPO1 native vs. PCR | 5,402 | 0.930 [0.92,0.94] | 0.506 [0.50,0.51] | 0.515 [0.51,0.52] | 0.508 [0.50,0.51] | 0.500 [0.50,0.50] | 0.516 [0.51,0.52] | 0.511 [0.51,0.51] |

#### S9.10. Reproducibility

The release contains everything needed to rerun this benchmark. The benchmark code is in `test/` in `https://github.com/STORMgroup/RawMod`. It contains one download script per dataset with the public links we used, the fixed read sets, the ground-truth positions and the RawMod training positions. It also contains the scripts that run every tool and regenerate every table and figure of this section. Samples, tools, claims, rows, bisulfite sources and motifs are tab-separated configuration files. The location of each tool is set once in a site file. Every derived file carries a tag of its inputs and scoring parameters, so that results

**Table S25: (continued) Table 1 with 95% confidence intervals, panel (b).**

| *(b)* **Dataset** | **RawMod** | **UM-FT$^\dagger$ 5hmC** | **UM-FT$^\dagger$ 4mC** | **UM-FT$^\dagger$ 5hmU** | **RF 5mCG** | **UM-Agn$^\dagger$ all** | **Unc4$^*$ control** | **NM$^*$ control** |
|---|---|---|---|---|---|---|---|---|
| **5mC, CpG** | | | | | | | | |
| *A. thaliana* | 0.994 [0.99,1.00] | 0.998 [1.00,1.00] | 0.999 [1.00,1.00] | 0.507 [0.50,0.51] | 0.999 [1.00,1.00] | 0.999 [1.00,1.00] | 0.829 [0.82,0.84] | – |
| HG002 | 0.990 [0.99,0.99] | 0.995 [0.99,1.00] | 0.997 [1.00,1.00] | 0.375 [0.35,0.39] | 0.999 [1.00,1.00] | 0.998 [1.00,1.00] | 0.903 [0.89,0.92] | – |
| *O. sativa* | 0.999 [1.00,1.00] | 0.995 [0.99,1.00] | 1.000 [1.00,1.00] | 0.505 [0.50,0.51] | 1.000 [1.00,1.00] | 1.000 [1.00,1.00] | 0.894 [0.89,0.90] | – |
| Mouse brain | 0.977 [0.98,0.98] | 0.998 [1.00,1.00] | 0.998 [1.00,1.00] | 0.435 [0.43,0.44] | 1.000 [1.00,1.00] | 0.977 [0.97,0.98] | 0.837 [0.83,0.84] | – |
| *E. coli* M.SssI vs. Δ*dam*Δ*dcm* | 1.000 [1.00,1.00] | 1.000 [1.00,1.00] | 1.000 [1.00,1.00] | 0.498 [0.50,0.50] | 1.000 [1.00,1.00] | 1.000 [1.00,1.00] | 0.990 [0.99,0.99] | 1.000 [1.00,1.00] |
| **5mC, non-CpG** | | | | | | | | |
| *E. coli* Dcm vs. Δ*dam*Δ*dcm* | 0.999 [1.00,1.00] | 0.979 [0.98,0.98] | 0.942 [0.94,0.95] | 0.500 [0.50,0.50] | 0.479 [0.47,0.48] | 1.000 [1.00,1.00] | 0.960 [0.96,0.96] | 1.000 [1.00,1.00] |
| *A. thaliana* | 0.957 [0.93,0.98] | 0.994 [0.99,1.00] | 0.998 [1.00,1.00] | 0.497 [0.49,0.52] | 0.516 [0.50,0.54] | 0.983 [0.97,0.99] | 0.756 [0.69,0.82] | – |
| **5mC, all contexts** | | | | | | | | |
| ONT Benchmark – 5mC vs. control | 0.936 [0.87,0.99] | 1.000 [1.00,1.00] | 0.946 [0.90,0.98] | 0.513 [0.47,0.55] | 0.490 [0.46,0.53] | 1.000 [1.00,1.00] | 0.971 [0.93,1.00] | 0.594 [0.52,0.67] |
| **5hmC** | | | | | | | | |
| ONT Benchmark – 5hmC vs. control | 0.953 [0.90,0.99] | 1.000 [1.00,1.00] | 0.924 [0.87,0.97] | 0.500 [0.46,0.54] | 0.532 [0.50,0.57] | 1.000 [1.00,1.00] | 0.988 [0.97,1.00] | 0.611 [0.51,0.71] |
| **6mA** | | | | | | | | |
| *E. coli* Dam vs. Δ*dam*Δ*dcm* | 0.978 [0.98,0.98] | 0.499 [0.50,0.50] | 0.499 [0.50,0.50] | 0.496 [0.49,0.50] | 0.500 [0.50,0.50] | 0.953 [0.95,0.96] | 0.624 [0.61,0.63] | 1.000 [1.00,1.00] |
| *Anabaena* | 0.995 [0.99,1.00] | 0.498 [0.50,0.50] | 0.498 [0.50,0.50] | 0.499 [0.50,0.50] | 0.499 [0.50,0.50] | 0.996 [1.00,1.00] | 0.409 [0.40,0.42] | – |
| *H. pylori* 26695 vs. WGA (GCATG) | 0.995 [0.99,1.00] | 0.501 [0.50,0.51] | 0.501 [0.50,0.51] | 0.495 [0.49,0.50] | 0.500 [0.50,0.50] | 1.000 [1.00,1.00] | 0.961 [0.95,0.97] | 0.535 [0.51,0.55] |
| ONT Benchmark – 6mA vs. control | 0.927 [0.87,0.97] | 0.512 [0.50,0.54] | 0.512 [0.50,0.54] | 0.500 [0.46,0.54] | 0.512 [0.50,0.54] | 1.000 [1.00,1.00] | 0.815 [0.71,0.90] | 0.492 [0.38,0.59] |
| **4mC** | | | | | | | | |
| *H. pylori* 26695 vs. WGA (TCTTC) | 0.981 [0.97,0.99] | 0.813 [0.79,0.83] | 0.988 [0.98,0.99] | 0.716 [0.69,0.73] | 0.505 [0.50,0.51] | 0.994 [0.99,1.00] | 0.739 [0.71,0.76] | 0.595 [0.58,0.61] |
| **5hmU** | | | | | | | | |
| SPO1 native vs. PCR | 0.930 [0.92,0.94] | 0.516 [0.51,0.52] | 0.516 [0.51,0.52] | 0.965 [0.96,0.97] | 0.500 [0.50,0.50] | 0.825 [0.81,0.83] | 0.745 [0.73,0.76] | 0.517 [0.51,0.52] |

from different settings are never mixed. Here the tag is `own1_k10_seed0_q10`: our own basecalls, $K$=10, seed 0 and minimum read quality 10. One option selects the held-out evaluation and leaves the per-site files unchanged: `--holdout region` keeps the sites inside the held-out regions (`test/data/holdout/regions`) and scores the region models (Table 1). From the per-site files, one command regenerates all tables and figures (`rawmod_bench.py score`), and one regenerates the leave-out table (`rawmod_bench.py leaveout`).

#### S10. Computational resources

Table S26 reports wall time, CPU time, and peak memory for every stage of a genome-scale scan with RawMod. Measurements come from the *Amphidinium carterae* contig_3762 run, whose input was every read mapping to the contig from two PromethION flow cells consisting of 4.05 GB of POD5 and 28,870 reads. Every stage is parallel over reads or genome chunks, so costs scale linearly with sequencing yield and scored sites.

##### S10.1. Basecalling and refinement

Dorado basecalls each flow-cell subset in 9 min on one RTX A5000 and aligns reads to the reference in the same pass, emitting move tables. Refining the moves against the 9-mer level table is CPU-only, with each of 16 shards finishing in 2–3 min on four cores with at most 2.6 GB of RAM.

##### S10.2. Featurization

RawMod builds the pileup of each site directly from the refined reads and scores it immediately, so no input is written to disk. The genome is split into eight chunks of similar read coverage, each handled by one process. Each process streams the reads of its chunk, resamples each read's refined segments to $L$=10 samples per base, assembles the pileup of every site for up to $N$=15 overlapping reads, computes the reference row and deviation channels online, and passes the pileups to the model in batches.

##### S10.3. Inference

On contig_3762, the eight processes build and score all 2,170,929 forward-strand T sites in 4.8 min on one RTX A5000 with 8 CPU cores (about 7,500 sites per second with 0.57 CPU-hours), and in 37.7 min without a GPU (4.46 CPU-hours). Together they peak at 57 GB of RAM (Table S26).

**Table S26: Time and memory costs per stage of RawMod. Evaluated on the *Amphidinium carterae* contig_3762 scan of 7.73 Mb and 2.17 M scored sites.**

| Stage | Hardware (per job) | Jobs | Wall / job | CPU-h (total) | Peak RAM |
|---|---|---|---|---|---|
| Basecalling + alignment (Dorado) | 1×RTX A5000, 16 CPU | 2 | 8.9–9.6 min | 0.74 | 10.0 GB |
| Signal refinement | 4 CPU | 16 | 2.0–3.2 min | 0.60 | 2.6 GB |
| merge | 8 CPU | 1 | 1.0 min | 0.01 | 9.1 GB |
| Featurization and scoring, fused (GPU) | 1×RTX A5000, 8 CPU | 1 | 4.8 min | 0.57 | 57.3 GB |
| Featurization and scoring, fused (CPU only) | 8 CPU | 1 | 37.7 min | 4.46 | 57.6 GB |
| Training (final `mixed` model) | 1×RTX A5000, 16 CPU | 1 | 2.5 h | 14.7 | 5.8 GB host |

#### S10.4. Computational cost comparison

We compare the costs of every caller on the same 13,110 reads of *H. pylori* 26695 (1.67 Mb) on one node with 8 CPU cores, one NVIDIA RTX A5000 GPU, and 64 GB of RAM (Table S27). Every caller starts with Dorado basecall with move tables (3.2 min), so we report each caller's cost following this step. We record wall time, GPU utilization, GPU energy, CPU time, peak host memory over all processes, and peak GPU memory. Each Dorado modification model runs the basecaller again together with the modification model instead of reusing the basecall. Its four models take four passes in 12.5 min (12.2 GPU-min). The 6mA model can share a pass with the 4mC+5mC model, but the three cytosine models cannot share one, so three passes are required (9.9 min, 9.7 GPU-min). UniMeth's two released models take 37.9 min, almost all of it on the GPU. RawMod refines the signal segmentation, then builds one pileup per position and strand, and scores each pileup as it is built, finishing in 5.4 min. Of this time, the GPU is busy for 0.9 min, 11× to 14× less than Dorado, with one model that covers every modification. The cost instead moves to the CPU: RawMod uses 0.59 CPU-hours against 0.35–0.45 for Dorado, and its eight processes together peak at 17.4 GB of RAM against 4.5 GB. In a separate run on the same GPU, scoring after refinement takes 8.7 min with one process at a peak of 6.3 GB of RAM and 3.4 GB of GPU memory. It takes 5.9 min and 8.2 GB of RAM with two processes and 4.1 min and 12.1 GB with four. A single RawMod process therefore still uses more RAM than Dorado, but less than a sixth of its GPU memory. With the eight parallel processes of Table S27, it uses as much GPU memory as Dorado (23.5 against 23.3 GB). Without a GPU, the same map takes RawMod 24.5 min and 2.9 CPU-hours, about twice the wall time of Dorado's four passes. Its CPU scores differ from its GPU scores by at most 0.01 at every one of the 1.05 million sites.

**Table S27: Cost of every caller after the shared basecalling step. Measured on 13,110 *H. pylori* 26695 reads (1.67 Mb) and one node (8 CPU cores, one RTX A5000, 64 GB RAM). GPU busy is the GPU utilization integrated over time. GPU energy includes the idle draw of the GPU, so it is not zero without GPU use. Peak RAM is summed over all processes of a step.**

| Tool | Covers | Wall (min) | GPU busy (min) | GPU energy (kJ) | CPU (h) | Peak RAM (GB) | GPU mem. (GB) |
|---|---|---|---|---|---|---|---|
| Shared basecall | – | 3.2 | 2.6 | 39 | 0.06 | 3.3 | 22.4 |
| Dorado, 4 models, 4 passes | 6mA, 4mC, 5(h)mC | 12.5 | 12.2 | 168 | 0.45 | 4.4 | 23.4 |
| Dorado, 4 models, 3 passes | 6mA, 4mC, 5(h)mC | 9.9 | 9.7 | 133 | 0.35 | 4.5 | 23.3 |
| DeepMod2 | 5mCpG | 1.3 | 0.2 | 8 | 0.09 | 9.1 | 7.1 |
| Rockfish | 5mCpG | 3.9 | 3.3 | 48 | 0.09 | 2.2 | 15.3 |
| UniMeth, 2 models | 5mC, 6mA | 37.9 | 35.7 | 509 | 0.82 | 3.1 | 23.5 |
| RawMod, GPU | any | 5.4 | 0.9 | 28 | 0.59 | 17.4 | 23.5 |
| RawMod, CPU only | any | 24.5 | 0.0 | 36 | 2.91 | 27.6 | 0.0 |

#### S10.5. Basecaller and boundary refinement comparison

RawMod depends on a basecaller and on the refinement of each read's segment boundaries, and both add cost. We therefore vary each on the benchmark read sets and score the same model (Table S28).

To vary the basecaller, we re-basecall the read sets with the Dorado `hac` and `fast` models (v5.0.0, move tables emitted, aligned to the reference) in place of `sup`. On every genomic dataset the AUROC stays within 0.01 of the `sup` result under both. Only the three oligonucleotide datasets lose accuracy, where the `hac` AUROC falls to 0.67–0.83 and most reads do not map at all with `fast`. Basecalling the *H. pylori* read set on one RTX A5000 takes 102 s with `sup`, 37 s with `hac` and 23 s with `fast`. On the *E. coli* read set it takes 227, 49 and 28 s.

To vary refinement, we skip the Remora refining of segment boundaries against the k-mer level table. The refinement moves half of the boundaries by more than 5 signal samples and a fifth by more than 10. Without refinement the mean AUROC falls from 0.974 to 0.725 with `sup` basecalls and to near chance with `hac` (0.557) and `fast` (0.476) basecalls on every dataset. The refinement is CPU-only and takes 2–3 min per shard of a flow cell (Table S26).

**Table S28: Site-level AUROC of the region-holdout RawMod model inside the held-out regions. All columns use the same reads, basecalled with one of three Dorado models. Left: segment boundaries refined by Remora against the k-mer level table. Right: boundaries taken from the move table and the alignment, without refinement. Where some sites receive no read, the AUROC over the sites that do follows in parentheses. Mean over modifications first averages the datasets of each modification, as in Table 1.**

| | | | refined boundaries | | | unrefined boundaries | | |
|---|---|---|---|---|---|---|---|---|
| **Dataset** | $n_+$ | $n_-$ | **sup** | **hac** | **fast** | **sup** | **hac** | **fast** |
| *E. coli* Dam 6mA | 5,597 | 5,581 | 0.978 | 0.974 | 0.975 | 0.709 | 0.567 | 0.562 |
| *E. coli* Dcm 5mC | 3,531 | 3,526 | 0.999 | 0.996 | 0.996 | 0.775 | 0.520 | 0.537 |
| *E. coli* M.SssI 5mC | 101,197 | 101,125 | 1.000 | 1.000 | 1.000 | 0.850 | 0.516 | 0.500 |
| *Anabaena* 6mA | 2,921 | 50,000 | 0.995 | 0.994 | 0.995 | 0.798 | 0.672 | 0.694 |
| *H. pylori* 4mC | 694 | 694 | 0.981 | 0.984 | 0.985 | 0.629 | 0.622 | 0.588 |
| *H. pylori* 6mA | 846 | 852 | 0.995 | 0.995 | 0.995 | 0.687 | 0.664 | 0.612 |
| *A. thaliana* CpG 5mC | 3,956 | 20,859 | 0.994 | 0.993 | 0.994 | 0.802 | 0.524 | 0.535 |
| *A. thaliana* non-CpG 5mC | 87 | 48,571 | 0.957 | 0.949 | 0.950 | 0.758 | 0.500 | 0.537 |
| HG002 CpG 5mC | 640 | 428 | 0.990 | 0.990 | 0.988 | 0.893 | 0.767 | 0.741 |
| *O. sativa* CpG 5mC | 2,667 | 3,522 | 0.999 | 0.999 | 0.999 | 0.729 | 0.660 | 0.613 |
| Mouse CpG 5mC | 77,301 | 8,012 | 0.977 | 0.974 | 0.975 | 0.750 | 0.606 | 0.567 |
| ONT oligos 5mC | 40 | 40 | 0.936 | 0.829 | 0.246 (0.616) | 0.667 | 0.414 | 0.139 (0.348) |
| ONT oligos 5hmC | 40 | 40 | 0.953 | 0.782 | 0.014 (0.550) | 0.712 | 0.515 | 0.001 (0.025) |
| ONT oligos 6mA | 40 | 40 | 0.927 | 0.666 (0.833) | 0.126 (0.628) | 0.746 | 0.383 (0.478) | 0.079 (0.395) |
| SPO1 5hmU | 2,701 | 2,701 | 0.930 | 0.918 | 0.926 | 0.366 | 0.421 | 0.435 |
| *mean over datasets* | | | 0.974 | 0.936 | 0.811 | 0.725 | 0.557 | 0.476 |
| *mean over modifications* | | | 0.964 | 0.911 | 0.718 | 0.644 | 0.538 | 0.406 |
| *mean over datasets, scored sites only* | | | 0.974 | 0.947 | 0.905 | 0.725 | 0.563 | 0.513 |
